# Biosynthesis of Macrocyclic Peptides by Formation and Crosslinking of *ortho*-Tyrosines

**DOI:** 10.1101/2025.04.04.647296

**Authors:** Chandrashekhar Padhi, Lingyang Zhu, Jeff Y. Chen, Ryan Moreira, Wilfred A. van der Donk

## Abstract

Ribosomally synthesized and posttranslationally modified peptides (RiPPs) are a growing class of natural products that possess many activities that are of potential translational interest. Multinuclear non-heme iron dependent oxidative enzymes (MNIOs), until recently termed domain of unknown function 692 (DUF692), have been gaining interest because of their involvement in a range of biochemical reactions that are remarkable from a chemical perspective. Over 13,500 putative MNIO-encoding biosynthetic gene clusters (BGCs) have been identified by sequence similarity networks (SSNs). In this study, we identified a set of precursor peptides containing a conserved FHAFRF-motif in MNIO-encoding BGCs. These BGCs follow a conserved synteny with genes encoding an MNIO, a RiPP recognition element (RRE)-containing partner protein, an arginase, and a B12-dependent radical SAM enzyme (rSAM). Using heterologous reconstitution of a representative BGC from *Peribacillus simplex* (*pbs* cluster) in *E. coli*, we demonstrated that the MNIO in conjunction with the partner protein catalyzes *ortho*-hydroxylation of each of the phenylalanine residues in the conserved FRF-motif, the arginase forms an ornithine by deguanidination of the arginine in the motif, and the B12-rSAM crosslinks the *ortho*-Tyr side side chains by a C-C linkage forming a novel macrocyclic molecule. Substrate scope studies suggested tolerance of the MNIO and the B12-rSAM towards substituting the Phe residues with tyrosines in the conserved motif with the position of hydroxylation and crosslinking being maintained. Overall, this study expands the diverse array of posttranslational modifications catalyzed by MNIOs and B12-rSAM enzymes.

**TOC Graphic:** 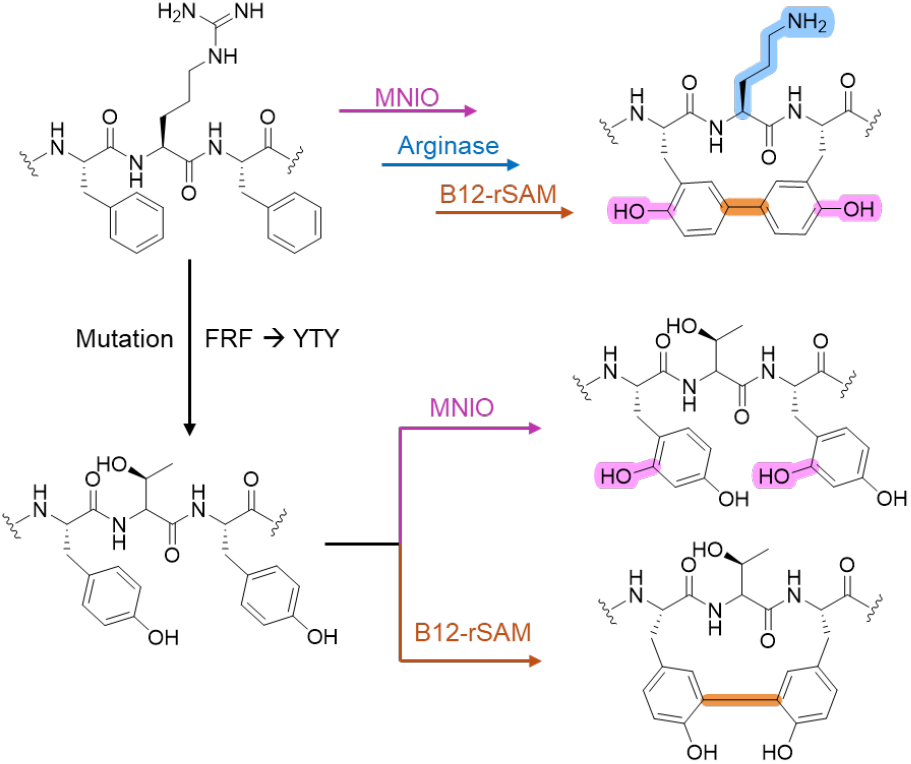

## Introduction

The use of biocatalysis in industry has witnessed a remarkable rise with most pharmaceutical companies having programs to use enzymes in especially the production phase of drug development.^1–4^ Enzymes decrease step-count of synthetic routes, provide high stereoselectivities, and result in more cost-effective and greener manufacturing.^5–9^ However, a key limitation is the repertoire of available enzymes. Whereas a process chemist benefits from a century of synthetic chemistry methods to devise various potential routes to a drug candidate, retrosynthetic biosynthesis^10^ is at present much more limited. Directed evolution has proven remarkably successful in improving low enzyme activities to commercially useful levels,^11^ but if a given transformation has no precedent in enzyme catalyzed reactions, the process of finding a starting point for such evolution is challenging. One source of new enzymes is the microbial genomes that encode a myriad of proteins with unknown function, especially those involved in natural product biosynthesis,^12^ but assignment of their catalytic power is difficult because their substrates are unknown and often not readily accessible.

Natural products (NPs) and NP-derived molecules exhibit tremendous chemical and structural diversity and serve as a major source of modern therapeutics^13–15^ and novel enzymes.^12^ One class of NPs is the family of ribosomally synthesized and posttranslationally modified peptides (RiPPs)^16^ that contain intriguing chemical functionalities that confer various biomedically relevant activities such as antibiotic, antiviral and protease inhibitory activities.^17–19^ Enzymes that introduce these functionalities are typically encoded by genes near the gene encoding the RiPP precursor peptide(s) in biosynthetic gene clusters (BGCs). This proximity of the substrate gene provides a key advantage for assigning function to proteins of unknown function.^20^ Within each RiPP family, the types of modifications are numerous and include methylation, epimerization, peptide skeleton rearrangements, and macrocyclization.^16^ Classical approaches in NP discovery typically employ chemical extraction and rely on producer cultivation and NP production. However, recent advancements in meta(genomic) sequencing techniques have resulted in the detection of novel RiPP BGCs in genomes of uncultivated bacteria, thus expanding the chemical space of NPs.^21,22^ Since RiPPs are ribosomally produced and modified by enzymes, transplanting these genomic elements from an uncultivated organism into a heterologous laboratory strain has greatly accelerated the discovery of novel RiPP families,^16^ and thereby their biosynthetic enzymes.^20^

A recently characterized family of RiPP modification enzymes are the multinuclear non-heme iron-dependent oxidative enzymes (MNIOs), formerly known as domain of unknown function 692 (DUF692). MNIOs catalyze diverse posttranslational chemistries^23^ (Figure 1) such as oxazolone and thioamide formation in methanobactins,^24^ carbon excision in thiaglutamate,^25^ generation of thiooxazoles in bufferins,^26^ formation of two rings in chryseobasins,^27^ C-terminal amidation,^28^ and the transformation of an Asp residue into a C-terminal α-keto acid in pyruvatides.^29^ Given the highly diverse reactions, at present, prediction of the outcome of MNIO-catalyzed transformations on precursor peptides is not possible for MNIOs that are part of groups that have not been previously investigated.

**Figure 1.**
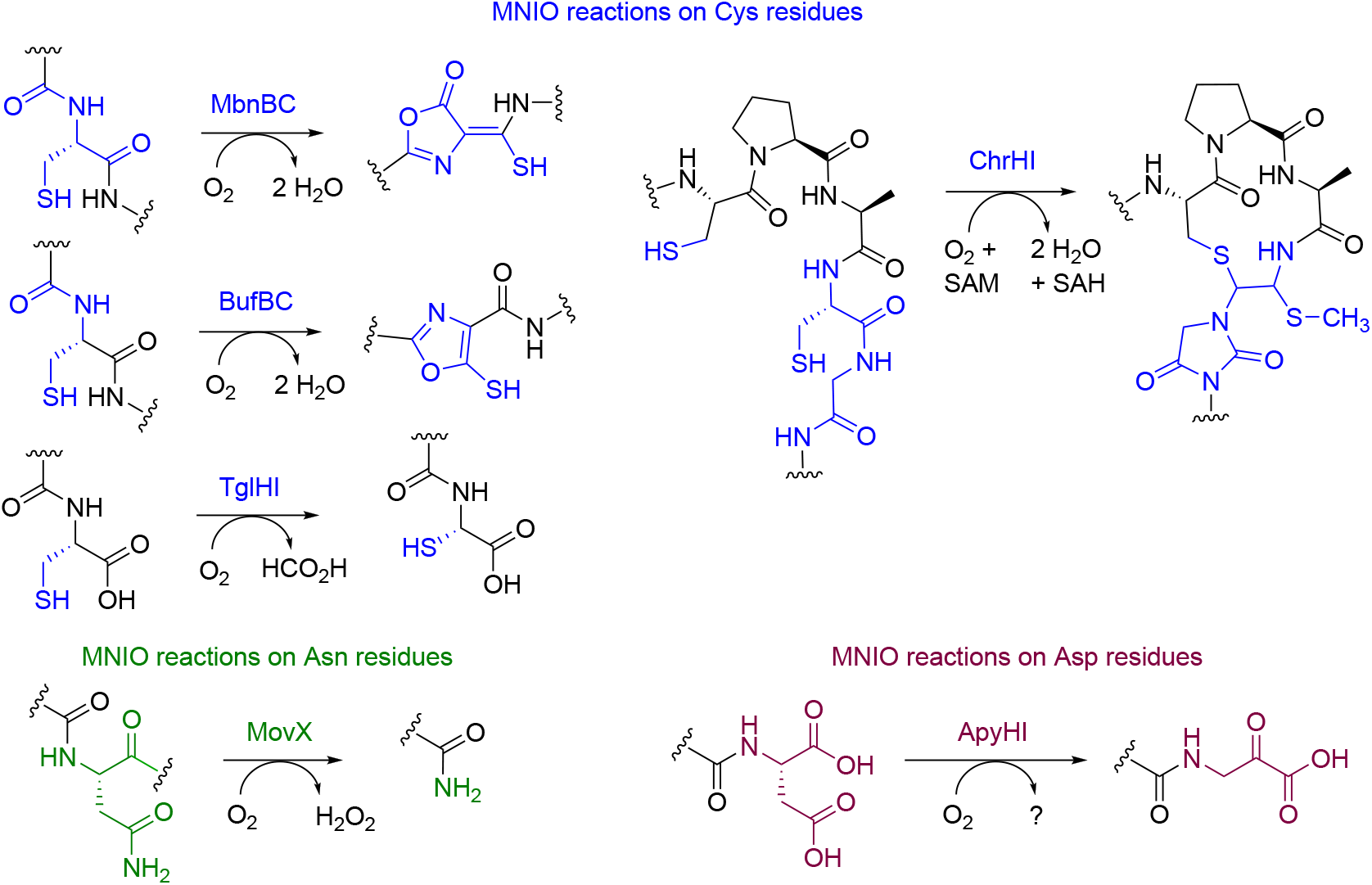
MNIO catalyzed reactions. The majority of known MNIO reactions utilize Cys residues as substrate (in blue). Only two non-cysteine-based reactions acting on Asn (green) and Asp (maroon) residues have been reported to date.

In this study, we generated a sequence similarity network (SSN) for over 13,500 MNIOs using the Enzyme Function Initiative (EFI) tools,^30,31^ and extracted the harboring BGCs using the RODEO program.^32^ We then analyzed the precursor sequences to identify putative RiPP precursor peptides sharing a conserved motif. Based on this approach, putative precursors with conserved FHAFRF-, FHTFMF-, and YHx^1^Yx^2^Y-motifs (x^1^ = S/T/A; x^2^ = T/V/A) were identified. The corresponding BGCs contain genes encoding an MNIO, a hypothetical partner protein, and a B12-dependent rSAM. In certain cases, multiple B12-rSAM-encoding genes were detected, while an arginase-encoding element was only observed in BGCs containing the FHAFRF-motif precursor. Using pathway reconstitution in *Escherichia coli* as a heterologous host, the modified products of the FHAFRF motif-containing precursor from *Peribacillus simplex* BE23 (*pbs* cluster) were structurally characterized using a combination of biophysical techniques. The MNIO (PbsC) partners with a RiPP recognition element-containing^33^ protein (PbsD) to introduce *ortho-*hydroxylations on the two aromatic residues of the FRF-part of the conserved motif, followed by the deguanidination of the Arg to Orn catalyzed by the arginase (PbsE), a transformation that took place exclusively on the bis-hydroxylated precursor peptide. Subsequently, the B12-rSAM (PbsB) crosslinked the hydroxylated aromatic residues via a C-C linkage. In addition to the studies in *E. coli*, the activity of the MNIO and the arginase enzymes of the *pbs* cluster were reconstituted in vitro. We assessed the substrate tolerance of the MNIO PbsC and the B12-rSAM towards a series of FHAFRF-variants including motifs containing tyrosines at the modification sites identified in orthologous BGCs. Finally, utilizing AlphaFold models, we predict the interactions of the pathway enzymes with the precursor peptides. To our knowledge, an MNIO catalyzing hydroxylation of aromatic residues, an arginase selectively acting on a bis-hydroxylated aromatic substrate and a B12-rSAM introducing C-C crosslinks on hydroxylated aromatic side chains, have not been described before. Overall, this study characterizes the activity of three different metalloenzymes in the biosynthesis of a novel RiPP.

## Results

### Genome mining of MNIO-containing BGCs using a sequence similarity network

MNIOs belong to the protein family (Pfam) PF05114 and constitute approximately 13,500 entries in the Uniprot database as of September 2024. Utilizing the Enzyme Function Initiative Enzyme Similarity Tool (EFI-EST) version 2024_04/101,^31^ a sequence similarity network was created with default parameters and processed with an alignment score of 55 (Figure 2A). This analysis resulted in 73 clusters with at least 3 interconnected nodes. Cluster coloring was conducted using the EFI-EST’s Color SSNs functionality and the resultant representative node (rep-node) 50, where sequences over 50% identity threshold were grouped into a meta-node was visualized in Cytoscape v3.10. Subsequently, the Uniprot IDs of the nodes from individual clusters were extracted and subjected to RODEO analysis^32^ using default parameters to obtain the genome neighborhood of the selected MNIOs. Open reading frames (ORFs) ranging from a length of 30 to 150 amino acids were extracted from the RODEO output files (a combination of multiple putative clusters and standalone ORFs) and subjected to multiple sequence alignment with free end gaps at a similarity cost matrix of 65%, gap open penalty of 12, and gap extension penalty of 3 using the Geneious tool. The alignment was refined with two iterations. Subsequently, aligned precursors containing a conserved motif were extracted manually and curated based on their genome neighborhood.

**Figure 2.**
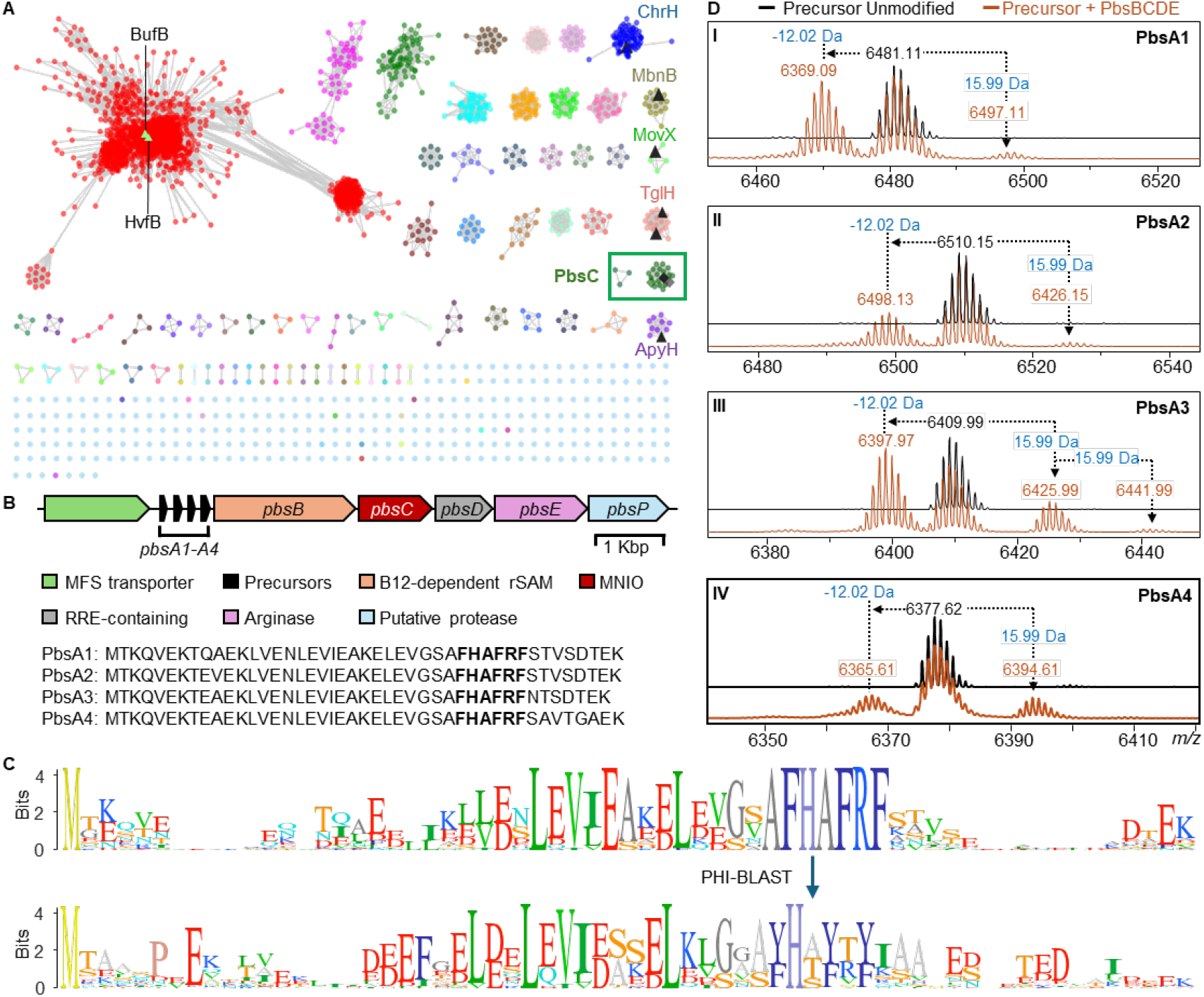
Genome mining of MNIO-containing BGCs encoding precursors with conserved motifs. (A) SSN of ca. 13,500 MNIO homologues was generated with the EFI tools using the UNIREF90 database. RepNode 50 is shown as visualized in cytoscape v3.10. Previously characterized MNIOs are depicted in black triangles. The lime-colored triangles represent the nodes for HvfB and BufB that remain underneath the other nodes in the red cluster. The cluster boxed in green contains the MNIO PbsC that is the focus of this study. (B) BGC architecture of the *pbs* cluster containing the genes encoding four precursor peptides (PbsA1-A4), a B12-dependent radical SAM enzyme (PbsB), an MNIO (PbsC), a putative RiPP recognition element-containing partner protein (PbsD), and an arginase (PbsE). (C) Sequence logo of precursor peptides containing the conserved FHAFRF motif, extracted from orthologous BGCs in the SSN (dark green nodes enclosed in box in panel A). (D) MALDI-TOF mass spectra of N-terminally 6xHis-tagged PbsA1-A4 peptides expressed alone (black trace) or co-expressed with PbsB, PbsC, PbsD and PbsE in *E. coli* (orange trace); sub-panels assigned to Roman numerals I-IV for PbsA1-A4, respectively. For sequences of the translated full length precursors, see Table S7.

### Identification of BGCs based on conserved motifs in the RiPP precursor sequence

Based on the aforementioned pipeline, a set of precursors containing a highly conserved FHAFRF motif was identified in over 30 potential BGCs (Figure 2C, S1). The majority of these BGCs were found in genomes of members of the phyla Actinomycetota and Bacillota. These BGCs contained genes encoding a putative MNIO, a potential partner protein for the MNIO, a putative vitamin B12-dependent radical *S*-adenosylmethionine enzyme (B12-rSAM), and a tetratricopeptide repeat (TPR)-containing putative arginase enzyme (Figure 2B, S2A). We then used the conserved FHAFRF motif as a query parameter for Pattern Hit Initiated BLAST (PHI-BLAST)^34^ analysis for 12 iterations at an inclusion threshold of 0.01 to further identify 99 hits containing similar conserved motifs i.e. FHTFMF and YHx_1_Yx_2_Y (x_1_ = S/T/A; x_2_ = T/V/A) in orthologous clusters (Figure S1). Deeper analysis of the BGCs containing the FHAFRF-, YHx_1_Yx_2_Y- and FHTFMF-motif harboring peptides suggested that the associated MNIOs of all 99 hits are part of the same cluster in the MNIO SSN (Figure 2A; green box), suggesting they may represent isofunctional MNIOs. All orthologous clusters encoded at least one B12-dependent radical SAM (B12-rSAM) enzyme with slight variations in the remaining BGC elements (Figure S2A). For BGCs containing precursor peptides with the FHTFMF and YHx_1_Yx_2_Y motifs, the arginase was absent suggesting it may act on the Arg residue in the FHAFRF motif (Figure S2A). Conversely, a second B12-dependent rSAM and a DUF5825-containing protein was encoded adjacent to the core enzyme genes in BGCs containing the YHxYxY precursors. A recent study has characterized the function of a homologous B12-rSAM and a DUF5825 pair in the biosynthesis of clavusporins,^35^ showing *C-*methylations of amino acids.

### Selection of a BGC for enzyme function elucidation

We selected a BGC (*pbs* cluster) from the genome of a recently reported organism, *Peribacillus simplex* BE23 isolated from the rhizospere of maize plants in France (GenBank: RRZF00000000.1).^36^ Previous studies have reported that *P. simplex* strains produce siderophores that help iron uptake in potatoes, and produce bioactive molecules for inhibition of plant pathogens such as various fungi, bacteria and nematodes.^37–39^ While some antimicrobial activity has been linked to certain lipopeptides^38^ and volatile organic compounds,^40^ structural characterization of the bioactive molecules remains largely understudied. The *pbs* cluster (Figure 2B) encodes four putative precursor peptides containing an FHAFRF motif (*pbsA1*-*A4*), followed by ORFs encoding a putative B12-dependent rSAM enzyme (*pbsB*), a xylose isomerase-like TIM barrel family protein with a typical signature of an MNIO (*pbsC*; ENA ID: RRN73831.1; RefSeq: WP_125160327), a hypothetical ORF suspected to be a RiPP recognition element (RRE)-containing partner protein^41^ for the MNIO (*pbsD*) and a TPR-rich family protein belonging to the (α/β)-hydrolase superfamily believed to be an arginase (*pbsE*). The *pbs* cluster also contained ORFs for a transporter of the Major Facilitator Superfamily (MFS) and a putative metalloprotease (Figure S2A).

### Characterization of the posttranslational modifications of *pbs* peptides

For pathway reconstitution, we co-expressed different genetic elements heterologously in *E. coli*. Codon-optimized genes encoding the precursor peptides PbsA1-A4 were cloned into a pETDuet-1 plasmid backbone encoding a N-terminal 6xHis tag. Codon-optimized ORFs for PbsB, PbsC, PbsD and PbsE were cloned into a pRSFDuet vector in a polycistronic manner, separated by ribosomal binding sequences (RBS). Respective constructs were expressed in *E. coli* BL21 (DE3) TUNER cells, followed by immobilized metal affinity chromatography (IMAC) to purify the peptides.

First, we confirmed the production of the unmodified precursors, PbsA1-A4, by matrix-assisted laser desorption/ionization time-of-flight mass spectrometry (MALDI-TOF MS) analysis (Figure 2D; spectra in black). Second, we co-expressed PbsA1-A4 in combination with PbsB, PbsC, PbsD and PbsE (PbsBCDE) wherein, all four precursor peptides underwent identical mass shifts, showing mass gains of 15.99 Da and 31.99 Da and a mass loss of −12.02 Da with respect to the unmodified precursor mass (Figure 2D; spectra in brown). We next focused on just one precursor (PbsA3) for subsequent experiments because of its higher solubility.

For co-expression of PbsA3 with the individual enzymes, we cloned *pbsB, pbsC, pbsD, pbsBE* (polycistronic), and *pbsCD* (polycistronic) into pRSFDuet vectors. Similarly, *pbsE* was cloned into pCDFDuet. Next, PbsA3 was co-expressed with individual or different combinations of the pathway enzymes. PbsA3 did not undergo any modification (determined by a shift in mass) when co-expressed with PbsB, PbsC, PbsD or PbsE individually in *E. coli* (Figure 3A ii-v). However, when co-expressed with PbsCD, PbsA3 underwent mass gains of 15.99 Da and 31.99 Da suggesting that PbsC and PbsD acted in partnership (Figure 3A vi). PbsA3, co-expressed with PbsBCD did not undergo any further modification suggesting that the PbsCD-modified products did not act as substrates for PbsB (Figure 3A vii). Co-expression of PbsA3 and PbsCDE resulted in a mass loss of 42 Da relative to the PbsCD-modified +31.99 Da-product leading to a net mass loss of 10 Da from the unmodified peptide (Figure 3A viii). Therefore, PbsE requires modification of PbsA3 by PbsCD for activity. We then co-expressed PbsA3 with PbsBCDE with supplementation with B^12^ (hydroxocobalamin) and use of a plasmid encoding the iron-sulfur cluster assembly proteins and the BtuCEDFB proteins for B^12^ uptake (pIGB240).^42–45^ This experiment resulted in a product that underwent a further mass loss of 2 Da relative to the PbsCDE product (Figure 3A ix, Figure S3). Collectively, these data show that PbsB requires prior modification by PbsCD and PbsE for activity.

**Figure 3.**
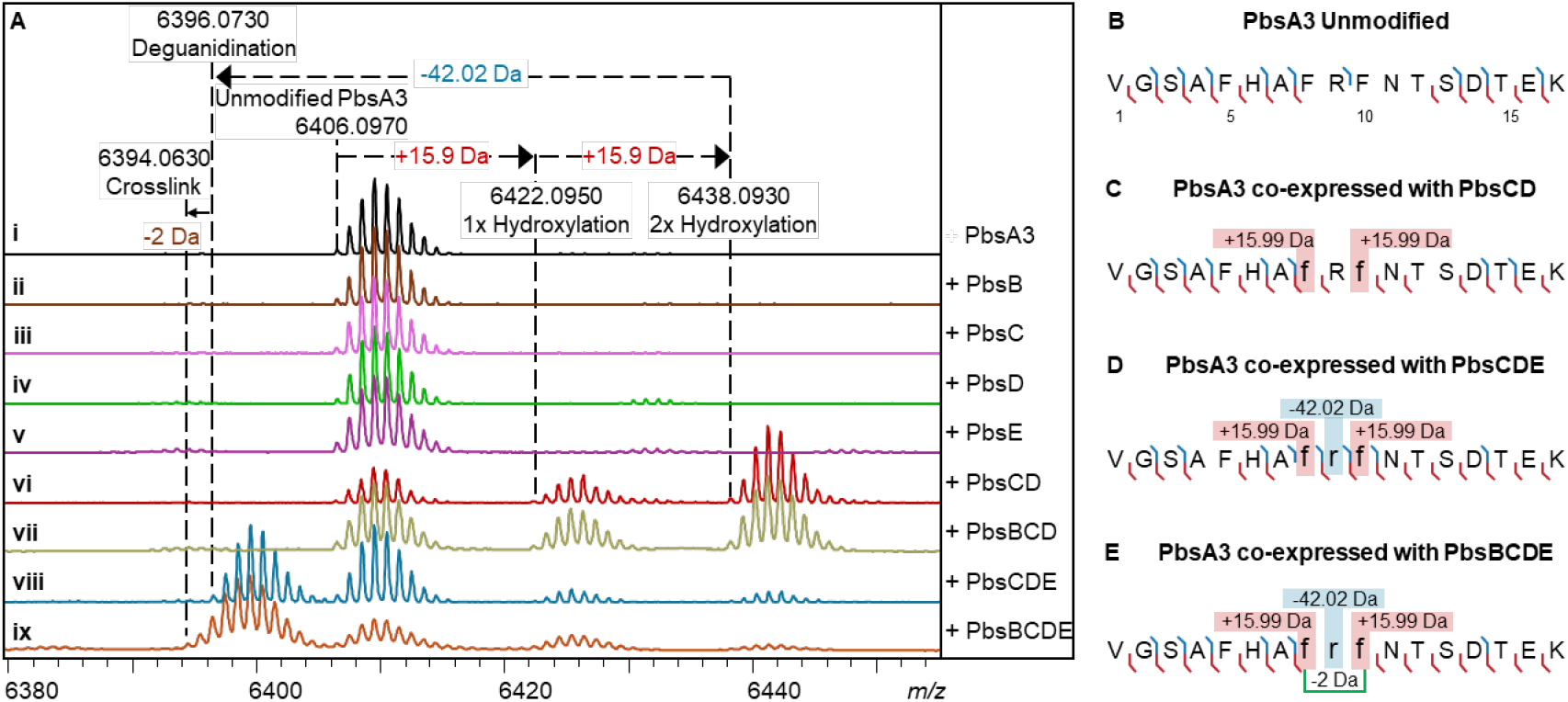
Mass spectrometric analysis of PbsA3 co-expressed with PbsB, PbsC, PbsD and PbsE in different combinations. (A) MALDI-TOF MS spectra of the PbsA3 peptides when expressed (i) alone; or co-expressed with (ii) the putative B12-dependent radical SAM, PbsB; (iii) the putative MNIO, PbsC; (iv) the putative RRE-containing partner protein, PbsD; (v) the putative arginase, PbsE; (vi) PbsC and PbsD; (vii) PbsB, PbsC and PbsD; (viii) PbsC, PbsD and PbsE; and (ix) PbsB, PbsC, PbsD and PbsE. HR-MS/MS-derived fragmentation for the endopeptidase GluC-digested peptide fragments are shown in (B) for unmodified PbsA3, (C) for PbsA3 co-expressed with PbsCD, (D) for PbsA3 co-expressed with PbsCDE and (E) for PbsA3 co-expressed with PbsBCDE. The peptide fragments after endoproteinase GluC digestion are numbered starting at V1 with b-ions displayed in blue, and y-ions displayed in red. The residues undergoing mass changes are shown in small font, further highlighted in boxes with their respective mass shifts. The hypothesized crosslinking residues are joined by a green bracket in panel E. For ESI MS/MS spectra, see Figure S4.

Through high resolution tandem mass spectrometry (HR-MS/MS) of the GluC endopeptidase-digested PbsCD product (C-terminal fragment starting at Val1 as shown in Figure 3B), we assigned each 15.99 Da mass gain to the Phe8 and Phe10 residues in the conserved FRF-motif (Figure 3C and Figure S4B; Table S1). Similarly, the mass loss of 42.02 Da in the PbsCDE product was localized to Arg9 in the FRF-motif (Figure 3D and Figure S4C; Table S1), suggesting that an Orn residue may be formed by the loss of a urea equivalent from the Arg side chain. However, for the PbsBCDE product, where an additional 2 Da mass loss was observed, no fragment ions were detected inside the FRF-motif (Figure 3E and Figure S4D; Table S1). This observation suggested the possibility of a crosslink being formed between the two modified Phe residues preventing fragmentation.

### Large scale purification of modified PbsA3

In an attempt to obtain the modified core peptide of PbsA3, we cloned the putative protease PbsP encoded in the *pbs* BGC into a pET28a vector with an N-terminal 6x His tag. However, our attempts at purifying PbsP were unsuccessful as the desired protein was not observed under various expression conditions. Therefore, endoproteinase GluC was used to access the modified fragment peptides. PbsA3 was co-expressed with PbsCD, PbsCDE and PbsBCDE in 5 L scale and the products were purified by immobilized metal affinity chromatography (IMAC). After desalting to remove imidazole, and proteolysis by GluC endoproteinase, IMAC was performed again to remove the His-tagged leader peptide fragment and the His-tagged GluC. The collected flowthrough was subjected to HPLC purification, which led to the isolation of the bis-hydroxylated PbsA3 C-terminal fragment (which we will term PbsA3-CD henceforth; 1.2 mg/L of culture), the bis-hydroxylated and deguanidinated product, PbsA3-CDE (0.5 mg/L of culture) and the bis-hydroxylated, deguanidinated and crosslinked product, PbsA3-BCDE (0.2 mg/L of culture).

### Structural elucidation of modified PbsA3

The 15.99 Da mass gains on Phe8 and Phe10 in PbsA3 when co-expressed with PbsCD are typical of hydroxylation events. To probe this hypothesis, we conducted advanced Marfey’s analysis^46^ on the HPLC-purified, GluC-digested PbsA3-CD product and compared the derivatized amino acids to different hydroxylated tyrosine standards (Figure S5). Based on the LC-MS analysis, we inferred that a putative hydroxyl moiety was either introduced at the *ortho*-position of the Phe rings (Cδ1 of the aromatic ring) or on the β-carbon. We next elucidated the structures of the three products PbsA3-CD, PbsA3-CDE and PbsA3-BCDE by one-dimensional (1D) and two-dimensional (2D) NMR experiments including ^1^H-^1^H TOCSY, ^1^H-^1^H NOESY, ^1^H-^13^C-HSQC and ^1^1H-^13^C HMBC.

For the PbsA3-CD peptide, all backbone and side chain protons including the 16 amide protons of the 17-mer peptide were observed in the TOCSY spectrum of the sample in 90% H_2_O and 10% D_2_O (Table S2). The aromatic protons of Phe8 and Phe10 showed different splitting-patterns from a normal aromatic group of a Phe residue. Their aromatic side chain protons integrated to four protons for each former Phe residue, and the 1D ^1^H-^1^H TOCSY spectra showed four protons that displayed two doublets and two triplets (Figure S6A). More detailed analysis of ^1^H-^13^C HSQC and HMBC spectra was consistent with the addition of a hydroxyl (OH) group to the aromatic side chains at the *ortho*-position (Cδ1, Figure 4A). Both β-protons of Phe8 and Phe10 showed a cross peak to the Cδ1 carbon carrying a hydroxyl group at 154.1 ppm (Figure 4B). This assignment is consistent with the 15.998 Da and 31.998 Da mass gains for the PbsA3-CD product that were observed by HR-MS/MS and indicated that both Phe residues in the conserved FRF-motif where hydroxylated.

**Figure 4.**
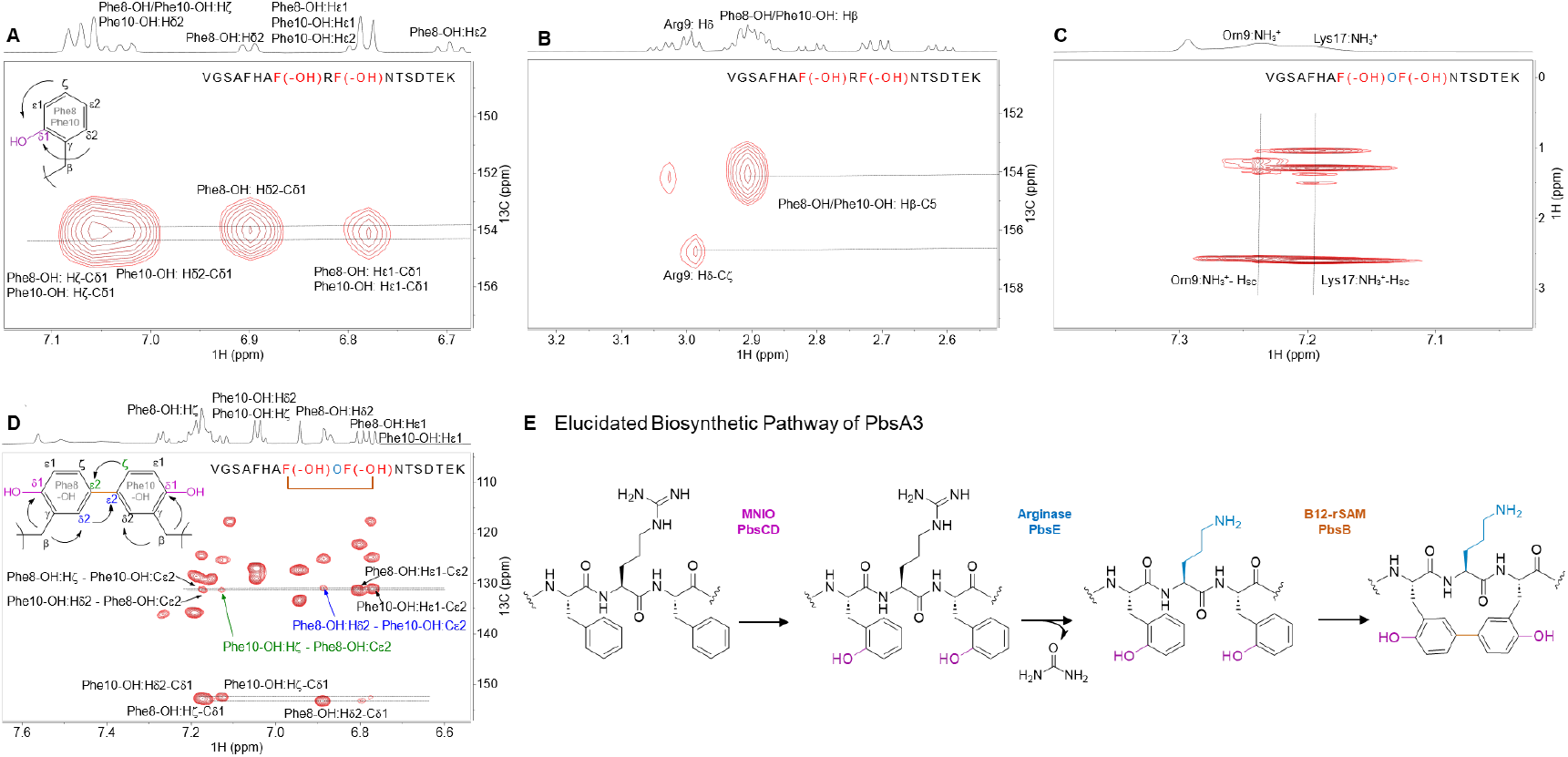
Structure determination and biosynthetic pathway of modified PbsA3 digested with endoproteinase GluC. (A) ^1^H-^13^C HMBC spectrum of PbsA3-CD highlighting the *ortho*-hydroxylated Phe8-OH and Phe10-OH residues. (B) ^1^H-^13^C HMBC spectrum highlighting the cross-peaks between the β-protons of Phe8-OH and Phe10-OH and their respective hydroxylated aromatic carbons. This part of the spectrum also shows a diagnostic cross peak for Arg9 that is consistent of an unmodified residue. (C) ^1^H-^1^H TOCSY spectra of PbsA3-CDE collected at 3 ^°^C in acidic condition (0.1% formic acid) highlighting the cross peaks of the NH^3+^ moiety on the C^δ^ of the residue at position 9 with the other side chain protons (labeled as H^sc^). (D) ^1^H-^13^C HMBC correlations of PbsA3-BCDE highlighting the crosslink formed between Cε2 of *ortho*-hydroxylated Phe8-OH and Phe10-OH residues. Cross peaks for the Hζ of Phe10-OH to the Cε2 carbon of Phe8-OH (in green arrow/font) and for the Hδ2-proton of Phe8-OH to the Cε2 carbon of Phe10-OH (in blue arrow/font) are shown. For panels A-D, the sequences of the analyzed peptides are shown in the spectra. Modified residues are in red (hydroxylated Phe8/10) or blue font (ornithine). Crosslinked residues are indicated in panel D. (E) Biosynthetic pathway of PbsA3 modified by PbsCD, followed by PbsE and PbsB.

Based on the advanced Marfey’s analysis the hydroxylated Phe residues co-eluted with both the L-*ortho*-tyrosine and DL-3-hydroxy-phenylserine standards. Hydroxylation of Cβ is inconsistent with the observed integration in the NMR spectra indicating four protons at each of the aromatic rings of Phe8 and Phe10 (Figure S6A). Additionally, the ^1^H-^13^C multiplicity edited HSQC showed the Cβ represented a methylene group and not a methine (Figure S6B). Thus, both Phe residues were hydroxylated at Cδ1.

To confirm the formation of Orn by PbsE and that no isobaric modification (e.g. epimerization) was performed on this residue by the rSAM PbsB, we conducted advanced Marfey’s analysis on the GluC-digested and HPLC-purified PbsA3-CDE and PbsA3-BCDE peptides (Figure S7). L-Orn, D-Orn and L-Arg were used as standards. LC-MS analysis of the derivatized samples and standards supported the conclusion that the Arg was converted into L-Orn by PbsE and that no epimerization had occurred.

This conclusion was confirmed by NMR spectroscopic analysis. The ^1^H spectrum of the PbsA3-CDE product was similar to that of PbsA3-CD (Figure S8). The 1D ^1^H-^1^H TOCSY showed four protons each for the aromatic side chains of Phe8 and Phe10 that was also observed in the PbsA3-CD product (Figure S9). The major difference between PbsA3-CDE and PbsA3-CD was the disappearance of the side chain NHε proton of Arg9 in the PbsA3-CDE spectrum (Figure S8B). In PbsA3-CD the NHε peak formed a sharp triplet at 6.99 ppm showing cross peaks with the other side chain protons and the amide proton of Arg9 in the TOCSY spectrum (Figure S10). However, in PbsA3-CDE, this peak was not detected (Figure S8B and Table S3). After lowering the temperature to 3 ^o^C to slow down exchange with bulk water and adding 0.1% formic acid to maintain the NHε in protonated state, a broad peak was observed at 7.46 ppm integrating for approximately three protons, which showed cross peaks in a TOCSY spectrum with the other side chain protons of the residue at the 9^th^ position in the peptide (Figure 4C and S11). All of these observations provided further evidence that Arg9 was transformed into an Orn residue.

For PbsA3-BCDE all backbone and side chain protons including 16 amide protons were observed in a TOCSY spectrum of a sample in 90% H_2_O and 10% D_2_O with 0.1% formic acid-d_2_ (^1^H and ^13^C NMR assignments in Table S4). Compared to the PbsA3-CD and PbsA3-CDE peptides, the aromatic protons of Phe8-OH and Phe10-OH in PbsA3-BCDE showed splitting patterns unlike the aromatic side chain of Phe and also different from the aromatic peaks observed in the PbsA3-CD and PbsA3-CDE peptides. In PbsA3-BCDE, integration of the aromatic signals from the former Phe8-OH and Phe10-OH indicated three protons each, and the 1D TOCSY spectrum (Figure S12) showed for each residue one doublet (d, *J* = 8.5 Hz), one doublet of doublets (dd, *J* = 8.5 Hz, 1.9 Hz) and one singlet (br). The ^1^H-^13^C HMBC revealed a C-C cross link between the Cε2-carbons of Phe8-OH and Phe10-OH (Figure 4D, E). The β-protons of both Phe8-OH and Phe10-OH showed cross peaks to the hydroxylated Cδ1 carbons, as well as to the Cδ2-carbons (Figure S13). Additionally, the presence of a cross link between the two Cε2-carbons of the aromatic rings of Phe8-OH and Phe10-OH was suggested by the observation of additional cross peaks between the two rings. For example, the peak at 7.12 ppm (Hζ of Phe10-OH) showed a cross peak to the Cε2 carbon of Phe8-OH at 131.3 ppm (Figure 4D, green arrow). Similarly, the Hδ2-proton at 6.89 ppm of Phe8-OH showed a cross peak to the Cε2 carbon of Phe10-OH at 131.0 ppm (Figure 4D, blue arrow). The HMBC bond connectivity of Phe8-OH and Phe10-OH in the PbsA3-BCDE peptide is depicted in Figure 4D. Furthermore, the NMR analysis confirmed an Orn instead of an Arg at position 9, as observed in the PbsA3-CDE peptide (Figure S14).

### Investigation of unmodified residues in the conserved FHAFRF motif

All enzymatic modifications are restricted to the FRF region of PbsA3, but the sequence that is fully conserved in all precursor peptides is considerably larger (AFHAFRF, Figure 2C). Therefore, we investigated the importance of the conserved Phe5 and His6 residues by replacement with Ala (F5A and H6A) and co-expression with PbsCD and PbsBCDE in *E. coli*. The F5A variant resulted in significantly diminished MNIO activity (Figure S15A; red spectra). No arginase- and B12-rSAM enzyme-mediated modification was observed, likely because of the low hydroxylation activity which is required for PbsB and PbsE activity (Figure S15A; orange spectra). The MNIO retained considerable activity with the H6A variant, resulting in wild type-like bis-hydroxylation (Figure S15B; red spectra), but the arginase and the rSAM enzymes did not introduce any further modifications (Figure S15B; orange spectra). Together, these results suggest that Phe5 is important for MNIO catalysis and that His6 is not necessary for MNIO activity but seems critical for the activity of the arginase PbsE and the B12-rSAM enzyme PbsB.

### *In vitro* reconstitution of enzyme activity

Following the successful reconstitution of the PbsA3 biosynthetic pathway in *E. coli*, we focused on *in vitro* reconstitution of the individual enzyme activities. For the purification of PbsCD, we constructed a pRSFDuet plasmid encoding an N-terminally 6xHis-tagged PbsC with a TEV protease cleavage site between the His-tag and the PbsC sequence. The plasmid also contained untagged *pbsD* under a second T7 promoter in the multiple cloning site (MCS) 2 of the pRSFDuet vector. When expressed in *E. coli*, PbsD co-purified with PbsC, validating that PbsD is an interacting partner for PbsC (Figure S16). HPLC-purified unmodified PbsA3 was reacted with the as-purified PbsCD, and no modification was observed (Figure 5A ii). Addition of freshly prepared FeSO^4^ to the reaction (under aerobic conditions) led to the formation of mono- and bis-hydroxylated products (Figure 5A iii), suggesting the involvement of Fe(II) ions in enzyme catalysis. Pre-incubation of as-purified PbsCD with sodium ascorbate followed by addition of PbsA3 also led to the formation of mono- and bis-hydroxylated products (Figure 5A iv).

**Figure 5.**
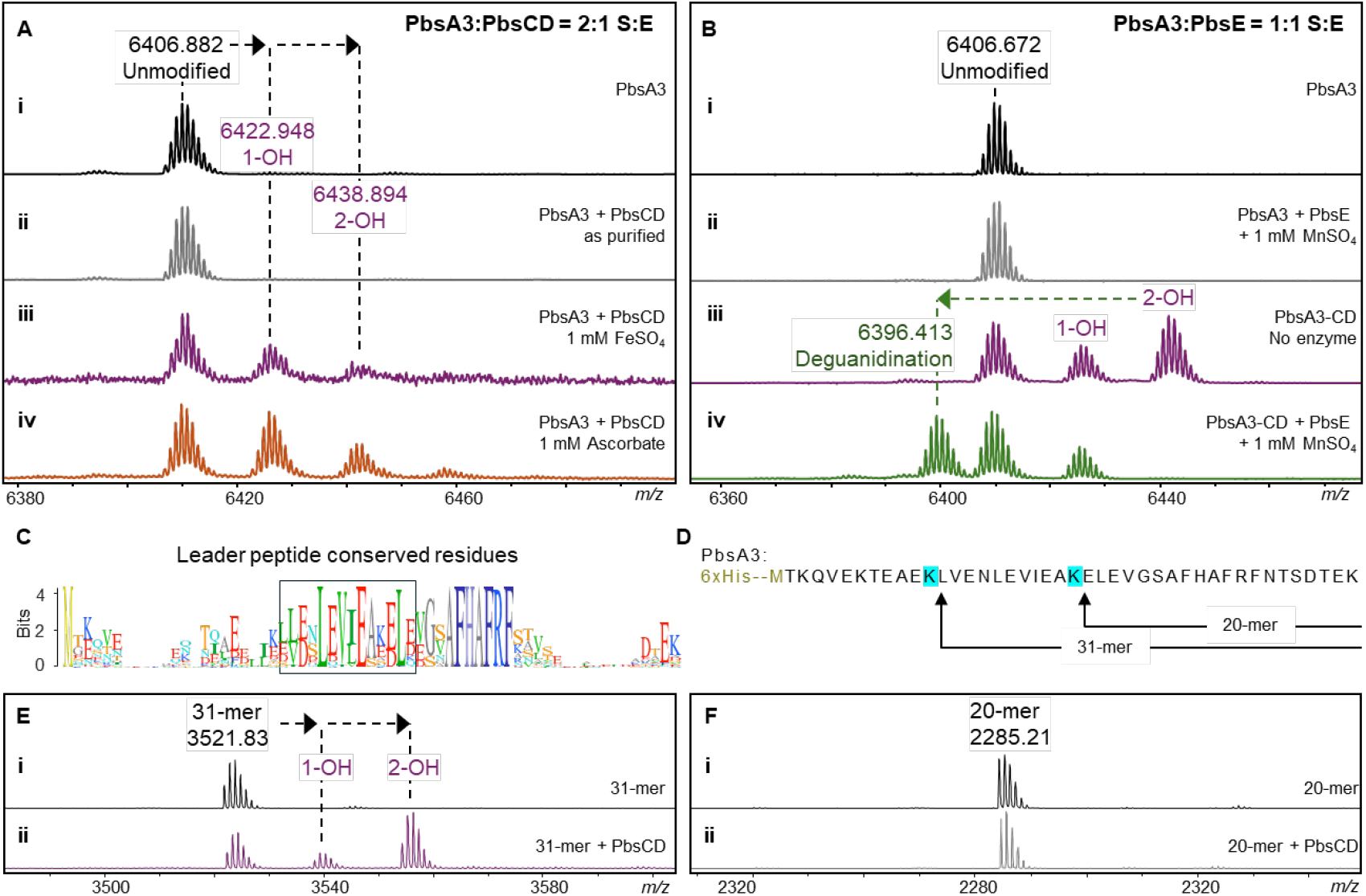
In vitro reconstitution of MNIO and arginase activity and minimal substrate determination for the MNIO. (A) MALDI-TOF MS spectra of PbsA3 after in vitro reaction with (i) no enzyme, (ii) PbsCD, as isolated, (iii) PbsCD, in the presence of FeSO_4_, and (iv) PbsCD, in the presence of sodium ascorbate. (B) MALDI-TOF mass spectra of PbsA3 after in vitro reaction with (i) no enzyme and (ii) PbsE in the presence of MnSO_4_. No arginase activity was detected on unmodified PbsA3. MALDI-TOF mass spectra of (iii) PbsCD-modified PbsA3, containing a mixture of unmodified, singly hydroxylated and bis-hydroxylated product and (iv) PbsA3-CD when reacted with PbsE in the presence of MnSO_4_. (C) Sequence logo of FHAFRF motif containing precursors showing conserved residues in the leader peptide, highlighted in the box. (D) Partial enzymatic hydrolysis of PbsA3 displaying the 20-mer and 31-mer fragments purified by HPLC. The 31-mer fragment contains the conserved residues of the leader peptide, which are absent in the 20-mer fragment. (E) MALDI-TOF mass spectra of the 31-mer fragment after in vitro reaction with (i) no enzyme, (ii) PbsCD in the presence of sodium ascorbate. (F) MALDI-TOF mass spectra of the 20-mer fragment after in vitro reaction with (i) no enzyme, and (ii) PbsCD in the presence of sodium ascorbate.

For the purification of PbsE, another plasmid was constructed to express the enzyme with an N-terminal 6xHis tag and TEV protease cleavage site. The purified 6xHis-TEV-PbsE fusion protein was directly used for *in vitro* assays (in the presence of manganese sulfate) with either unmodified PbsA3 (Figure 5B i, ii) or the PbsCD product containing PbsA3 and mono- and bis-hydroxylated PbsA3 (Figure 5B iii, iv). As observed in the *in vivo* co-expression studies, PbsE selectively modified the bis-hydroxylated product whereas unmodified PbsA3 and the mono-hydroxylated product did not undergo deguanidination (Figure 5B iv). Apart from establishing the *in vitro* activity of PbsE, this observation validated our hypothesis that the PbsCD-mediated hydroxylation is the first step in the biosynthetic pathway, providing the substrate for the arginase PbsE.

### Minimal substrate required for MNIO activity

Sequence alignment of the predicted precursors containing the conserved core motifs also revealed a conservation pattern in the putative leader peptide (LP) region (Figure 5C). To investigate the importance of these residues we made truncated substrates by conducting a partial digestion of unmodified PbsA3 peptide using LysC endopeptidase. After 15 min at room temperature with 1:10000 enzyme:substrate, peptide fragments of varying length were isolated by HPLC (Figure 5D) including a 20-mer fragment without the conserved LP stretch, and a 31-mer fragment with the conserved stretch of amino acids in the LP (Figure 5D). These purified fragments were reacted with PbsCD *in vitro*, which resulted in the successful bis-hydroxylation of the 31-mer fragment (Figure 5E). No modification was observed for the 20-mer fragment, suggesting that the conserved LP-region is important for MNIO activity (Figure 5F).

### Determining the substrate scope of PbsCD

To investigate the possibility of a required order of hydroxylation of Phe8 and Phe10, we mutated these residues to Ala. When co-expressed with PbsCD or PbsBCDE in *E. coli*, the F8A and F10A variants (hereafter named **A**RF and FR**A**-variants, respectively, mutated residues in bold) underwent only one hydroxylation (Figure S17A, S17B). Using HR-MS/MS we assigned these hydroxylations to the Phe residues in the **A**R<u>F</u>- and <u>F</u>R**A**-variants, respectively (Figure S18, S19; hydroxylated residues underlined). Upon co-expression with PbsBCDE, no arginase activity and no PbsB-mediated crosslinking was detected for either peptide suggesting that both Phe8 and Phe10 need to be hydroxylated for these activities. As expected based on the data in the previous sections, the ARA variant of PbsA3 was not modified by any of the enzymes (Figure S17C).

The SSN cluster harboring PbsC (Figure 2A) also contains enzymes that are encoded in BGCs with precursor peptides containing alternative conserved motifs in the core region, such as FHTFMF and YHxYxY motifs. We therefore made a series of variants of PbsA3 to investigate if the Pbs enzymes would accept these as substrates. The **Y**RF, FR**Y** and **Y**R**Y** variants were processed by PbsCD (Figure S20) resulting in mostly one and two hydroxylations (Figure S21-S23). The patterns observed in HR-MS/MS analysis showed the hydroxylation to occur on the aromatic residues. The fragment ions for the doubly hydroxylated **Y**R**Y** product suggested that both Tyr8 and Tyr10 were modified (Figure S23). Co-expression of the **Y**RF, FR**Y** and **Y**R**Y** variants with PbsBCDE resulted in successful arginase activity for all three variants (Figure S20). Only the **Y**RF and FR**Y** variants were accepted as substrates for PbsB resulting in a crosslink as inferred from the 2 Da mass loss in HR-MS. These data show that PbsCD can hydroxylate Tyr residues, consistent with the motifs seen in the substrates for PbsCD orthologs.

### Characterization of tyrosine modifications catalyzed by *pbs* enzymes

PbsA3-YRY variants were bis-hydroxylated by PbsCD, but not crosslinked by PbsB (Figure S20C). We therefore replaced the FRF motif in PbsA3 with the YTY motif found in the precursor from the *stg* BGC (Figure S2) in the genome of *Streptomyces turgidiscabies* ATCC 700248 (YHTYTY-motif). Upon co-expression with PbsCD, the PbsA3-YTY precursor was successfully bis-hydroxylated (YTY-CD product; Figure S24B and S25B; Table S1). A mono-hydroxylated product (YTY-CD 1-OH; Figure S24B; Table S1) was also detected with the hydroxylation localized on Tyr10 as determined by HR-MS/MS (Figure S25A). We also observed crosslinking of the PbsA3-YTY precursor that was not hydroxylated upon co-expression with PbsBCD (YTY-B product; Figure S24B and S26A). In addition to the aforementioned products, we also observed a minor bis-hydroxylated and crosslinked product (YTY-BCD 2-OH) along with a minor mono-hydroxylated-crosslinked product (YTY-BCD 1-OH; Figure S24B and S26B; Table S1).

### Structure elucidation of YTY-B and the two products of YTY-CD

Large scale purification of the PbsA3-YTY products from co-expression with PbsCD (YTY-CD) and PbsB (YTY-B) was conducted followed by GluC endoproteinase digestion and subsequent HPLC purification. The NMR structures of the YTY-B, and mono and bishydroxylated YTY-CD peptides were determined by 1D and 2D NMR experiments (^1^H-^1^H TOCSY, ^1^H-^1^H NOESY, ^1^H-^13^C-HSQC and ^1^1H-^13^C HMBC). For the YTY-B peptide, all backbone and side chain protons including 16 amide protons of the 17-mer peptide, were observed in the TOCSY spectrum for the sample in 90% H_2_O and 10% D_2_O and 0.1% formic acid-d_2_ (Table S5). The aromatic protons of Tyr8 and Tyr10 showed different splitting patterns from a normal Tyr, integrating to three protons each. The 1D TOCSY spectrum showed three protons containing two doublet peaks and 1 singlet (or 1 doublet with a small J coupling of 2.5 Hz; Figure S27). More detailed analysis of ^1^H-^13^C HSQC (Figure S28) and HMBC (Figure S29) data revealed a C-C cross link between the two aromatic rings of Tyr8 and Tyr10 at the Cε2-position. The Hδ2 proton at 6.64 ppm of Tyr8 showed a cross peak to the Cε2 carbon at 125.4 ppm of Tyr10. Both beta protons of Tyr8 and Tyr10 showed NOE cross peaks to their respective Hδ1 and Hδ2 protons in a ^1^H-^1^H NOESY spectrum, which provides additional support that the C-C cross link occurred between the Cε2-carbons of the two rings (Figure S30). The structure in Figure 6 is also consistent with the HR-MS and HR-MS/MS analysis as shown in Figure S24B and S26A, respectively.

**Figure 6.**
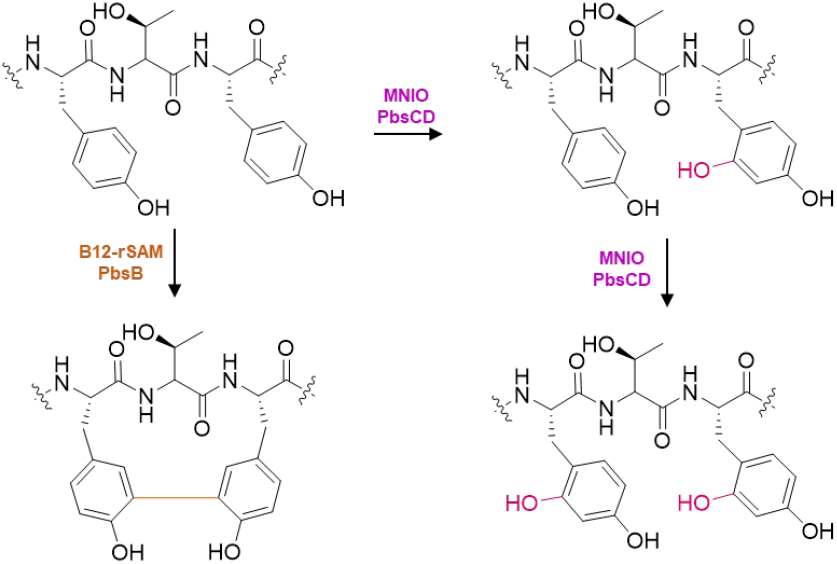
PbsA3-YTY variant modification catalyzed by PbsBCD. The B12-rSAM enzyme PbsB-mediates crosslinking of Tyr residues in the PbsA3-YTY variant. PbsCD first hydroxylates Tyr10 residue and then Tyr8.

The mono and bishydroxylated YTY-CD peptides co-eluted during HPLC purification. Hence, their structural characterization was conducted as a mixture. Although the ^1^H spectrum is complicated, two sets of signals with a ratio of 2:3 (based on peak integration values) were clearly observed. The 1D ^1^H-^1^H TOCSY spectrum in the aromatic region revealed two distinct types of spin systems (Figure S31). One displayed the original pattern of a Tyr residue, the other showed three peaks for each ring: one doublet, one doublet of doublets, and one singlet. Based on the proton integration values, one of the peptides contains one intact Tyr residue and one modified Tyr residue, whereas the other contained two modified Tyr residues. Further analysis of 2D ^1^H-^1^H TOCSY (Figure S32), ^1^H-^13^C HSQC (Figure S33) and HMBC (Figure S34) revealed that the modified Tyr residue was hydroxylated at the Cδ1 position. The proton Hδ2 at 6.80 ppm of Tyr8 in the bishydroxylated YTY-CD peptide showed a cross peak to the Cδ1 carbon at 155.3 ppm and a cross peak to the Cζ carbon at 155.7 ppm (carrying the original −OH of Tyr). Likewise, the Hδ2 proton at 6.87 ppm of Tyr10 in monohydroxylated YTY-CD showed a cross peak to the Cδ1 carbon at 155.3 ppm and a cross peak to its original Cζ-OH at 155.7 ppm (Figure S34). The beta-protons of Tyr10 in this peptide showed cross peaks to their respective downfield shifted Cδ1 carbon at 155.3 ppm (Figure S35), confirming the location of the hydroxylation. The assignments for both peptides are given in Table S6, and the structures of both peptides are consistent with the HR-MS/MS results (Figure S25). Collectively, these data show that the site of hydroxylation by PbsCD is the same for Phe and Tyr, and that PbsB crosslinks Tyr, *ortho-*Tyr and hydroxy-Tyr residues with the same regiochemistry.

### AlphaFold 3 prediction of Pbs pathway enzymes in complex with precursor peptides

To understand the mechanism of substrate recognition and modification by PbsCD, we generated AlphaFold 3 models^47^ of PbsCD bound to PbsA1-A4 and three iron ions (Figure S36; model shown for PbsCD in complex with PbsA2). The predicted model showed PbsC and PbsD as interacting heterodimeric molecules, consistent with co-purification of untagged PbsD with tagged PbsC (Figure S16). The per-residue measure of local confidence (pLDDT) values for both proteins are generally very high (>90), with lower confidence in the orientation of the PbsA substrate peptides. The model suggests an anti-parallel β-sheet interaction between a conserved region of the leader peptide of the four precursors (Figure S36B, S37) with the RiPP recognition element (RRE) in the PbsD partner protein, as observed in various other enzymes for which the substrate engagement has been structurally characterized.^48–55^ The model placed three Fe ions in the putative active site of the enzyme (Figure S36C), consistent with other MNIO structures, although it is still unclear whether the active form of this family of enzymes contains two or three Fe ions.^23^ His97, Asp133 and Glu178 potentially coordinate with one Fe ion, His216, Asp260 and His262 may coordinate with the second Fe ion, and Asp213, His247 and Glu290 are predicted to coordinate with the third iron. Interestingly, for PbsA1, A2 and A4, although the pLDDT values are considerably lower, the predicted substrate bound structure placed one of the Phe residues in close proximity to the Fe ions in the enzyme active site (Figure S38A, S38B, S38D). However, for the PbsA3-complexed model, the His residue (His5) of the FHAFRF motif was placed in proximity to the active site metals (Figure S38C). Since PbsA3 has been experimentally shown to undergo modification at Phe8/Phe10 residues, the prediction shows the limitations of AlphaFold 3 and the need for experimental validations of the model. Nevertheless, the predicted complexes shed light on the putative RRE-containing partner protein that may interact with the leader region, helping the enzyme recognize the substrate.

A similar prediction was also made for the arginase PbsE in conjunction with the precursors PbsA1-A4 and two manganese ions (Figure S39A). The putative metal-coordinating residues in the active site were predicted to be His10, Asp31, His33, Asp35, Asp165 and Asp167 (Figure S39B). The prediction model placed the modified Arg9 residue of the FHAFRF motif proximal to the Mn ions, with Arg9 from PbsA4 in the closest proximity compared to the other precursor complexes (Figure S39D-S39G). In general, we believe that the complexes of the arginase and the unmodified precursor predictions may not be completely reliable, considering the specific selectivity of PbsE for PbsCD-modified bis-hydroxylated intermediates as the substrate. We did not use AlphaFold to model the B12-rSAM PbsB and the unmodified precursor interactions since the enzyme selectively crosslinks bis-hydroxylated and deguanidinated intermediates, which are not accessible in the current versions of AlphaFold3.

## Discussion

Enzymes involved in posttranslational modifications (PTMs) of RiPPs offer a plethora of diverse and interesting chemical transformations that are challenging to perform synthetically.^20^ To identify enzymes with new functions, SSNs are a valuable tool to group thousands of uncharacterized enzymes based on variable sequence similarity cut offs. In many cases, enzymes that make up one cluster have been reported as being isofunctional.^56,57^ In this study, we used an SSN for MNIOs and focused on associated substrate peptides containing conserved motifs that were predicted to result in novel chemistry. Indeed, investigation of one such BGC uncovered three different metalloenzymes involved in the PTM of a conserved FHAFRF motif-containing RiPP precursor from the genome of *Peribacillus simplex* BE23.

Thus far, MNIO-catalyzed reactions have been characterized that involve modification of Cys, Asn, and Asp residues (Figure 1);^23^ aromatic residues have not been reported as sites for MNIO modification. The hydroxylation of aromatic side chains reported in this study thereby further expands the diverse range of chemical reactions performed by MNIOs. Hydroxylation of aromatic amino acids has been reported to be catalyzed by heme-containing cytochrome P450 enzymes, pterin-dependent hydroxylases, flavin-dependent monooxygenases, nonheme mononuclear iron dioxygenases and diiron hydroxylases.^58–61^ While enzymatic *ortho*-hydroxylation of phenolic moieties has been reported for certain natural products,^62^ reports on Cδ1-hydroxylation of Phe residues in RiPPs are scarce. The only known example is in the case of thioviridamide-like compounds where *ortho*-hydroxylation of a Phe residue is catalyzed by a cytochrome P450 enzyme.^63^

AlphaFold3 prediction models of the MNIO and precursor substrate complexes provide structural insights of the interaction of the MNIO and precursor peptides. The RRE-containing partner protein PbsD likely recognizes the substrate through an anti-parallel beta sheet interaction with the leader region of the precursor peptide. Consistent with the model, deletion of the leader fragment predicted to form the beta-sheet led to loss of enzymatic processing (Figure 5F). Interestingly, in all the *pbs* precursors except PbsA3, one of the Phe residues involved in modification was placed close to the Fe ions in the predicted active site of the MNIO in the AlphaFold3 models. The mechanism of hydroxylation of the two Phe residues in the FRF motif is at present not clear. Most MNIOs catalyze four-electron oxidations of their substrates, shuttling the four electron equivalents to molecular oxygen and regenerating a ferrous ion that can initiate the next catalytic cycle.^23^ Only one MNIO has thus far been shown to catalyze a two-electron oxidation, MovX that oxidatively cleaves the Cα-N bond in an Asn residue generating hydrogen peroxide in the process (Figure 1).^28^ Furthermore, thus far, all proposed MNIO mechanisms have involved C-H activation at an sp^3^-hybridized carbon, which has been hypothesized to be carried out by a ferric superoxide species.^23^ The double hydroxylation of two Phe residues in PbsA in principle could be a four-electron process in which both oxygens from O_2_ are incorporated into two different Phe residues, but that would require the oxidations of the two Phe residues to be carried out by two different oxidating species on PbsC (e.g. Figure S40). Alternatively, the reaction could involve two sequential two-electron oxidations that each involve a different molecule of O_2_. In other enzymes that carry out aromatic ring oxidations such as cytochrome P450 or tetrahydrobiopterin-dependent enzymes, an Fe(IV)-oxo is required.^60,61^ If PbsC uses an Fe(IV)-oxo or a peroxo species for the two-electron oxidation of one of the Phe residues (Figure 7A-B), reducing equivalents would be needed from the environment. The observation that ascorbate activated the enzyme (Figure 5A-iv) could be consistent with such a mechanism, but detailed studies will be needed to distinguish the various possibilities.

**Figure 7.**
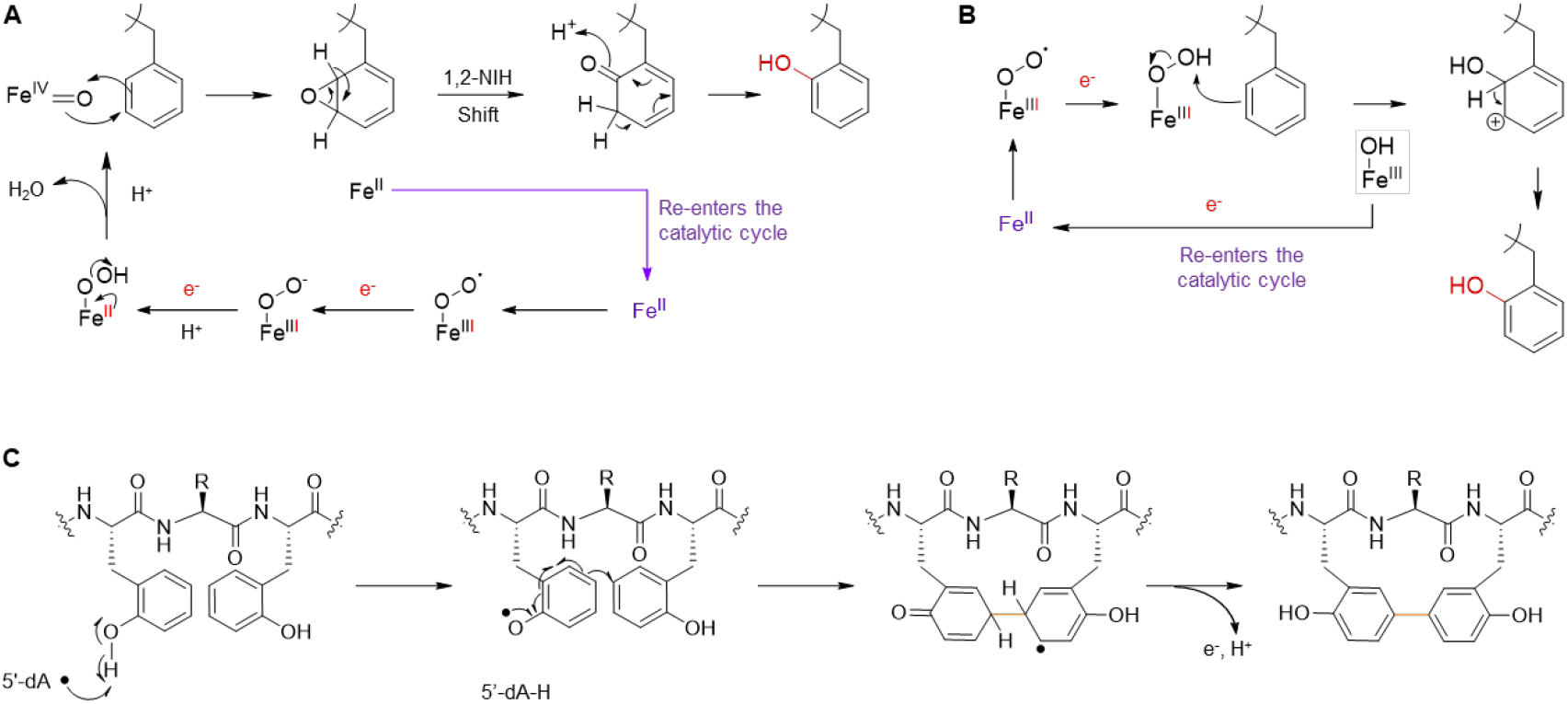
Proposed reaction mechanism for the MNIO and B12-dependent rSAM enzyme from the *pbs* BGC. (A) Possible mechanism of hydroxylation catalyzed by the MNIO PbsC using an Fe(IV)-oxo through an epoxide intermediate followed by a hydride shift (1,2-NIH shift).^64^ Other mechanisms that do not involve a hydride migration (electrophilic aromatic substitution like) can also be drawn. (B) An alternate mechanism potentially initiated by a Fe(III)-peroxo species. (C) Proposed mechanism of C-C crosslinking between the hydroxylated Phe8 and Phe10 residues initiated by hydrogen abstraction by a 5’-deoxyadenosyl radical.

Ornithine-containing peptides are found in both non-ribosomal peptides and RiPPs. Orn is present in various bioactive compounds such as daptomycin,^65^ gramicidin,^66^ and landornamide.^67^ In the latter case, two ornithine residues are installed by the arginase OspR, which showed considerable substrate tolerance.^68^ In contrast, the arginase PbsE selectively acts on an Arg residue that is bordered by hydroxylated Phe residues. Such selectivity perhaps guides the directionality of the *pbs* biosynthetic pathway and provides the substrate for downstream processing by the B12-dependent rSAM enzyme. Interestingly, orthologous precursor peptides missing the conserved Arg residue in the precursor peptide also lacked the gene for an arginase in the BGC suggesting a precursor-mediated evolution of the BGCs across different organisms.

C-C crosslink formation between the aromatic side chains of Tyr has been reported for cytochrome P450 enzymes involved in the biosynthesis of cittilin,^69^ arylomycin,^70^ the biarylitides,^71–74^ pyruvatides,^29^ and vancomycin,^75,76^ but has not been reported for B12-rSAMs. Other families of rSAM enzymes generate crosslinks between the side chains of various aliphatic amino acids and the side chains of aromatic amino acids in the triceptide group of RiPPs.^77,78^ The mechanism of these latter reactions involves hydrogen atom abstraction from the aliphatic amino acid and then addition of the resulting carbon-based radical to the aromatic rings of the partner amino acid.^79,80^ PbsB on the other hand crosslinks two aromatic amino acids, a process that has not been previously reported for a B12-rSAM enzyme. Originally believed to be limited to methylation,^81^ this group of enzymes has been implicated in a variety of other transformations including ring rearrangement reactions.^82^ Regarding the possible mechanism of the PbsB-catalyzed crosslinking reaction, the required MNIO-catalyzed introduction of hydroxyl groups on two Phe residues likely activates the aromatic rings, by making the aromatic ring more electron rich and by providing a site for hydrogen atom abstraction from the phenol. We suggest the formation of an *ortho*-Tyr radical by hydrogen atom transfer to the canonical 5’-deoxyadenosyl radical (Figure 7B), which results in spin density at the Cε2 carbon. Addition to the partner *ortho-*Tyr in the confines of the enzyme active site would result in an aromatic radical that can lose an electron (possibly to an FeS cluster as shown for streptide)^79^ and a proton to rearomatize the ring (tautomerization also rearomatizes the other ring). It is not clear, however, why the reaction requires a vitamin B12-dependent rSAM enzyme as the proposed mechanism does not suggest an obvious role for methylcobalamin. It is possible that the B12-dependence is vestigial or that it acts as an electron transfer conduit as suggested for the ring contraction B12-rSAM enzyme enzyme.^83^ Our data show that the YTY sequence gave rise to the same crosslinking pattern, consistent with the hypothesis that cyclization requires phenolic substrates. The position of hydroxylation (on Cδ1 or on Cζ) appears not critical for cyclization as the resulting tyrosyl radicals are both activated for coupling at Cε2. But if the mechanism in Figure 7C is indeed operational, then it is surprising that the 5’-deoxy adenosyl radical would be able to perform hydrogen atom abstraction from both a phenolic O-H bond at the δ1 position as well as the ζ position. Another interesting question is why MNIO activation of Phe residues would have evolved when Tyr residues apparently give the same cyclized framework. One possibility is that crosslinking of two *ortho-* Tyr residues probably does not result in two different atropisomeric structures as the rotational barrier of the biphenyl rings is likely not sufficiently high (assuming the cyclic peptide does not increase the barrier by conformational restrictions). Cyclization of two regular (not *ortho*) Tyr residues on the other hand will give rise to two atropisomeric structures, one of which is likely favored in the context of the enzyme active site, as seen for instance in the P450-catalyzed Tyr-Tyr crosslinks in pyruvatides.^29^ It is possible that the less conformationally constrained crosslink of two *ortho*-Tyr residues is favorable for the bioactivity of the final product of the *pbs* BGC. Investigation of this hypothesis will first require identification of the final structure after leader peptide removal which requires reconstitution of the protease encoded in the BGC; thus far this goal has not yet been achieved. The protease has sequence homology with the TldDE protease family,^84–86^ but unlike this family which usually constitutes a heterodimer, the *pbs* BGC contains only a single gene (*pbsP*) and a partner protein has not been identified in the genome.

The crosslinked peptides produced by the enzymes of the *pbs* BGC are examples of a growing number of RiPPs that generate one or more three-amino-acid membered rings that can be generally described as biaryl linked peptides ΩXΩ (Ω = aromatic amino acid; X = variable amino acid), or aryl-aliphatic crosslinked peptides ΩXY and YXΩ (Y = any amino acid crosslinked on an sp^3^ carbon of its side chain). These include the triceptides,^87,88^ biarylitides,^71–74,89–91^, dynobactin^92^ and darobactin.^93^ The crosslinks are introduced by both P450^73,94,95^ and rSAM enzymes^20,96^ and are diverse in structure but remarkably uniform in consisting of three amino acids. Only for dynobactin and darobactin are the biological activities well understood, with the crosslinked peptides generating conformationally restricted peptides that are locked in a β-strand-like mimic and that interact with a β-sheet of their target.^97^ Whether this is a general theme remains to be seen.

## AUTHOR INFORMATION

### Corresponding Authors

Wilfred A. van der Donk −Department of Chemistry and Howard Hughes Medical Institute, University of Illinois at Urbana-Champaign, Urbana, IL, United States.

### Authors

Chandrashekhar Padhi – Department of Chemistry and Howard Hughes Medical Institute, University of Illinois at Urbana-Champaign, Urbana, IL, United States.

Lingyang Zhu – School of Chemical Sciences NMR Laboratory, University of Illinois at Urbana-Champaign, Urbana, IL, United States.

Jeff Y. Chen – Department of Chemistry, the Carl R. Woese Institute for Genomic Biology, and the Howard Hughes Medical Institute at the University of Illinois at Urbana-Champaign, 1206 W Gregory Drive, Urbana, IL 61801, United States.

Ryan Moreira – Department of Chemistry and Howard Hughes Medical Institute, University of Illinois at Urbana-Champaign, Urbana, IL, United States.

### Author contributions

**C.P**. designed the study. **C.P**., **L.Z**. and **W.A.V**. prepared the manuscript. **C.P**., **L.Z**. and **J.Y.C**. conducted experiments. **J.Y.C**. performed in vitro reconstitution of MNIO activity. **L.Z**. and **R.M**. analyzed NMR data.

### Data availability

All data generated in this study are provided in the Supplementary Information accompanying this paper including materials and methods, Figures S1-S40, Tables S1-S7.

## ACKNOWLEDGMENTS

We thank Denys Lytviak for help with preparing reagents for experiments. R.M. was supported by the Life Science Research Foundation. This work was supported in part by a grant from the National Institutes of Health (GM058822 to W.A.vdD.). W.A.vdD is an Investigator of the Howard Hughes Medical Institute. A Bruker UltrafleXtreme mass spectrometer used was purchased with support from the National Institutes of Health (S10 RR027109). HHMI lab heads have previously granted a nonexclusive CC BY 4.0 license to the public and a sublicensable license to HHMI in their research articles. Pursuant to those licenses, the author-accepted manuscript of this article can be made freely available under a CC BY 4.0 license immediately upon publication.

## References

(1) Fryszkowska, A.; Devine, P. N. Biocatalysis in drug discovery and development. Curr. Opin. Chem. Biol. 2020, 55, 151–60.

(2) Wu, S.; Snajdrova, R.; Moore, J. C.; Baldenius, K.; Bornscheuer, U. T. Biocatalysis: enzymatic synthesis for industrial applications. Angew. Chem. Int. Ed. 2021, 60, 88–119.

(3) Intasian, P.; Prakinee, K.; Phintha, A.; Trisrivirat, D.; Weeranoppanant, N.; Wongnate, T.; Chaiyen, P. Enzymes, in vivo biocatalysis, and metabolic engineering for enabling a circular economy and sustainability. Chem. Rev. 2021, 121, 10367–451.

(4) Buller, R.; Lutz, S.; Kazlauskas, R. J.; Snajdrova, R.; Moore, J. C.; Bornscheuer, U. T. From nature to industry: Harnessing enzymes for biocatalysis. Science 2023, 382, eadh8615.

(5) Savile, C. K.; Janey, J. M.; Mundorff, E. C.; Moore, J. C.; Tam, S.; Jarvis, W. R.; Colbeck, J. C.; Krebber, A.; Fleitz, F. J.; Brands, J. et al. Biocatalytic asymmetric synthesis of chiral amines from ketones applied to sitagliptin manufacture. Science 2010, 329, 305–9.

(6) Huffman, M. A.; Fryszkowska, A.; Alvizo, O.; Borra-Garske, M.; Campos, K. R.; Canada, K. A.; Devine, P. N.; Duan, D.; Forstater, J. H.; Grosser, S. T. et al. Design of an in vitro biocatalytic cascade for the manufacture of islatravir. Science 2019, 366, 1255–9.

(7) McIntosh, J. A.; Benkovics, T.; Silverman, S. M.; Huffman, M. A.; Kong, J.; Maligres, P. E.; Itoh, T.; Yang, H.; Verma, D.; Pan, W. et al. Engineered Ribosyl-1-Kinase Enables Concise Synthesis of Molnupiravir, an Antiviral for COVID-19. ACS Cent. Sci. 2021, 7, 1980–5.

(8) McIntosh, J. A.; Liu, Z.; Andresen, B. M.; Marzijarani, N. S.; Moore, J. C.; Marshall, N. M.; Borra-Garske, M.; Obligacion, J. V.; Fier, P. S.; Peng, F. et al. A kinase-cGAS cascade to synthesize a therapeutic STING activator. Nature 2022, 603, 439–44.

(9) Reisenbauer, J. C.; Sicinski, K. M.; Arnold, F. H. Catalyzing the future: recent advances in chemical synthesis using enzymes. Curr. Opin. Chem. Biol. 2024, 83, 102536.

(10) Finnigan, W.; Lubberink, M.; Hepworth, L. J.; Citoler, J.; Mattey, A. P.; Ford, G. J.; Sangster, J.; Cosgrove, S. C.; da Costa, B. Z.; Heath, R. S. et al. RetroBioCat database: a platform for collaborative curation and automated meta-analysis of biocatalysis data. ACS Catal. 2023, 13, 11771–80.

(11) Arnold, F. H.; Volkov, A. A. Directed evolution of biocatalysts. Curr. Opin. Chem. Biol. 1999, 3, 54–9.

(12) van der Donk, W. A. Introduction: unusual enzymology in natural product synthesis. Chem. Rev. 2017, 117, 5223–5.

(13) Shen, B. A New Golden Age of Natural Products Drug Discovery. Cell 2015, 163, 1297–300.

(14) Newman, D. J.; Cragg, G. M. Natural products as sources of new drugs over the nearly four decades from 01/1981 to 09/2019. J. Nat. Prod. 2020, 83, 770–803.

(15) Atanasov, A. G.; Zotchev, S. B.; Dirsch, V. M.; Orhan, I. E.; Banach, M.; Rollinger, J. M.; Barreca, D.; Weckwerth, W.; Bauer, R.; Bayer, E. A. et al. Natural products in drug discovery: advances and opportunities. Nat. Rev. Drug Discov. 2021, 20, 200–16.

(16) Montalbán-López, M.; Scott, T. A.; Ramesh, S.; Rahman, I. R.; van Heel, A. J.; Viel, J. H.; Bandarian, V.; Dittmann, E.; Genilloud, O.; Goto, Y. et al. New developments in RiPP discovery, enzymology and engineering. Nat. Prod. Rep. 2021, 38, 130–239.

(17) Cao, L.; Do, T.; Link, A. J. Mechanisms of action of ribosomally synthesized and posttranslationally modified peptides (RiPPs). J. Ind. Microbiol. Biotechnol 2021, 48, kuab005.

(18) Ongpipattanakul, C.; Desormeaux, E. K.; DiCaprio, A.; van der Donk, W. A.; Mitchell, D. A.; Nair, S. K. Mechanism of action of ribosomally synthesized and post-translationally modified peptides. Chem. Rev. 2022, 122, 14722–814.

(19) Rebuffat, S. Ribosomally synthesized peptides, foreground players in microbial interactions: recent developments and unanswered questions. Nat. Prod. Rep. 2022, 39, 273–310.

(20) Nguyen, D. T.; Mitchell, D. A.; van der Donk, W. A. Genome mining for new enzyme chemistry. ACS Catal. 2024, 14, 4536–53.

(21) Zdouc, M. M.; van der Hooft, J. J. J.; Medema, M. H. Metabolome-guided genome mining of RiPP natural products. Trends. Pharmacol. Sci. 2023, 44, 532–41.

(22) Padhi, C.; Field, C. M.; Forneris, C. C.; Olszewski, D.; Fraley, A. E.; Sandu, I.; Scott, T. A.; Farnung, J.; Ruscheweyh, H.-J.; Narayan Panda, A. et al. Metagenomic study of lake microbial mats reveals protease-inhibiting antiviral peptides from a core microbiome member. Proc. Natl. Acad. Sci. USA 2024, 121, e2409026121.

(23) Chen, J. Y.; van der Donk, W. A. Multinuclear non-heme iron dependent oxidative enzymes (MNIOs) involved in unusual peptide modifications. Curr. Opin. Chem. Biol. 2024, 80, 102467.

(24) Kenney, G. E.; Dassama, L. M. K.; Pandelia, M.-E.; Gizzi, A. S.; Martinie, R. J.; Gao, P.; DeHart, C. J.; Schachner, L. F.; Skinner, O. S.; Ro, S. Y. et al. The biosynthesis of methanobactin. Science 2018, 359, 1411–6.

(25) Ting, C. P.; Funk, M. A.; Halaby, S. L.; Zhang, Z.; Gonen, T.; van der Donk, W. A. Use of a scaffold peptide in the biosynthesis of amino acid–derived natural products. Science 2019, 365, 280–4.

(26) Leprevost, L.; Jünger, S.; Lippens, G.; Guillaume, C.; Sicoli, G.; Oliveira, L.; Falcone, E.; de Santis, E.; Rivera-Millot, A.; Billon, G. et al. A widespread family of ribosomal peptide metallophores involved in bacterial adaptation to metal stress. Proc. Natl. Acad. Sci. USA 2024, 121, e2408304121.

(27) Ayikpoe, R. S.; Zhu, L.; Chen, J. Y.; Ting, C. P.; van der Donk, W. A. Macrocyclization and backbone rearrangement during RiPP biosynthesis by a SAM-dependent domain-of-unknown-function 692. ACS Cent. Sci. 2023, 9, 1008–18.

(28) Chioti, V. T.; Clark, K. A.; Ganley, J. G.; Han, E. J.; Seyedsayamdost, M. R. N-Cα bond cleavage catalyzed by a multinuclear iron oxygenase from a divergent methanobactin-like RiPP gene cluster. J. Am. Chem. Soc. 2024, 146, 7313–23.

(29) Nguyen, D. T.; Zhu, L.; Gray, D. L.; Woods, T. J.; Padhi, C.; Flatt, K. M.; Mitchell, D. A.; van der Donk, W. A. Biosynthesis of macrocyclic peptides with C-terminal β-amino-α-keto acid groups by three different metalloenzymes. ACS Cent. Sci. 2024, 10, 1022–32.

(30) Zallot, R.; Oberg, N.; Gerlt, J. A. The EFI web resource for genomic enzymology tools: leveraging protein, genome, and metagenome databases to discover novel enzymes and metabolic pathways. Biochem. 2019, 58, 4169–82.

(31) Oberg, N.; Zallot, R.; Gerlt, J. A. EFI-EST, EFI-GNT, and EFI-CGFP: Enzyme function initiative (EFI) web resource for genomic enzymology tools. J. Mol. Biol. 2023, 435, 168018.

(32) Tietz, J. I.; Schwalen, C. J.; Patel, P. S.; Maxson, T.; Blair, P. M.; Tai, H. C.; Zakai, U. I.; Mitchell, D. A. A new genome-mining tool redefines the lasso peptide biosynthetic landscape. Nat. Chem. Biol. 2017, 13, 470–8.

(33) Burkhart, B. J.; Hudson, G. A.; Dunbar, K. L.; Mitchell, D. A. A prevalent peptide-binding domain guides ribosomal natural product biosynthesis. Nat. Chem. Biol. 2015, 11, 564–70.

(34) Zhang, Z.; Miller, W.; Schäffer, A. A.; Madden, T. L.; Lipman, D. J.; Koonin, E. V.; Altschul, S. F. Protein sequence similarity searches using patterns as seeds. Nucleic Acids Res. 1998, 26, 3986–90.

(35) Zhang, C.; Seyedsayamdost, M. R. Widespread peptide surfactants with post-translational C-methylations promote bacterial development. bioRxiv 2024, 2024.01.23.576971.

(36) Esmaeel, Q.; Ait Barka, E. Draft genome sequences of Bacillus strains isolated from the rhizosphere of maize cultivated in the northeast of France. 2018. University of Reims. Accession number RRZF00000000.1.

(37) Manetsberger, J.; Caballero Gómez, N.; Soria-Rodríguez, C.; Benomar, N.; Abriouel, H. Simply versatile: The use of Peribacillus simplex in sustainable agriculture. Microorganisms 2023, 11, 2540.

(38) Allioui, N.; Driss, F.; Dhouib, H.; Jlail, L.; Tounsi, S.; Frikha-Gargouri, O. Two novel Bacillus Strains (subtilis and simplex species) with promising potential for the biocontrol of Zymoseptoria tritici, the causal agent of septoria tritici blotch of wheat. Biomed Res. Int. 2021, 2021, 6611657.

(39) Manetsberger, J.; Caballero Gómez, N.; Benomar, N.; Christie, G.; Abriouel, H. Antimicrobial profile of the culturable olive sporobiota and its potential as a source of biocontrol agents for major phytopathogens in olive agriculture. Pest Manag. Sci. 2024, 80, 724–33.

(40) Gu, Y.-Q.; Mo, M.-H.; Zhou, J.-P.; Zou, C.-S.; Zhang, K.-Q. Evaluation and identification of potential organic nematicidal volatiles from soil bacteria. Soil Biol. Biochem. 2007, 39, 2567–75.

(41) Kloosterman, A. M.; Shelton, K. E.; Wezel, G. P. v.; Medema, M. H.; Mitchell, D. A. RRE-Finder: a genome-mining tool for class-independent RiPP discovery. mSystems 2020, 5, 10.1128/msystems.00267-20.

(42) Zheng, L.; Cash, V. L.; Flint, D. H.; Dean, D. R. Assembly of iron-sulfur clusters: Identification of an iscSUA-hscBA-fdx gene cluster from Azotobacter vinelandii. J. Biol. Chem. 1998, 273, 13264–72.

(43) Bassford, P. J.; Kadner, R. J. Genetic analysis of components involved in vitamin B12 uptake in Escherichia coli. J. Bacteriol. 1977, 132, 796–805.

(44) Lanz, N. D.; Blaszczyk, A. J.; McCarthy, E. L.; Wang, B.; Wang, R. X.; Jones, B. S.; Booker, S. J. Enhanced solubilization of class B radical S-adenosylmethionine methylases by improved cobalamin uptake in Escherichia coli. Biochem. 2018, 57, 1475–90.

(45) Yu, Y.; van der Donk, W. A. Biosynthesis of 3-thia-α-amino acids on a carrier peptide. Proc. Natl. Acad. Sci. U. S. A. 2022, 119, e2205285119.

(46) Harada, K.-i.; Fujii, K.; Hayashi, K.; Suzuki, M.; Ikai, Y.; Oka, H. Application of d,l-FDLA derivatization to determination of absolute configuration of constituent amino acids in peptide by advanced Marfey’s method. Tetrahedron Lett. 1996, 37, 3001–4.

(47) Abramson, J.; Adler, J.; Dunger, J.; Evans, R.; Green, T.; Pritzel, A.; Ronneberger, O.; Willmore, L.; Ballard, A. J.; Bambrick, J. et al. Accurate structure prediction of biomolecular interactions with AlphaFold 3. Nature 2024, 630, 493–500.

(48) Ortega, M. A.; Hao, Y.; Zhang, Q.; Walker, M. C.; van der Donk, W. A.; Nair, S. K. Structure and mechanism of the tRNA-dependent lantibiotic dehydratase NisB. Nature 2015, 517, 509–12.

(49) Koehnke, J.; Mann, G.; Bent, A. F.; Ludewig, H.; Shirran, S.; Botting, C.; Lebl, T.; Houssen, W. E.; Jaspars, M.; Naismith, J. H. Structural analysis of leader peptide binding enables leader-free cyanobactin processing. Nat. Chem. Biol. 2015, 11, 558–63.

(50) Evans, R. L., 3rd; Latham, J. A.; Xia, Y.; Klinman, J. P.; Wilmot, C. M. Nuclear magnetic resonance structure and binding studies of PqqD, a chaperone required in the biosynthesis of the bacterial dehydrogenase cofactor pyrroloquinoline quinone. Biochemistry 2017, 56, 2735–46.

(51) Grove, T. L.; Himes, P. M.; Hwang, S.; Yumerefendi, H.; Bonanno, J. B.; Kuhlman, B.; Almo, S. C.; Bowers, A. A. Structural insights into thioether bond formation in the biosynthesis of sactipeptides. J. Am. Chem. Soc. 2017, 139, 11734–44.

(52) Chekan, J. R.; Ongpipattanakul, C.; Nair, S. K. Steric complementarity directs sequence promiscuous leader binding in RiPP biosynthesis. Proc. Natl. Acad. Sci. U. S. A. 2019, 116, 24049–55.

(53) Sumida, T.; Dubiley, S.; Wilcox, B.; Severinov, K.; Tagami, S. Structural basis of leader peptide recognition in lasso peptide biosynthesis pathway. ACS Chem. Biol. 2019, 14, 1619–27.

(54) Ghilarov, D.; Stevenson, C. E. M.; Travin, D. Y.; Piskunova, J.; Serebryakova, M.; Maxwell, A.; Lawson, D. M.; Severinov, K. Architecture of microcin B17 synthetase: an octameric protein complex converting a ribosomally synthesized peptide into a DNA gyrase poison. Mol. Cell 2019, 73, 749–62.

(55) Shelton, K. E.; Mitchell, D. A. Bioinformatic prediction and experimental validation of RiPP recognition elements. Methods Enzymol. 2023, 679, 191–233.

(56) Gerlt, J. A.; Bouvier, J. T.; Davidson, D. B.; Imker, H. J.; Sadkhin, B.; Slater, D. R.; Whalen, K. L. Enzyme function initiative-enzyme similarity tool (EFI-EST): a web tool for generating protein sequence similarity networks. Biochim. Biophys. Acta, Proteins Proteomics 2015, 1854, 1019–37.

(57) Atkinson, H. J.; Morris, J. H.; Ferrin, T. E.; Babbitt, P. C. Using sequence similarity networks for visualization of relationships across diverse protein superfamilies. PLoS One 2009, 4, e4345.

(58) Fitzpatrick, P. F. Mechanism of aromatic amino acid hydroxylation. Biochemistry 2003, 42, 14083–91.

(59) Mitchell, K. H.; Rogge, C. E.; Gierahn, T.; Fox, B. G. Insight into the mechanism of aromatic hydroxylation by toluene 4-monooxygenase by use of specifically deuterated toluene and p-xylene. Proc. Natl. Acad. Sci. U. S. A. 2003, 100, 3784–9.

(60) Ullrich, R.; Hofrichter, M. Enzymatic hydroxylation of aromatic compounds. Cell. Mol. Life Sci. 2007, 64, 271–93.

(61) de Visser, S. P. In Adv. Inorg. Chem.; Eldik, R. v., Ivanović-Burmazović, I., Eds.; Academic Press: 2012; Vol. 64, p 1–31.

(62) Shen, X.; Zhou, D.; Lin, Y.; Wang, J.; Gao, S.; Kandavelu, P.; Zhang, H.; Zhang, R.; Wang, B.-C.; Rose, J. et al. Structural insights into catalytic versatility of the flavin-dependent hydroxylase (HpaB) from Escherichia coli. Sci. Rep. 2019, 9, 7087.

(63) Frattaruolo, L.; Lacret, R.; Cappello, A. R.; Truman, A. W. A Genomics-Based Approach Identifies a Thioviridamide-Like Compound with Selective Anticancer Activity. ACS Chem. Biol. 2017, 12, 2815–22.

(64) Guroff, G.; Daly, J. W.; Jerina, D. M.; Renson, J.; Witkop, B.; Udenfriend, S. Hydroxylation-induced migration: the NIH shift. Recent experiments reveal an unexpected and general result of enzymatic hydroxylation of aromatic compounds. Science 1967, 157, 1524–30.

(65) Debono, M.; Abbott, B. J.; Molloy, R. M.; Fukuda, D. S.; Hunt, A. H.; Daupert, V. M.; Counter, F. T.; Ott, J. L.; Carrell, C. B.; Howard, L. C. et al. Enzymatic and chemical modifications of lipopeptide antibiotic A21978C: the synthesis and evaluation of daptomycin (LY146032). J. Antibiot. 1988, 41, 1093–105.

(66) Synge, R. L. M. ‘Gramicidin S’: over-all chemical characteristics and amino-acid composition. Biochem. J. 1945, 39, 363–7.

(67) Mordhorst, S.; Badmann, T.; Bösch, N. M.; Morinaka, B. I.; Rauch, H.; Piel, J.; Groll, M.; Vagstad, A. L. Structural and biochemical insights into post-translational arginine-to-ornithine peptide modifications by an atypical arginase. ACS Chem. Biol. 2023, 18, 528–36.

(68) Mordhorst, S.; Morinaka, B. I.; Vagstad, A. L.; Piel, J. Posttranslationally acting arginases provide a ribosomal route to non-proteinogenic ornithine residues in diverse peptide sequences. Angew. Chem. Int. Ed. 2020, 59, 21442–7.

(69) Hug, J. J.; Dastbaz, J.; Adam, S.; Revermann, O.; Koehnke, J.; Krug, D.; Müller, R. Biosynthesis of cittilins, unusual ribosomally synthesized and post-translationally modified peptides from Myxococcus xanthus. ACS Chem. Biol. 2020, 15, 2221–31.

(70) Molinaro, C.; Kawasaki, Y.; Wanyoike, G.; Nishioka, T.; Yamamoto, T.; Snedecor, B.; Robinson, S. J.; Gosselin, F. Engineered cytochrome P450-catalyzed oxidative biaryl coupling reaction provides a scalable entry into arylomycin antibiotics. J. Am. Chem. Soc. 2022, 144, 14838–45.

(71) Zdouc, M. M.; Iorio, M.; Vind, K.; Simone, M.; Serina, S.; Brunati, C.; Monciardini, P.; Tocchetti, A.; Zarazúa, G. S.; Crüsemann, M. et al. Effective approaches to discover new microbial metabolites in a large strain library. J. Ind. Microbiol. Biotechnol. 2021, 48.

(72) Zdouc, M. M.; Alanjary, M. M.; Zarazúa, G. S.; Maffioli, S. I.; Crüsemann, M.; Medema, M. H.; Donadio, S.; Sosio, M. A biaryl-linked tripeptide from Planomonospora reveals a widespread class of minimal RiPP gene clusters. Cell Chem. Biol 2021, 28, 733–9.

(73) Padva, L.; Gullick, J.; Coe, L. J.; Hansen, M. H.; De Voss, J. J.; Crüsemann, M.; Cryle, M. J. The biarylitides: Understanding the structure and biosynthesis of a fascinating class of cytochrome P450 modified RiPP natural products. ChemBioChem 2024, e202400916.

(74) Hug, J. J.; Frank, N. A.; Walt, C.; Šenica, P.; Panter, F.; Müller, R. Genome-guided discovery of the first myxobacterial biarylitide myxarylin reveals distinct C-N biaryl crosslinking in RiPP biosynthesis. Molecules 2021, 26.

(75) Schramma, K. R.; Forneris, C. C.; Caruso, A.; Seyedsayamdost, M. R. Mechanistic investigations of lysine-tryptophan cross-link formation catalyzed by streptococcal radical S-adenosylmethionine enzymes. Biochemistry 2018, 57, 461–8.

(76) Haslinger, K.; Peschke, M.; Brieke, C.; Maximowitsch, E.; Cryle, M. J. X-domain of peptide synthetases recruits oxygenases crucial for glycopeptide biosynthesis. Nature 2015, 521, 105–9.

(77) Sugiyama, R.; Suarez, A. F. L.; Morishita, Y.; Nguyen, T. Q. N.; Tooh, Y. W.; Roslan, M. N. H. B.; Lo Choy, J.; Su, Q.; Goh, W. Y.; Gunawan, G. A. et al. The biosynthetic landscape of triceptides reveals radical SAM enzymes that catalyze cyclophane formation on Tyr- and His-containing motifs. J. Am. Chem. Soc. 2022, 144, 11580–93.

(78) Phan, C.-S.; Morinaka, B. I. A prevalent group of Actinobacterial radical SAM/SPASM maturases involved in triceptide biosynthesis. ACS Chem. Biol. 2022, 17, 3284–9.

(79) Balo, A. R.; Caruso, A.; Tao, L.; Tantillo, D. J.; Seyedsayamdost, M. R.; Britt, R. D. Trapping a cross-linked lysine–tryptophan radical in the catalytic cycle of the radical SAM enzyme SuiB. Proc. Natl. Acad. Sci. U.S.A. 2021, 118, e2101571118.

(80) Schramma, K. R.; Forneris, C. C.; Caruso, A.; Seyedsayamdost, M. R. Mechanistic investigations of lysine–tryptophan cross-link formation catalyzed by Streptococcal radical S-adenosylmethionine enzymes. Biochem. 2018, 57, 461–8.

(81) Zhang, Q.; van der Donk, W. A.; Liu, W. Radical-mediated enzymatic methylation: a tale of two SAMs. Acc. Chem. Res. 2012, 45, 555–64.

(82) Bridwell-Rabb, J.; Li, B.; Drennan, C. L. Cobalamin-dependent radical S-adenosylmethionine enzymes: capitalizing on old motifs for new functions. ACS Bio. & Med. Chem. Au. 2022, 2, 173–86.

(83) Bridwell-Rabb, J.; Zhong, A.; Sun, H. G.; Drennan, C. L.; Liu, H.-w. A B12-dependent radical SAM enzyme involved in oxetanocin A biosynthesis. Nature 2017, 544, 322–6.

(84) Vobruba, S.; Kadlcik, S.; Janata, J.; Kamenik, Z. TldD/TldE peptidases and N-deacetylases: A structurally unique yet ubiquitous protein family in the microbial metabolism. Microbiol. Res. 2022, 265, 127186.

(85) Allali, N.; Afif, H.; Couturier, M.; Van Melderen, L. The highly conserved TldD and TldE proteins of Escherichia coli are involved in microcin B17 processing and in CcdA degradation. J. Bacteriol. 2002, 184, 3224–31.

(86) Ghilarov, D.; Serebryakova, M.; Stevenson, C. E. M.; Hearnshaw, S. J.; Volkov, D. S.; Maxwell, A.; Lawson, D. M.; Severinov, K. The origins of specificity in the microcin-processing protease TldD/E. Structure 2017, 25, 1549-61.e5.

(87) Phan, C. S.; Morinaka, B. I. A prevalent group of Actinobacterial radical SAM/SPASM maturases involved in triceptide biosynthesis. ACS Chem. Biol. 2022, 17, 3284–9.

(88) Sugiyama, R.; Suarez, A. F. L.; Morishita, Y.; Nguyen, T. Q. N.; Tooh, Y. W.; Roslan, M.; Lo Choy, J.; Su, Q.; Goh, W. Y.; Gunawan, G. A. et al. The biosynthetic landscape of triceptides reveals radical SAM enzymes that catalyze cyclophane formation on Tyr- and His-containing motifs. J. Am. Chem. Soc. 2022, 144, 11580–93.

(89) Zhao, Y.; Marschall, E.; Treisman, M.; McKay, A.; Padva, L.; Crüsemann, M.; Nelson, D. R.; Steer, D. L.; Schittenhelm, R. B.; Tailhades, J. et al. Cytochrome P450(Blt) enables versatile peptide cyclisation to generate histidine- and tyrosine-containing crosslinked tripeptide building blocks. Angew. Chem. Int. Ed. 2022, 61, e202204957.

(90) Coe, L. J.; Zhao, Y.; Padva, L.; Keto, A.; Schittenhelm, R.; Tailhades, J.; Pierens, G.; Krenske, E. H.; Crüsemann, M.; De Voss, J. J. et al. Reassignment of the structure of a tryptophan-containing cyclic tripeptide produced by the biarylitide crosslinking cytochrome P450(blt). Chemistry 2024, 30, e202400988.

(91) Hansen, M. H.; Keto, A.; Treisman, M.; Sasi, V. M.; Coe, L.; Zhao, Y.; Padva, L.; Hess, C.; Leichthammer, V.; Machell, D. L. et al. Structural insights into a side chain cross-linking biarylitide P450 from RiPP biosynthesis. ACS Catal. 2024, 14, 812–26.

(92) Miller, R. D.; Iinishi, A.; Modaresi, S. M.; Yoo, B. K.; Curtis, T. D.; Lariviere, P. J.; Liang, L.; Son, S.; Nicolau, S.; Bargabos, R. et al. Computational identification of a systemic antibiotic for gram-negative bacteria. Nat. Microbiol. 2022, 7, 1661–72.

(93) Lewis, K.; Lee, R. E.; Brötz-Oesterhelt, H.; Hiller, S.; Rodnina, M. V.; Schneider, T.; Weingarth, M.; Wohlgemuth, I. Sophisticated natural products as antibiotics. Nature 2024, 632, 39–49.

(94) Liu, J.; Liu, R.; He, B. B.; Lin, X.; Guo, L.; Wu, G.; Li, Y. X. Bacterial cytochrome P450 catalyzed macrocyclization of ribosomal peptides. ACS Bio. Med. Chem. Au 2024, 4, 268–79.

(95) Hu, Y. L.; Yin, F. Z.; Shi, J.; Ma, S. Y.; Wang, Z. R.; Tan, R. X.; Jiao, R. H.; Ge, H. M. P450-modified ribosomally synthesized peptides with aromatic cross-links. J. Am. Chem. Soc. 2023, 145, 27325–35.

(96) Clark, K. A.; Bushin, L. B.; Seyedsayamdost, M. R. RaS-RiPPs in streptococci and the human microbiome. ACS Bio. Med. Chem. Au 2022, 2, 328–39.

(97) Kaur, H.; Jakob, R. P.; Marzinek, J. K.; Green, R.; Imai, Y.; Bolla, J. R.; Agustoni, E.; Robinson, C. V.; Bond, P. J.; Lewis, K. et al. The antibiotic darobactin mimics a β-strand to inhibit outer membrane insertase. Nature 2021, 593, 125–9.

