## Supporting Information for "Biosynthesis of Macrocyclic Peptides by Formation and Crosslinking of *ortho*-Tyrosines"

[1] Department of Chemistry and Howard Hughes Medical Institute, University of Illinois at Urbana−Champaign, Urbana, Illinois 61801, USA.

[2] Carl R. Woese Institute for Genomic Biology, University of Illinois at Urbana- Champaign, Urbana, Illinois, 61801, USA.

[3] School of Chemical Sciences NMR Laboratory, University of Illinois at Urbana-Champaign, Urbana, 61801, IL, USA.

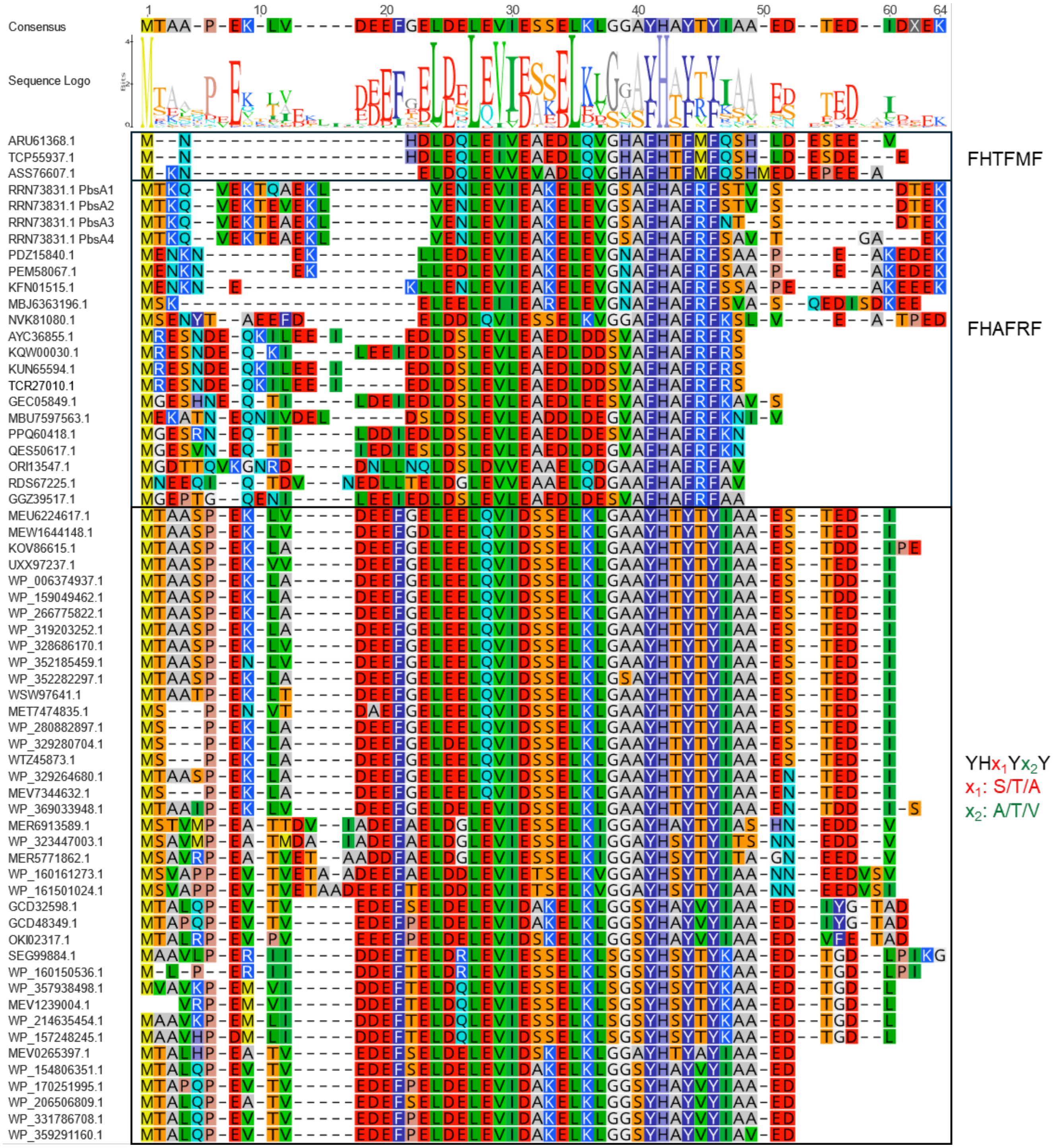


**Figure S1: Multiple sequence alignment of FHAFRF motif containing putative precursors and orthologous peptide sequences extracted through pattern hit initiated (PHI) BLAST.**

A

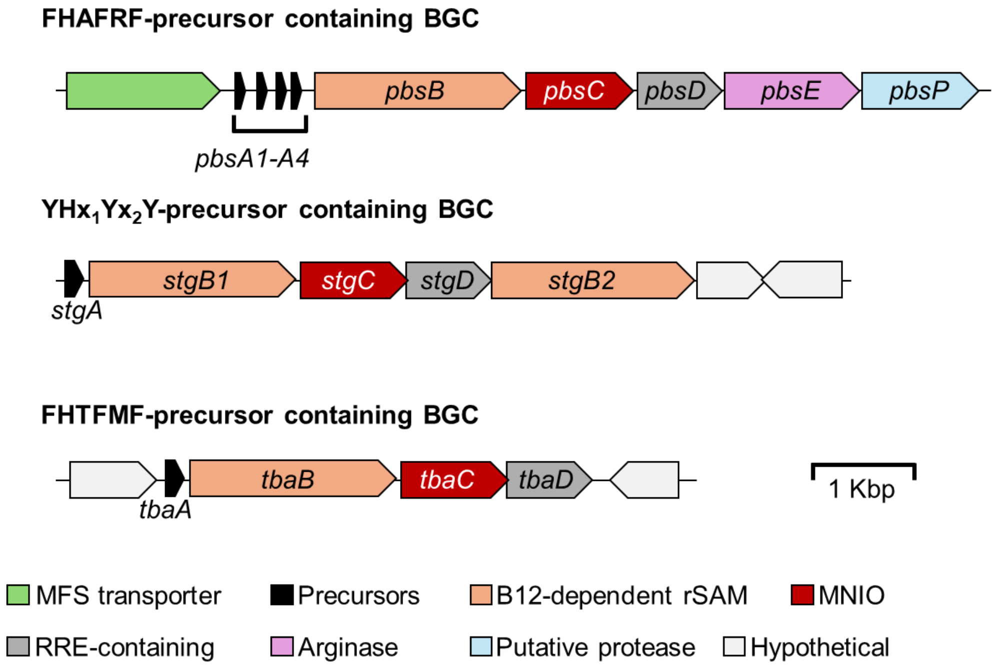


B

| | |
|---|---|
| PbsA1 | : MTKQVEKTQAEKLVENLEVIEAKELEVGSAFHAFRFSTVSDTEK |
| PbsA2 | : MTKQVEKTEVEKLVENLEVIEAKELEVGSAFHAFRFSTVSDTEK |
| PbsA3 | : MTKQVEKTEAEKLVENLEVIEAKELEVGSAFHAFRFNTSDTEK |
| PbsA4 | : MTKQVEKTEAEKLVENLEVIEAKELEVGSAFHAFRFSAVTGAEK |
| StgA | : MTAASPEKLADEEFGELEELQVIDSSELKLGAAYHTYTYIAAESTDDIPD |
| TbaA | : MNHDLDQLEIVEAEDLQVGHAFHTFMFQSHLDESEEV |

**Figure S2: Genome neighborhood architecture of *pbs* cluster and orthologous BGCs.**
(A) Genome neighborhood architecture of BGCs containing FHAFRF motif-containing precursors, $YHx_1Yx_2Y$ motif-containing precursors, where $x_1$ = S/T/A and $x_2$ = T/V/A, and FHTFMF motif-containing precursors. (B) Precursor peptide sequences of PbsA1-A4, StgA and TbaA.

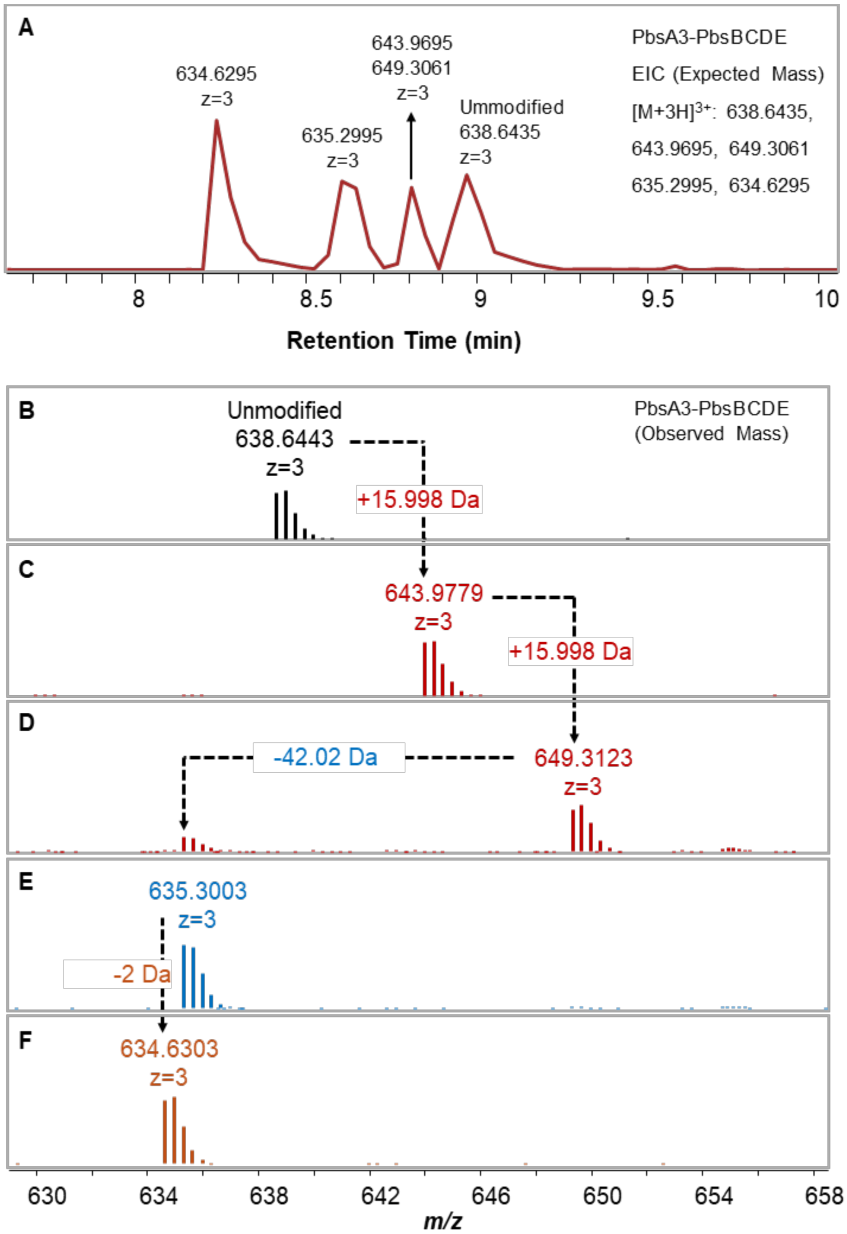


**Figure S3: LC-MS analysis of GluC-digested PbsA3 co-expressed with PbsBCDE. (A) Extracted ion chromatogram (EIC) of GluC-digested products.** The mixture contained unmodified PbsA3 (expected $[M+3H]^{3+}$ = 638.6429 Da), intermediates undergoing 15.99 Da and 31.99 Da mass gains, characteristic of mono- and bis-hydroxylations (expected $[M+3H]^{3+}$ = 643.9695 Da and 649.3061 Da, respectively), putative deguanidinated and bis-hydroxylated intermediate (expected $[M+3H]^{3+}$ = 635.2995 Da), and a putative deguanidinated, bis-hydroxylated and cross-linked product (expected $[M+3H]^{3+}$ = 634.6295 Da). (B) HR-MS spectra of the extracted ions showing the isotopic peak distribution for the unmodified PbsA3 (observed $[M+3H]^{3+}$ = 638.6443 Da), putative mono- and bis-hydroxylated intermediates (observed $[M+3H]^{3+}$ = 643.9779 Da and 649.3123 Da, respectively), putative deguanidinated and bis-hydroxylated intermediate (observed $[M+3H]^{3+}$ = 635.3003 Da), and the putative deguanidinated, bis-hydroxylated and cross-linked product (observed $[M+3H]^{3+}$ = 634.6303 Da).

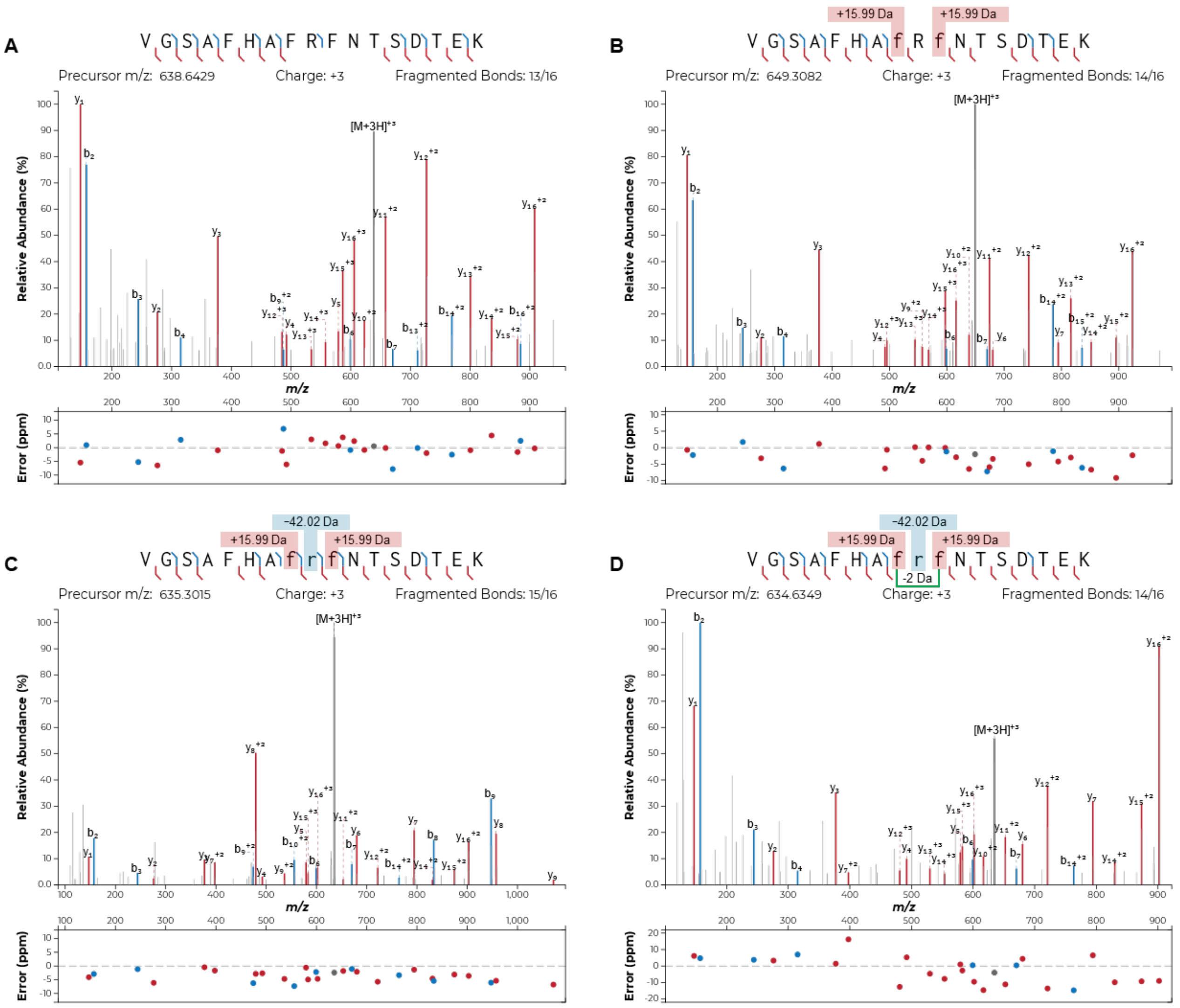


**Figure S4: HR-MS/MS analysis of GluC-digested PbsA3 co-expressed with PbsBCDE.**
HR-MS/MS fragmentation of (A) unmodified PbsA3, (B) putative bis-hydroxylated intermediate, PbsA3-CD showing the characteristic 15.99 Da and 31.99 Da mass gains on the Phe8 and Phe10 residues, (C) putative deguanidinated and bis-hydroxylated intermediate, PbsA3-CDE showing an additional mass loss of 42.02 Da on the R9 residue, and (D) the putative deguanidinated, bis-hydroxylated and cross-linked product displaying an additional 2 Da mass loss in the FRF region. No fragmentation in the FRF region was indicative of macrocyclization. Residues with mass shifts are denoted in lower case letters. Hypothesized crosslinking residues are connected by a green bracket. Plots were prepared using the Interactive Peptide Annotator Webtool.[1]

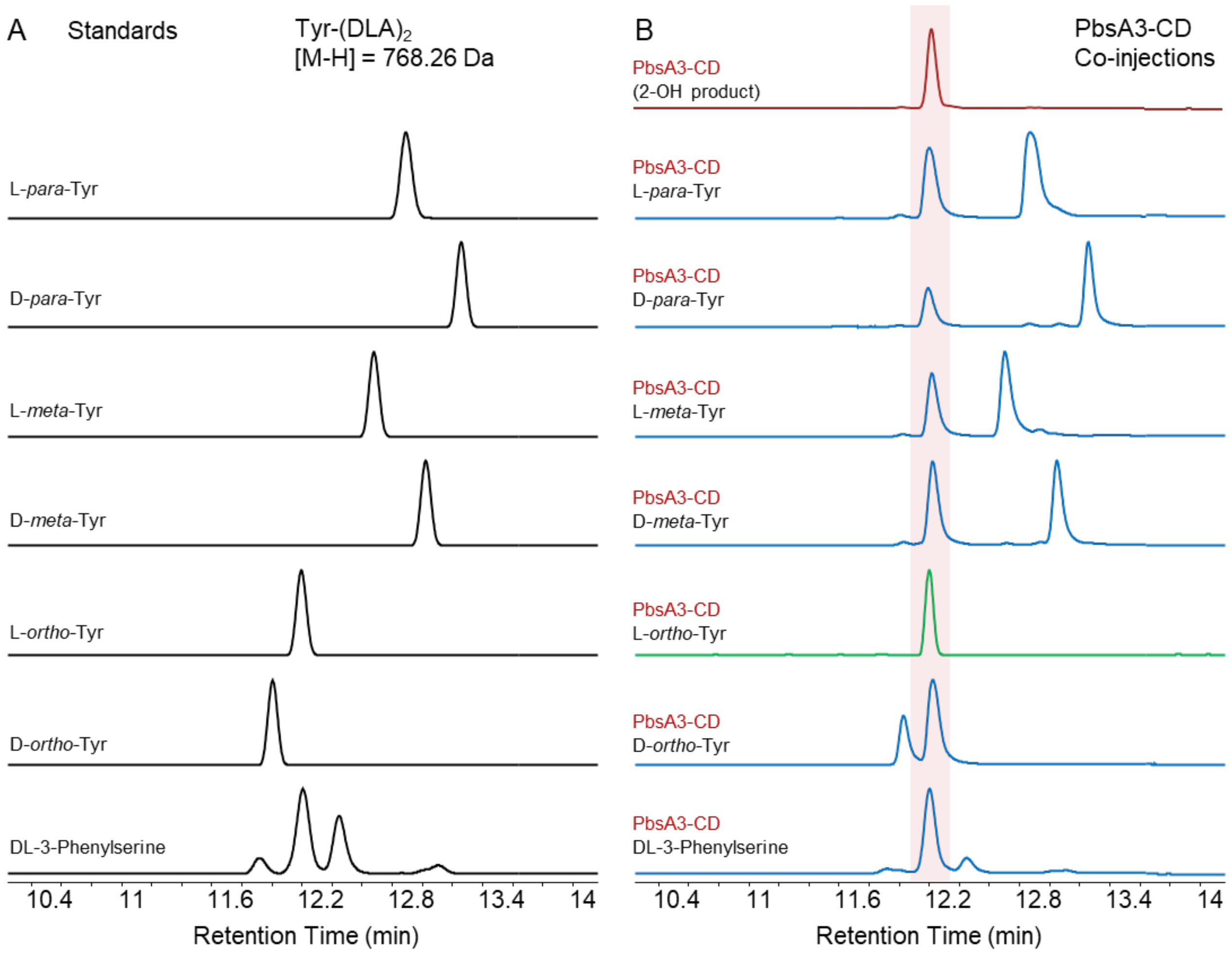


**Figure S5: Advanced Marfey's analysis of PbsA3-CD product.**

(A) Extracted ion chromatograms (EIC) of Tyr analogues that were bis-derivatized with L-FDLA (Tyr-$(DLA)_2$) with a calculated mass of [M-H] = 768.26 Da. (B) HPLC-purified GluC-digested PbsA3-CD product hydrolyzed with HCl and derivatized with L-FDLA (red). Derivatized PbsA3-CD product was co-injected with the standards from panel A (blue and green). Retention time of the EIC of PbsA3-CD coincided with that of L-*ortho*-Tyr (green chromatogram) and one of the stereoisomers of DL-3-phenylserine suggesting the hydroxylations to be either on the *ortho*-position or on the *beta*-carbon of the Phe8/Phe10 residues.

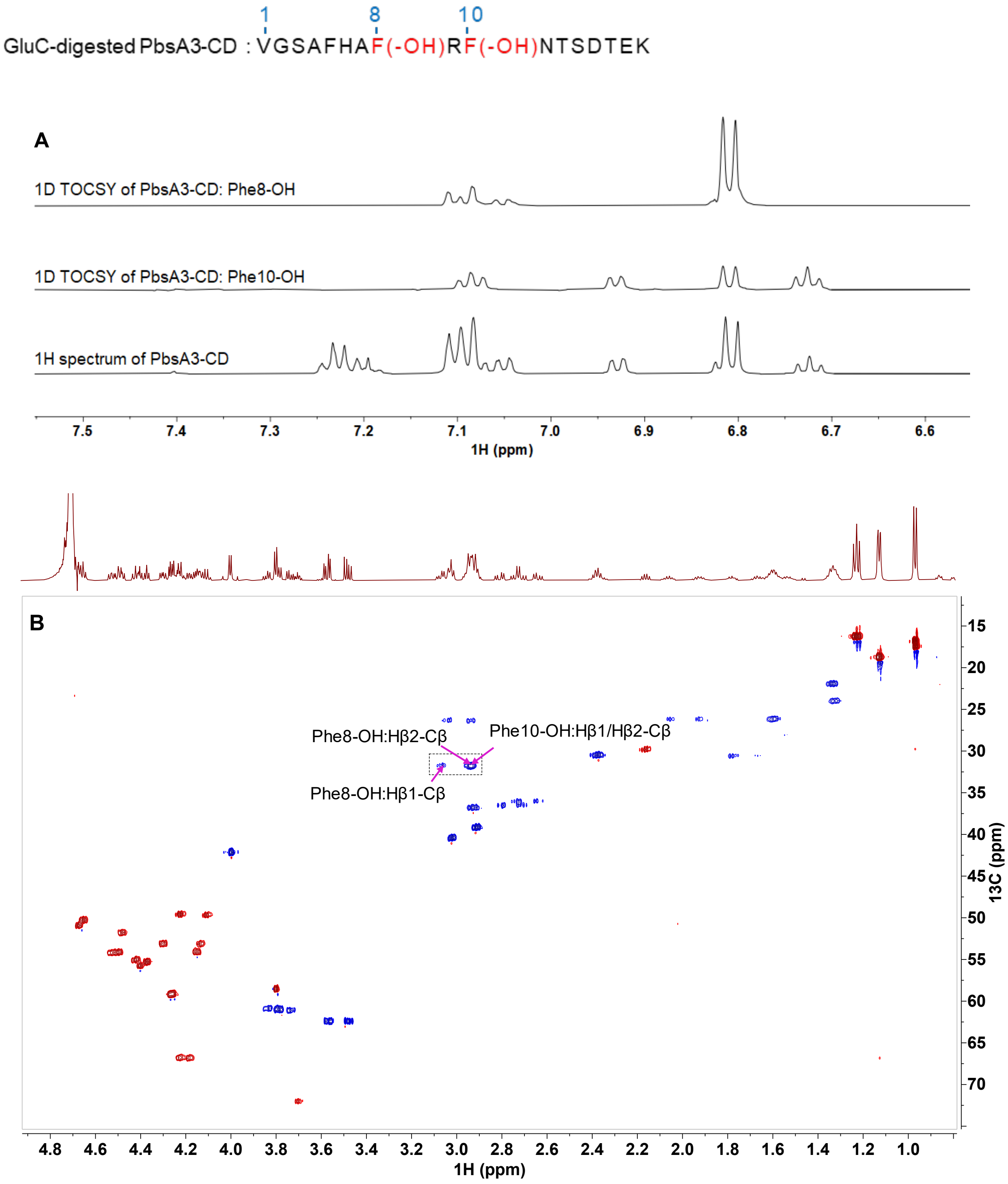


**Figure S6: 1D $^{1}H$-$^{1}H$ TOCSY and the aliphatic region of $^{1}H$-$^{13}C$ HSQC spectra of GluC-digested PbsA3-CD in 100%$D_2O$.** (A) 1D $^{1}H$ TOCSY spectra of GluC-digested PbsA3-CD show four aromatic protons for each oxidized Phe (Phe8-OH and Phe10-OH) with 2 doublets and 2 triplets peaks. In the Phe8-OH spectrum (top), peak at ~6.8 ppm integrated to 2 protons, one doublet and one triplet, which was confirmed by 2D $^{1}H$-$^{1}H$ TOCSY, $^{1}H$-$^{13}C$ HSQC and HMBC, as well as 1D $^{1}H$-$^{1}H$ TOCSY under different mixing times. (B) The aliphatic region of the 2D $^{1}H$-$^{13}C$ multiplicity edited HSQC spectrum. The CH and $CH_3$ peaks are shown in red, and $CH_2$ peaks are shown in blue. The Hβ-Cβ cross peaks for Phe8-OH and Phe10-OH are labeled. Both spectra eliminate the possibility of hydroxylation at the beta-carbon in addition to the information provided by $^{1}H$-$^{13}C$ HMBC (Fig 3A and 3B). Sequence of the analyzed peptide is shown at the top. Modified residues are indicated in brackets.

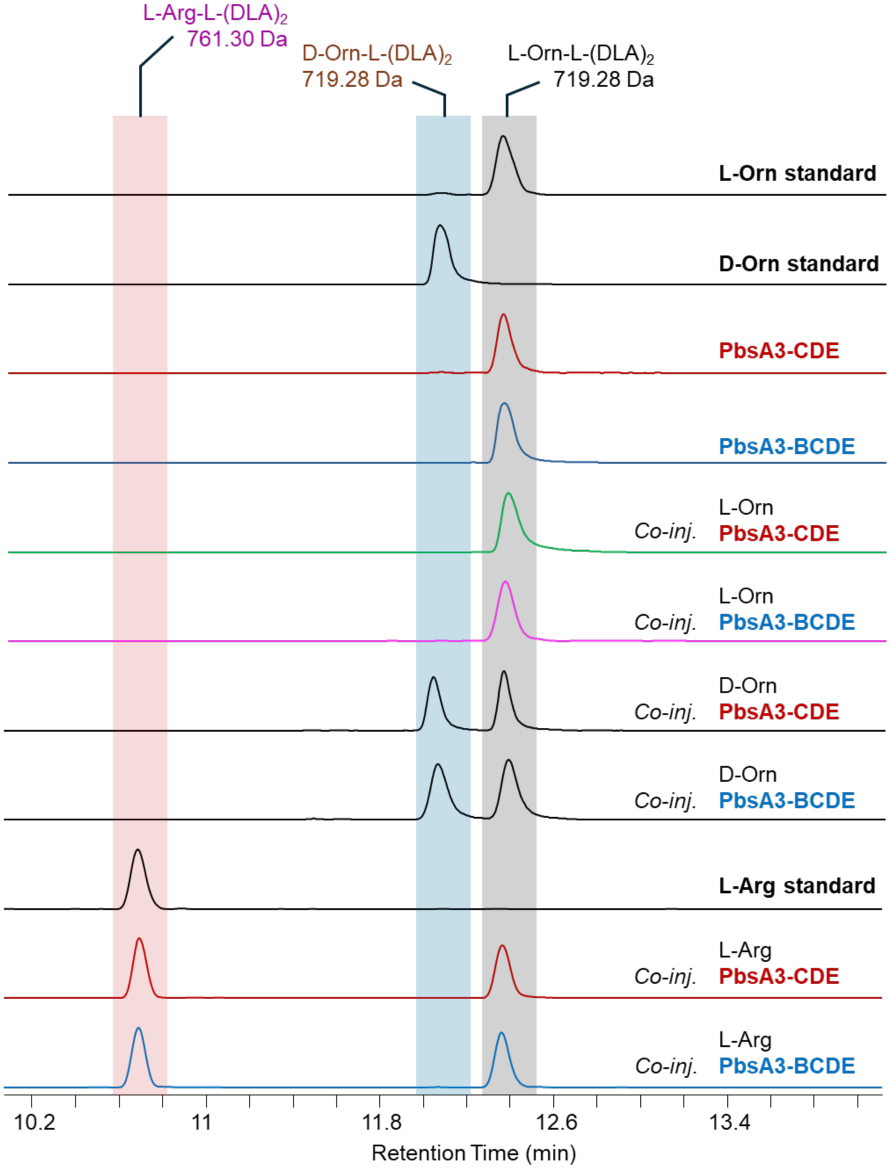


**Figure S7: Advanced Marfey's analysis of PbsA3-CDE product.** Extracted ion chromatograms (EIC) of Orn and Arg analogs bis-derivatized with L-FDLA (DL-Orn-L-$(DLA)_2$ and L-Arg-L-$(DLA)_2$) with a calculated mass of [M-H] = 719.28 Da and 761.30 Da, respectively. HPLC-purified GluC-digested PbsA3-CDE and PbsA3-BCDE products were hydrolyzed with DCl and derivatized with L-FDLA. EICs of the bis-derivatized PbsA3-CDE and -BCDE products (red and blue, respectively) are shown as well as co-injection with the Orn and Arg standards. The retention time of the analytes coincided with that of L-Orn (green and pink chromatograms)**.**

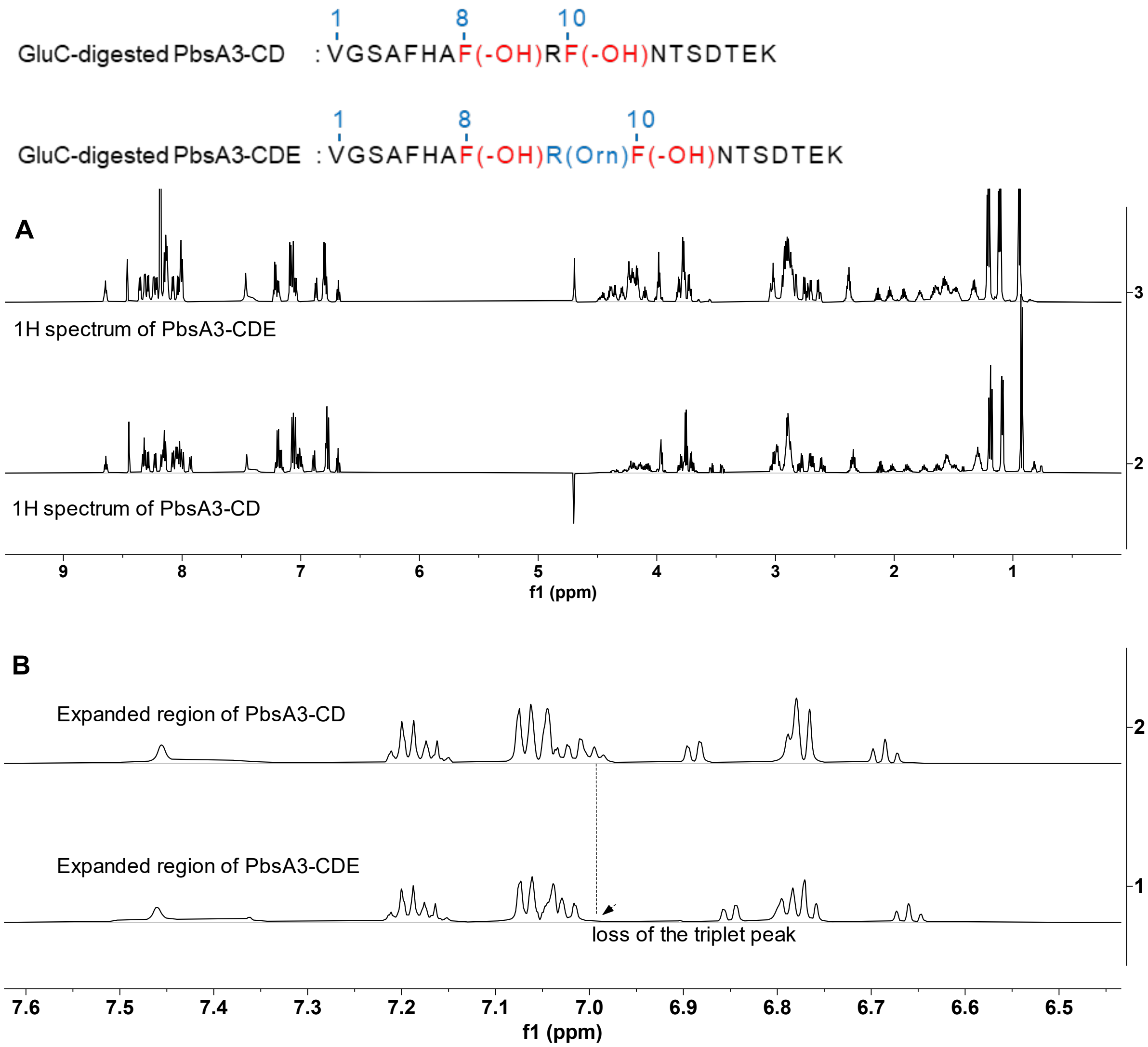


**Figure S8: $^1$H NMR spectra of GluC-digested PbsA3-CD and PbsA3-CDE in 90% $H_2O$ and 10% $D_2O$.** (A) Full range $^1$H spectrum of GluC-digested PbsA3-CDE compared with that of PbsA3-CD (also GluC-digested). (B) Expanded aromatic region of the $^1$H spectrum of PbsA3-CDE compared with that of PbsA3-CD, which shows the loss of the triplet NHε peak of residue 9 in PbsA3-CDE. Sequence of the analyzed peptide is shown at the top. Modified residues are indicated in brackets.

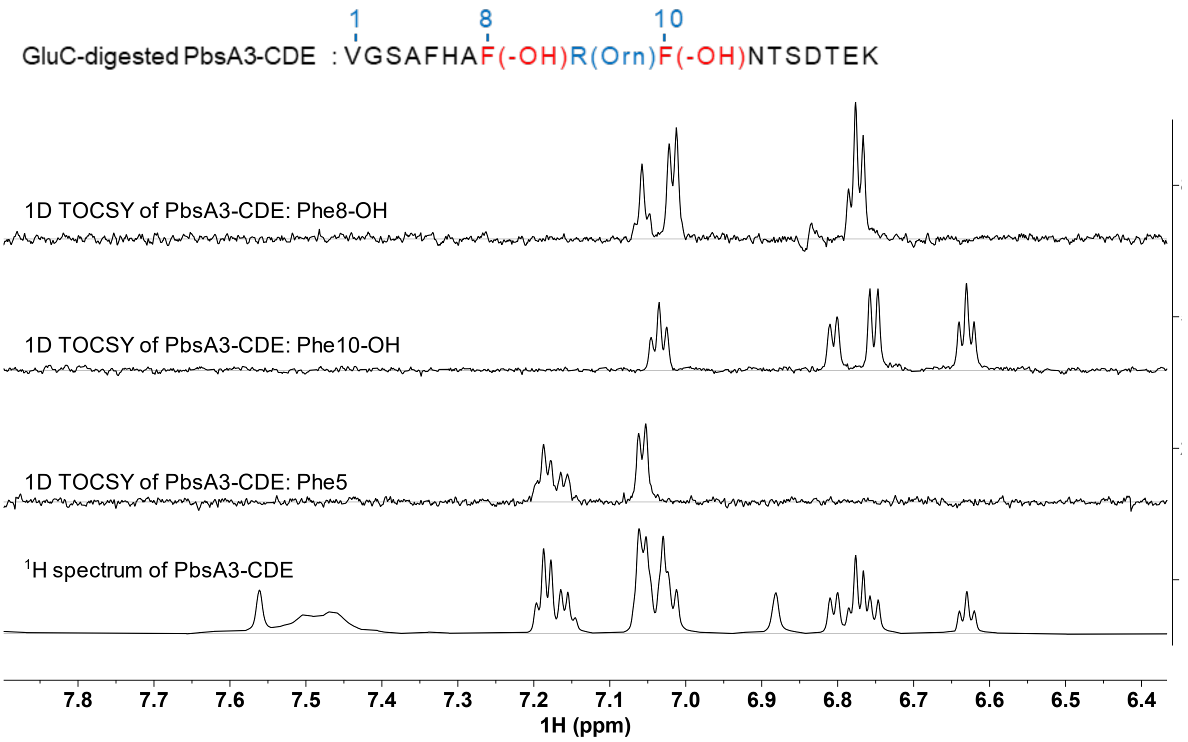


**Figure S9: 1D $^{1}$H-$^{1}$H TOCSY spectra of GluC-digested PbsA3-CDE in 90% $H_2O$ and 10% $D_2O$ at 3 °C**. Four aromatic protons were observed for Phe8-OH and Phe10-OH containing 2 triplets and 2 doublets at each ring, whereas Phe5 showed 5 protons with the normal splitting pattern. In the top spectrum of Phe8-OH, the peak around 6.8 ppm integrated to 2 protons, one doublet and one triplet which was confirmed by 2D $^{1}$H-$^{1}$H TOCSY, $^{1}$H-$^{13}$C HSQC and HMBC, as well as 1D $^{1}$H-$^{1}$H TOCSY at different mixing time. Sequence of the analyzed peptide is shown at the top. Modified residues are indicated in brackets.

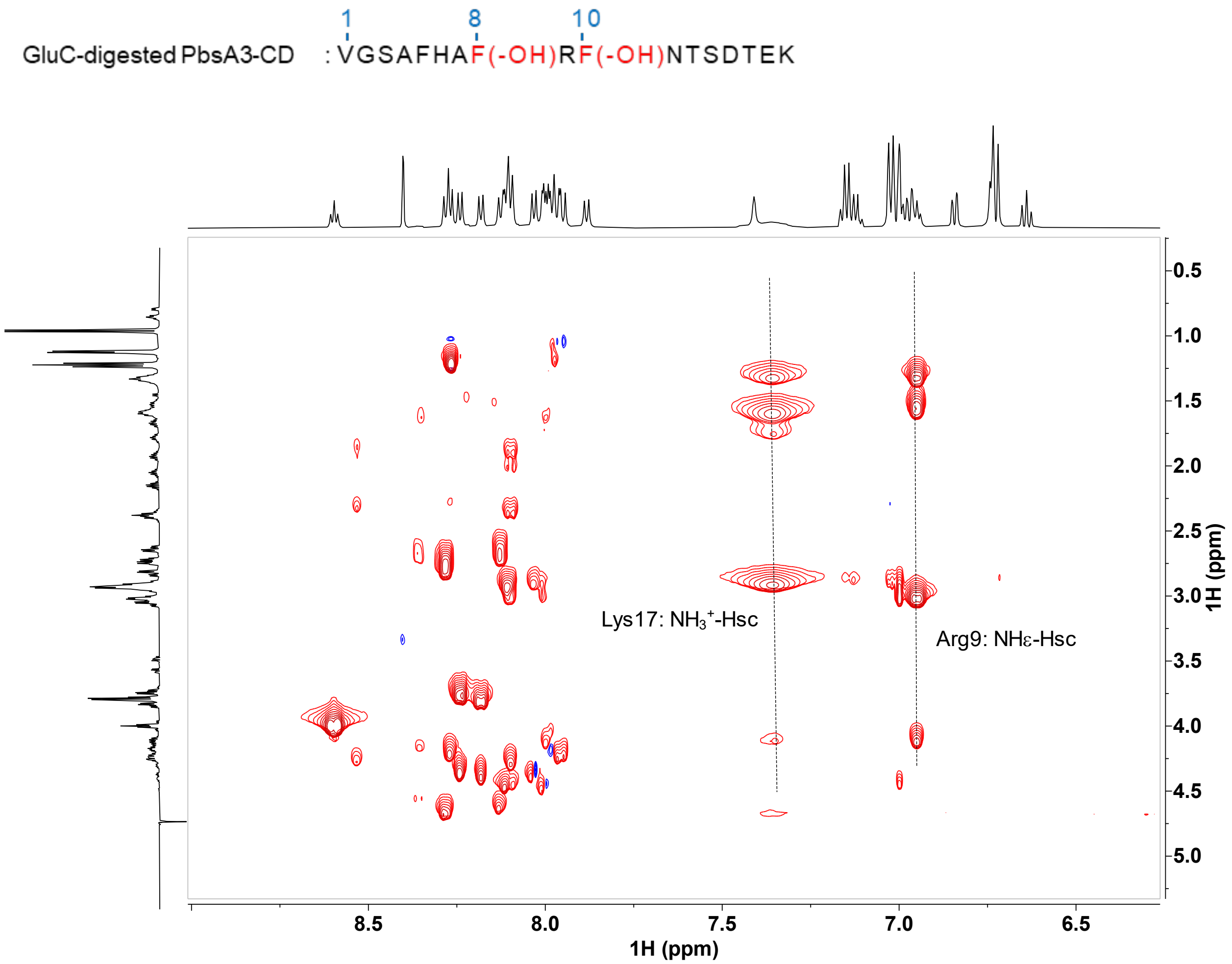


**Figure S10: The aromatic region of the 2D $^{1}H$-$^{1}H$ TOCSY spectra of PbsA3-CD in 90% $H_2O$ and 10% $D_2O$ at 25 ºC.** The NHε triplet of Arg9 showed cross peaks with the other side chain protons (labeled as $H_{sc}$ in the figure) and the amide proton of Arg9. Sequence of the analyzed peptide is shown at the top. Modified residues are indicated in brackets.

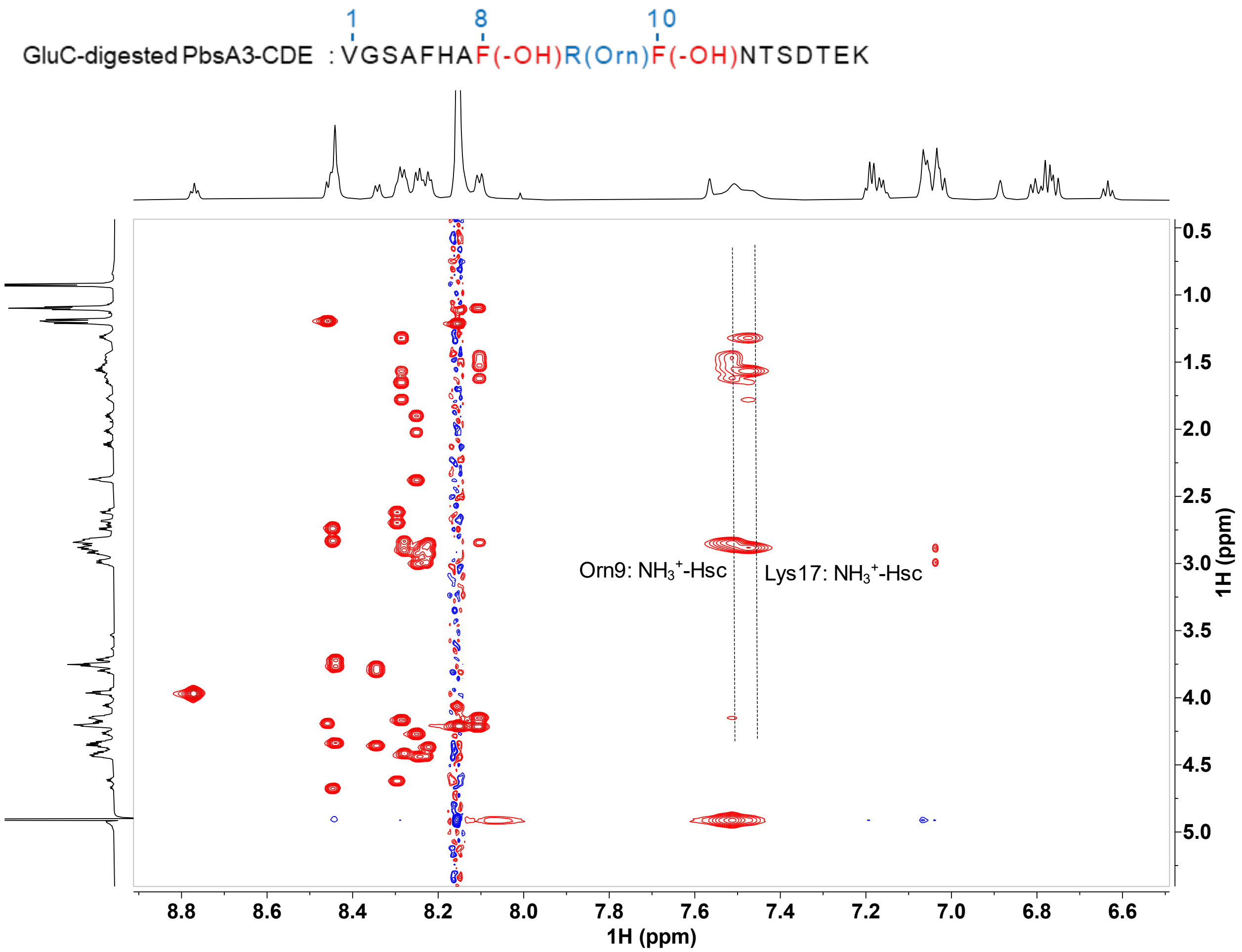


**Figure S11: The aromatic region of the 2D $^1H$-$^1H$ TOCSY spectra of GluC-digested PbsA3-CDE in 90% $H_2O$ and 10% $D_2O$ containing 0.1% formic acid-$d_2$ at 3 ºC.** The triplet NHε of Arg9 in PbsA3-CD was no longer present. Instead, a broad peak, which integrates to three protons, showed cross peak with the other side chain protons (labeled as $H_{sc}$ in the figure) of Arg9, consistent with the transformation of Arg9 to Orn9. Sequence of the analyzed peptide is shown at the top. Modified residues are indicated in brackets.

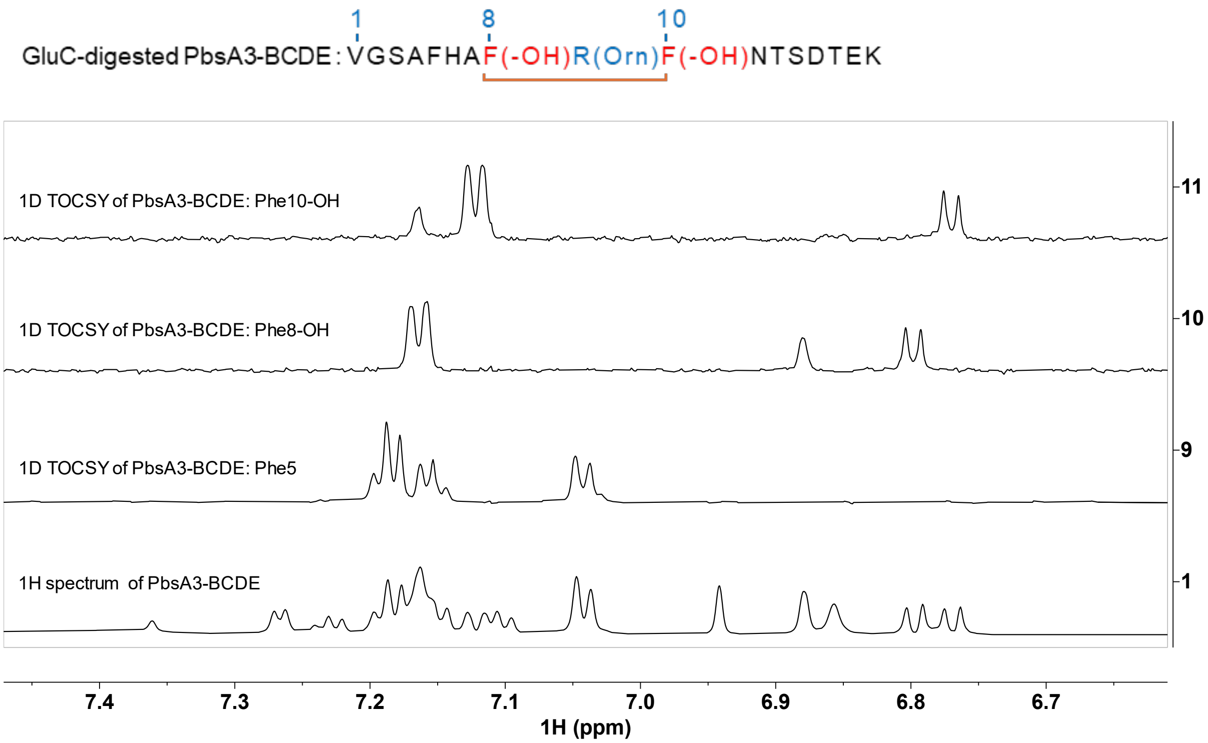


**Figure S12: 1D $^1$H-$^1$H TOCSY spectra of GluC-digested PbsA3-BCDE in 90% $H_2O$ and 10% $D_2O$ containing 0.1% formic acid-$d_2$.**

The Phe8-OH and Phe10-OH showed 3 protons each in their aromatic ring. Phe5 remained unchanged. Sequence of the analyzed peptide is shown at the top. Modified residues are indicated in brackets. Crosslinked residues are connected (in orange).

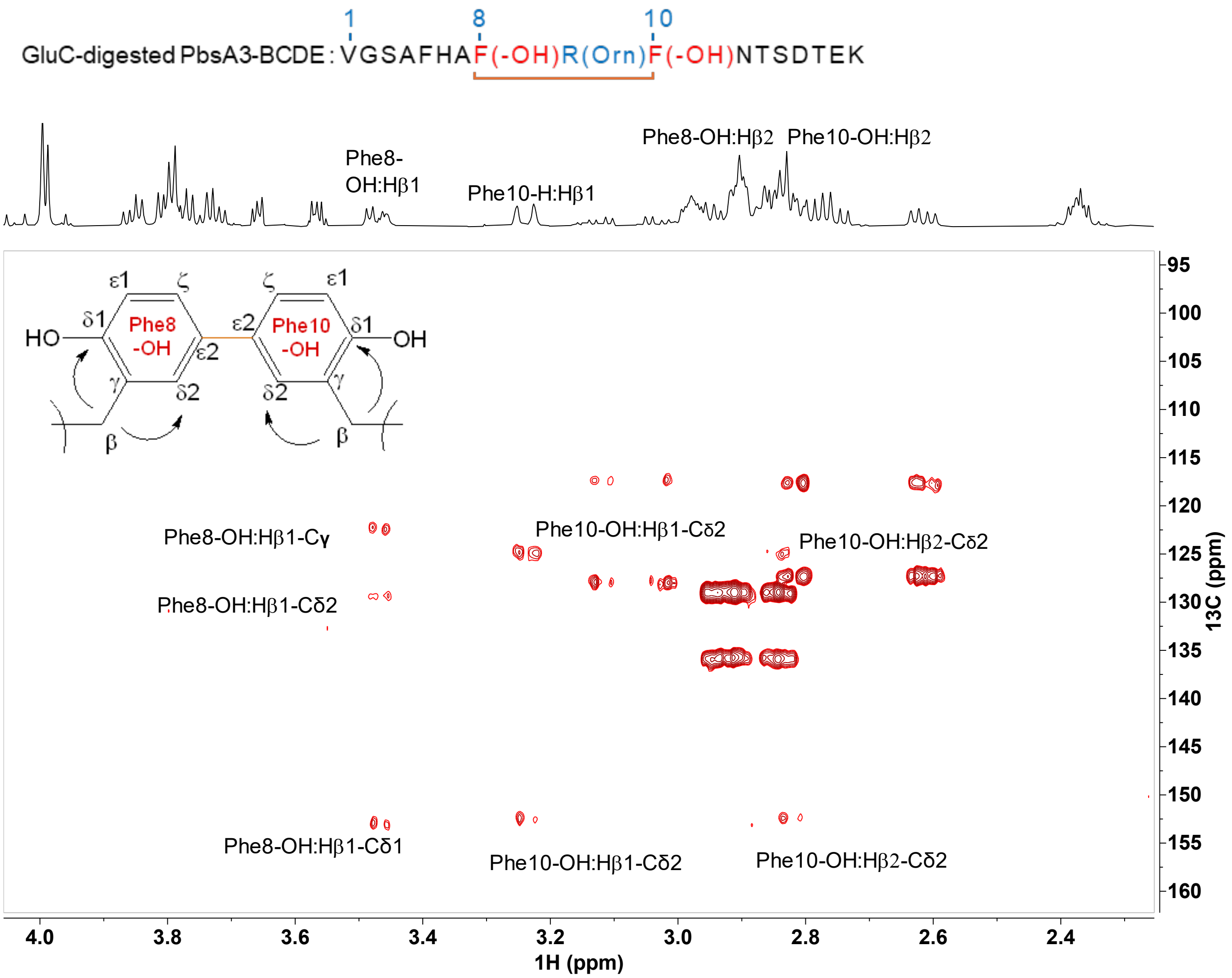


**Figure S13: The aliphatic region of a $^{1}H$-$^{13}C$ HSQC spectrum of the GluC-digested PbsA3-BCDE in 100% $D_2O$ containing 0.1% formic acid-$d_2$.**

Both beta protons of Phe8-OH and Phe10-OH showed a cross peak to their respective Cδ1 signals at 153.4 ppm and 152.6 ppm, and a cross peak to their respective Cδ2 carbon signals (129.7 ppm for Phe8-OH and 125.4 ppm for Phe10-OH). Sequence of the analyzed peptide is shown at the top. Modified residues are indicated in brackets.

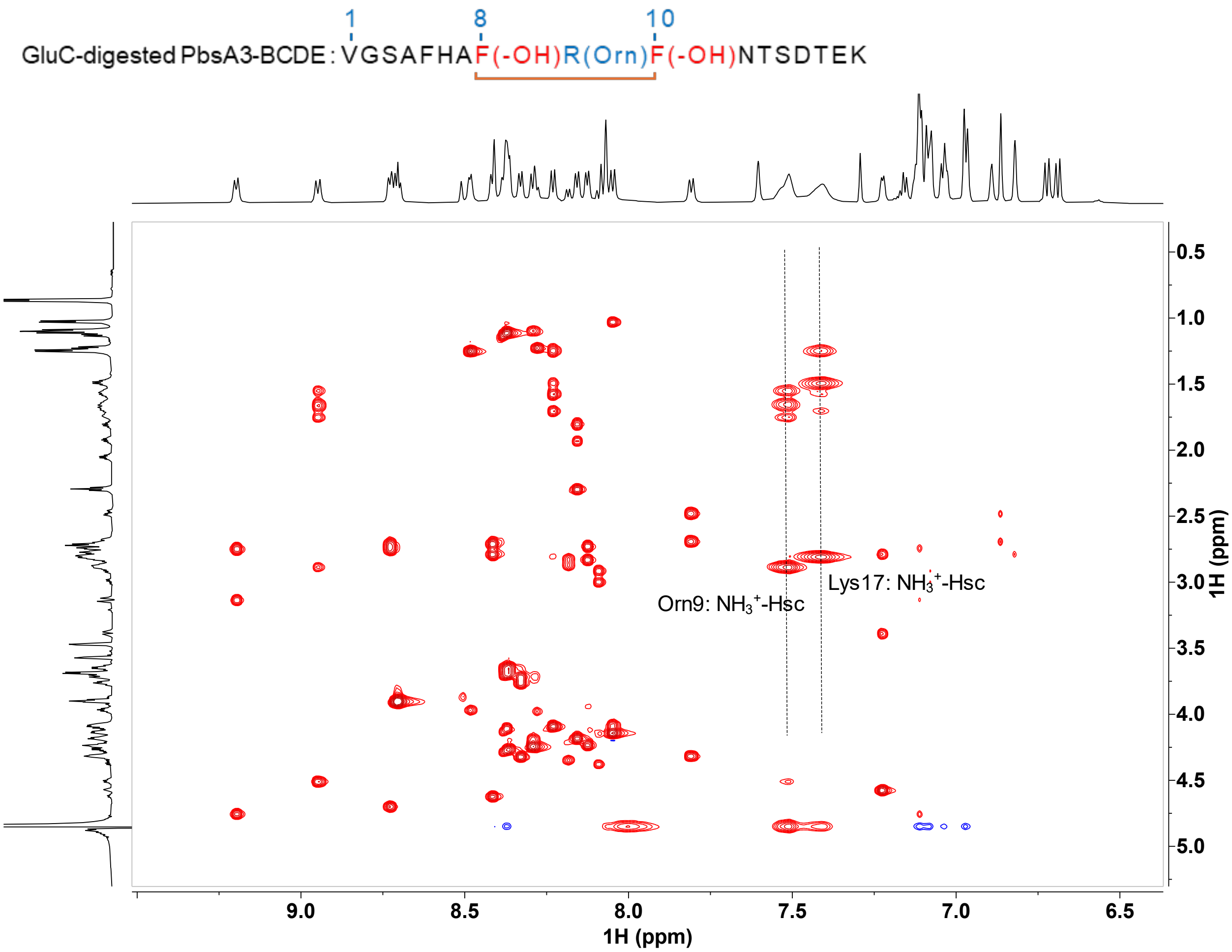


**Figure S14: 2D $^1H$-$^1H$ TOCSY spectra of the GluC-digested PbsA3-BCDE in 90% $H_2O$ and 10% $D_2O$ containing 0.1% formic acid-$d_2$ at 3 °C.**

A broad peak at 7.51 ppm was observed, which exhibited cross peaks with the other side chain protons (labeled as $H_{sc}$ in the figure) of residue 9. This peak integrated to 3 protons at low temperature (3 °C) similar to that of the PbsA3-CDE, corresponding to the loss of the urea group of Arg9, transforming it into an Orn residue. Sequence of the analyzed peptide is shown at the top. Modified residues are indicated in brackets.

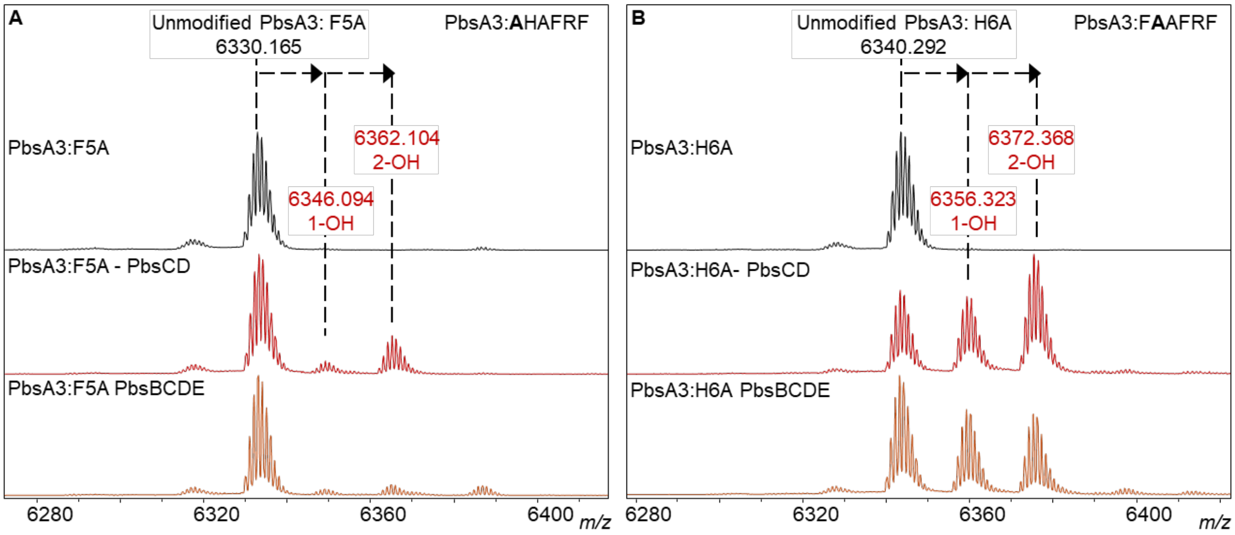


**Figure S15: Importance of conserved residues Phe5 and His6 for modification by *pbs* pathway enzymes.**

(A) Phe5 residue of the FHAFRF conserved motif in PbsA3 was mutated to Ala (F5A mutant). MALDI ToF MS spectra of PbsA3:F5A mutant alone (black spectra) or when co-expressed with PbsCD (red spectra) and PbsBCDE (orange spectra) is shown. Diminished PbsCD activity was observed further resulting in no downstream PbsE and PbsB-mediated modification. (B) His6 residue of the FHAFRF conserved motif in PbsA3 was mutated to Ala (H6A mutant). MALDI MS spectra of PbsA3:H6A mutant alone (black spectra) or when co-expressed with PbsCD (red spectra) and PbsBCDE (orange spectra) is shown. Wildtype-like modification was observed for PbsCD. However, no PbsE and PbsB activity was detected suggesting the significance of the H6 residue for PbsE activity.

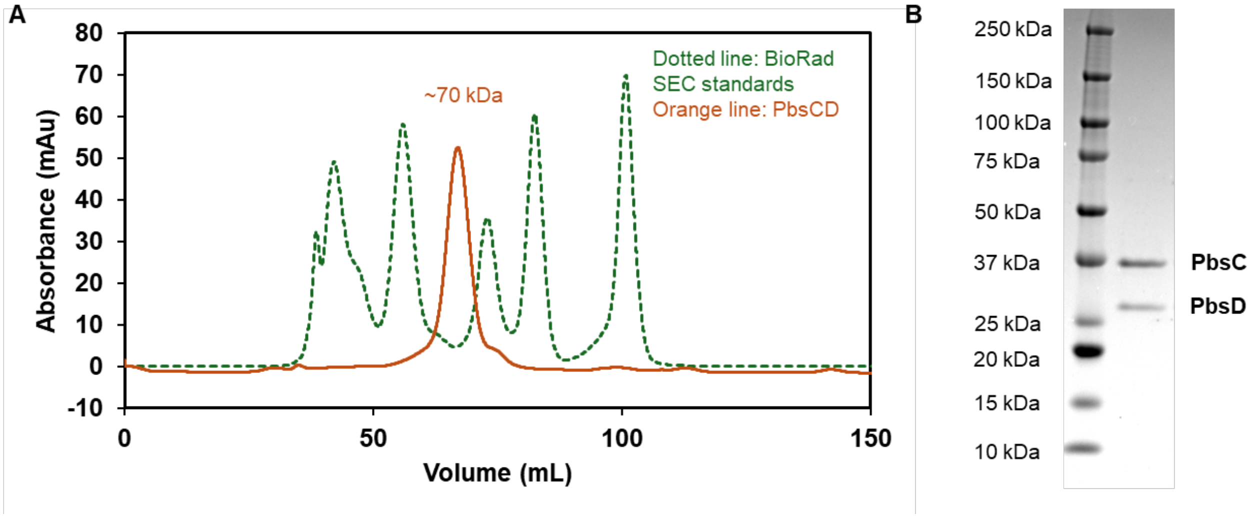


**Figure S16: Co-purification of PbsCD.**
PbsC and PbsD were co-purified by expressing N-terminally 6xHis-tagged PbsC and untagged PbsD. (A) Size exclusion chromatogram (SEC) of PbsCD is shown as a UV trace at 280 nm. PbsCD co-purified as a heterodimer of PbsC and PbsD monomers (orange trace). Green dotted lines represent SEC standards (Left to right: 670 kDa, 158 kDa, 44 kDa, 17 kDa, 1.35 kDa). (B) Denaturing SDS-PAGE image of PbsC and PbsD bands collected from one of the SEC fractions.

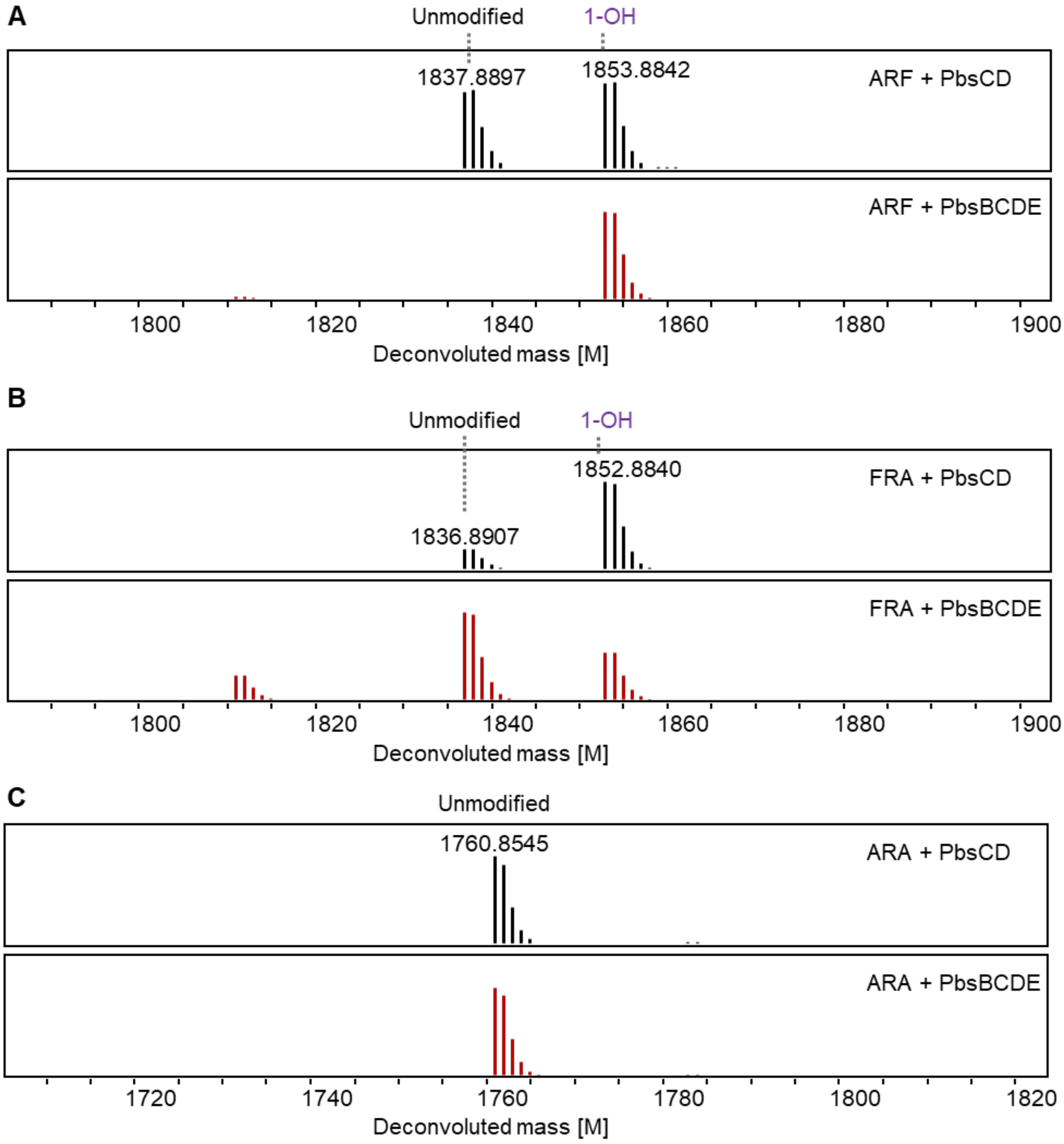


**Figure S17: Substrate scope assessment of *pbs* pathway enzymes against Ala mutants.**
ESI-HR-MS analysis of the extracted ions showing the isotopic peak distribution for the GluC-digested (A) PbsA3-F8A, **A**RF mutant, (B) PbsA3-F10A, FR**A** mutant and (C) the PbsA3-F8A:F10A, **A**R**A** mutant when co-expressed with PbsCD (black spectra and PbsBCDE (red spectra) in *E. coli*. Mono-hydroxylation catalyzed by PbsCD was observed in **A**RF and FR**A** mutants with no downstream PbsE and PbsB activity. No modifications were observed for the **A**R**A** mutant.

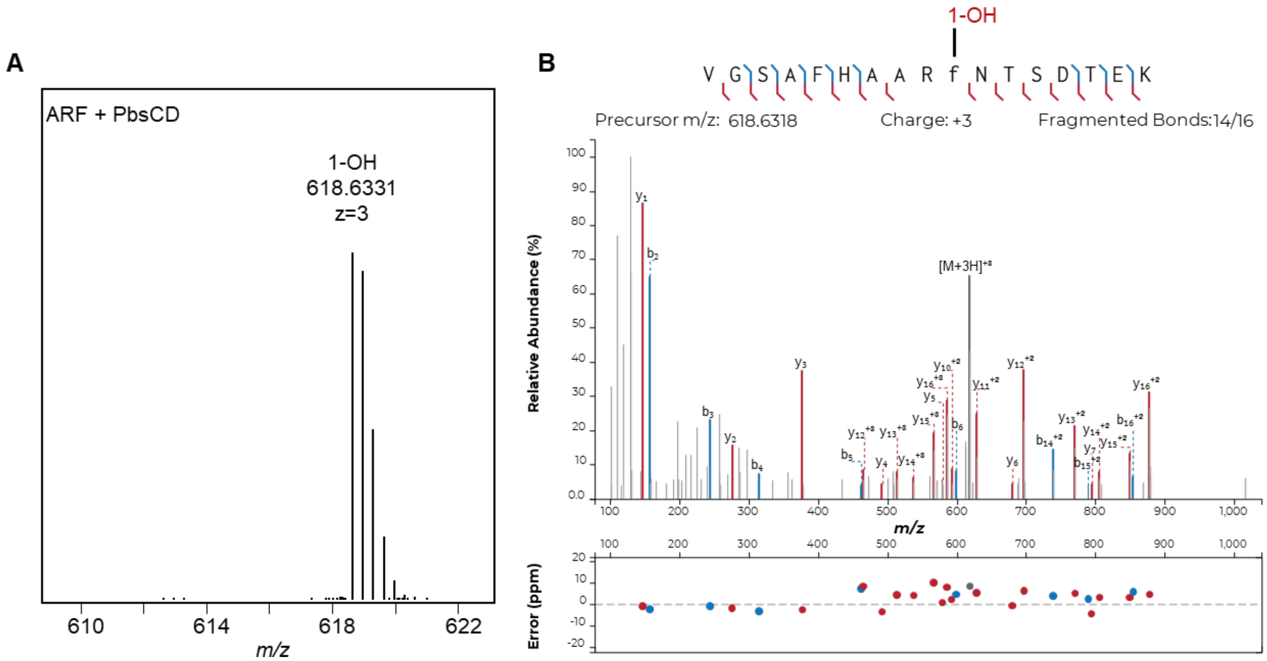


**Figure S18: ESI-HR-MS and MS/MS analysis of PbsCD-modified PbsA3-F8A.** (A) HR-MS spectra of the extracted ion showing the isotopic peak distribution for GluC-digested PbsA3-ARF co-expressed with PbsCD. Calcd. mass of mono-hydroxylated product $[M+3H]^{3+}$ = 618.6318 Da, obs. = 618.6331 Da (B) HR-MS/MS analysis of the mutant showing the b and y fragment ions. Hydroxylated residues are in lower case and annotated as 1-OH. Plots were prepared using the Interactive Peptide Annotator Webtool.[1]

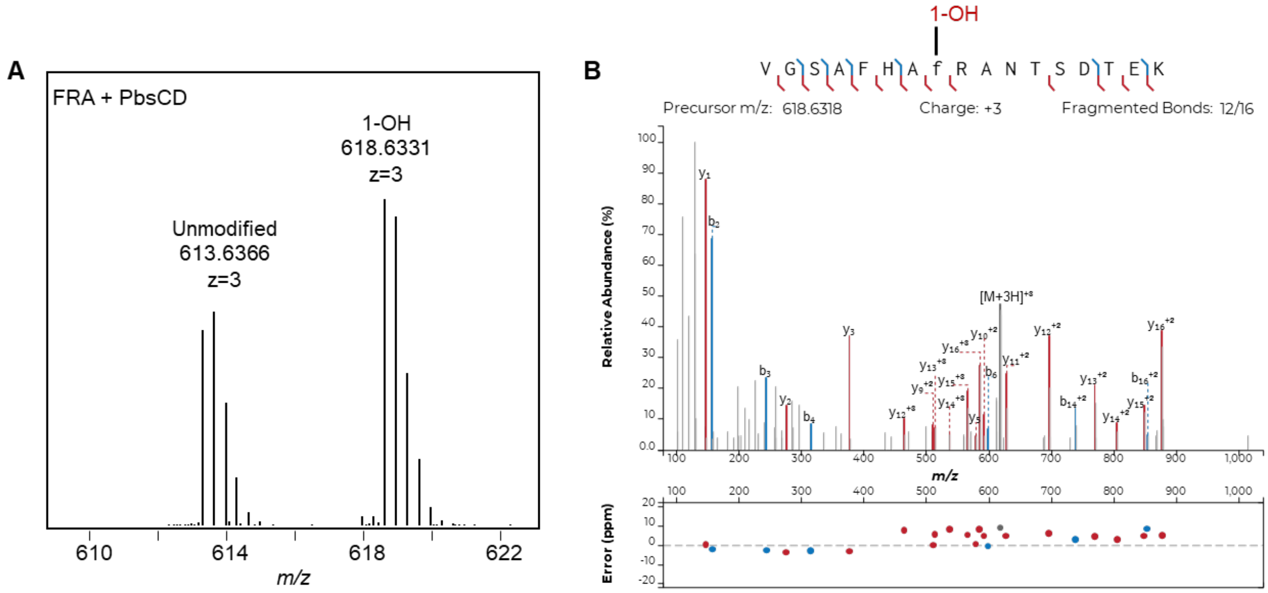


**Figure S19: ESI-HR-MS and MS/MS analysis of PbsCD-modified PbsA3-F10A.** (A) HR-MS spectra of the extracted ions showing the isotopic peak distribution for GluC-digested PbsA3-FRA co-expressed with PbsCD. Calcd. mass $[M+3H]^{3+}$ = 618.6318 Da, obs. 618.6331 Da (B) HR-MS/MS analysis showing the b and y fragment ions. Hydroxylated residues are in lower case and annotated as 1-OH. Plots were prepared using the Interactive Peptide Annotator Webtool.[1]

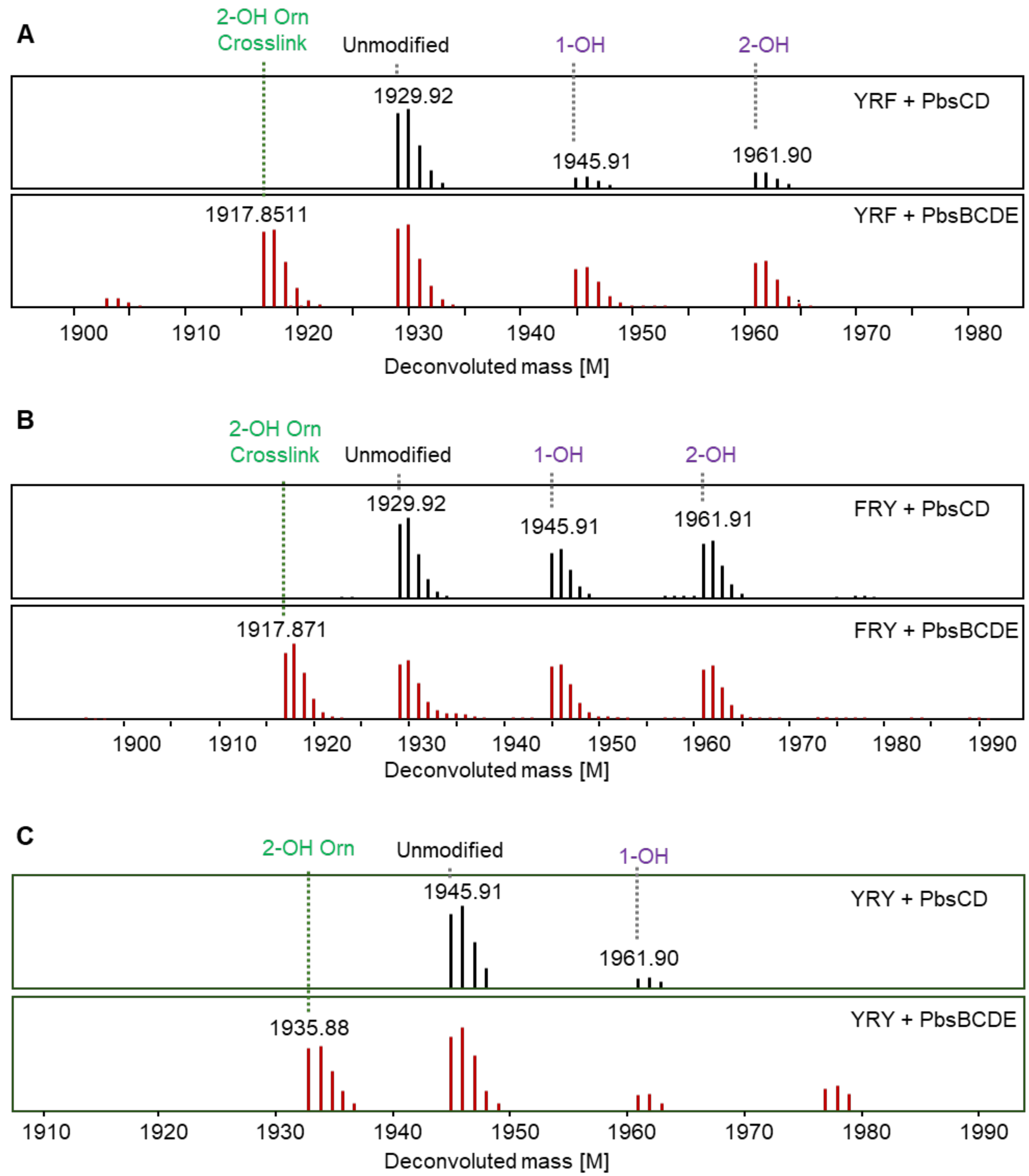


**Figure S20: Substrate scope assessment of *pbs* pathway enzymes with Phe-to-Tyr variants.** ESI-HR-MS analysis of the extracted ions showing the isotopic peak distribution for the GluC-digested (A) PbsA3-F8Y, YRF mutant (B) PbsA3-F10Y, FRY mutant and (C) the PbsA3-F8Y:F10Y, YRY mutant when co-expressed with PbsCD (black spectra), and PbsBCDE (red spectra) in *E. coli*.

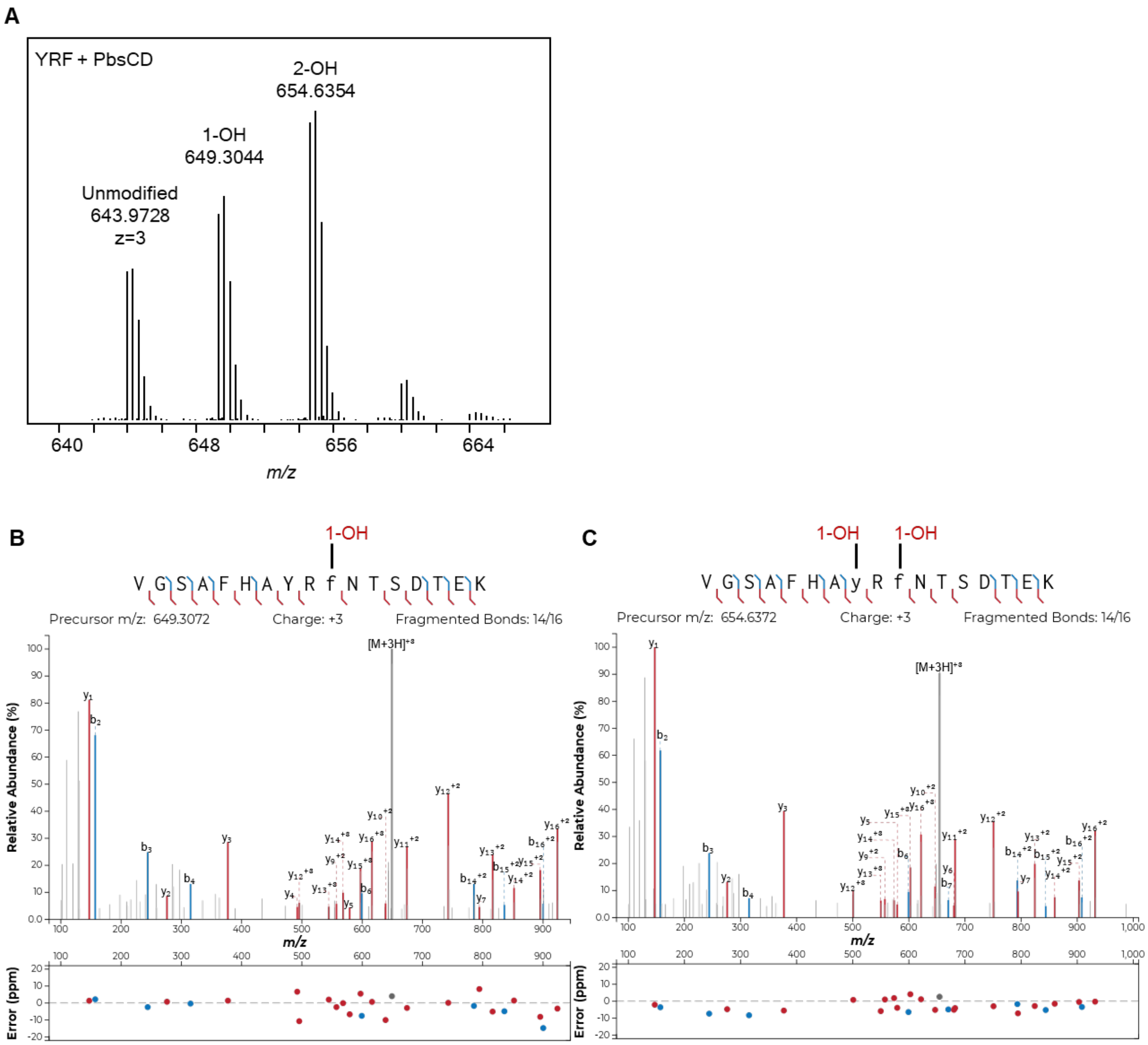


**Figure S21: ESI-HR-MS and MS/MS analysis of PbsCD-modified PbsA3-F8Y mutant.**

(A) HR-MS spectra of the extracted ions showing the isotopic peak distribution for the GluC-digested PbsA3-F8Y, **Y**RF mutant when co-expressed with PbsCD. Calcd. mass of mono-hydroxylated product $[M+3H]^{3+}$ = 649.3072 Da, observed mass $[M+3H]^{3+}$ = 649.3044 Da; bis-hydroxylated product $[M+3H]^{3+}$ = 654.6372 Da, observed mass $[M+3H]^{3+}$ = 654.6354 Da. HR-MS/MS analysis of the mutant showing the b and y fragment ions for the PbsCD-mediated (B) mono-hydroxylated product (C) bis-hydroxylated product. Hydroxylated residues are in lower case and annotated as 1-OH and 2-OH. Plots were prepared using the Interactive Peptide Annotator Webtool.[1]

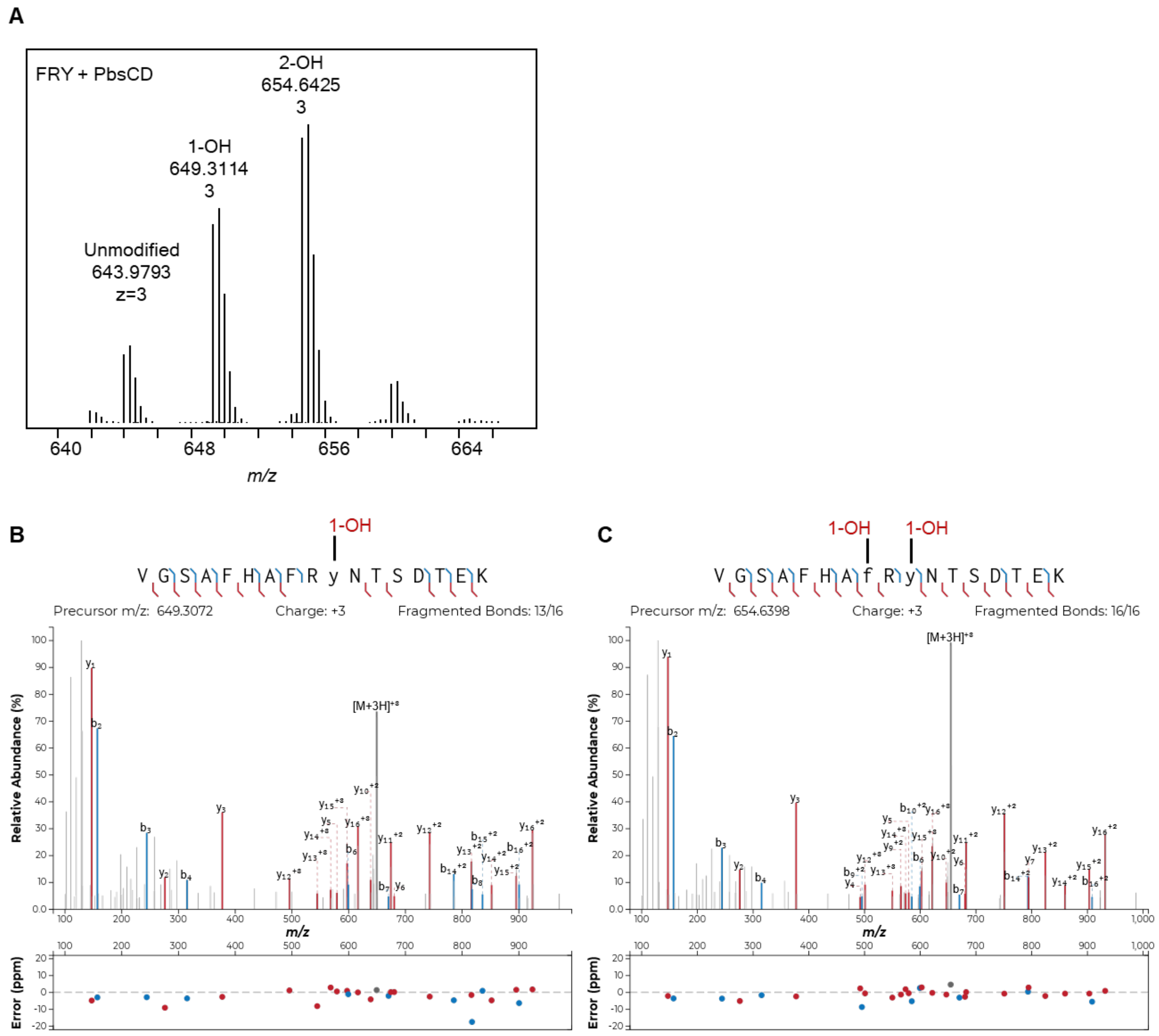


**Figure S22: ESI-HR-MS and MS/MS analysis of PbsCD-modified PbsA3-F10Y mutant.**
(A) HR-MS spectra of the extracted ions showing the isotopic peak distribution for the GluC-digested PbsA3-F10Y, FR**Y** mutant when co-expressed with PbsCD. Calcd. mass of mono-hydroxylated product $[M+3H]^{3+}$ = 649.3072 Da, observed mass $[M+3H]^{3+}$ = 649.3044 Da; bis-hydroxylated product $[M+3H]^{3+}$ = 654.6372 Da, observed mass $[M+3H]^{3+}$ = 654.6354 Da. HR-MS/MS analysis of the mutant showing the b and y fragment ions for the PbsCD-mediated (B) mono-hydroxylated product (C) bis-hydroxylated. Hydroxylated residues are in lower case and annotated as 1-OH and 2-OH. Plots were prepared using the Interactive Peptide Annotator Webtool.[1]

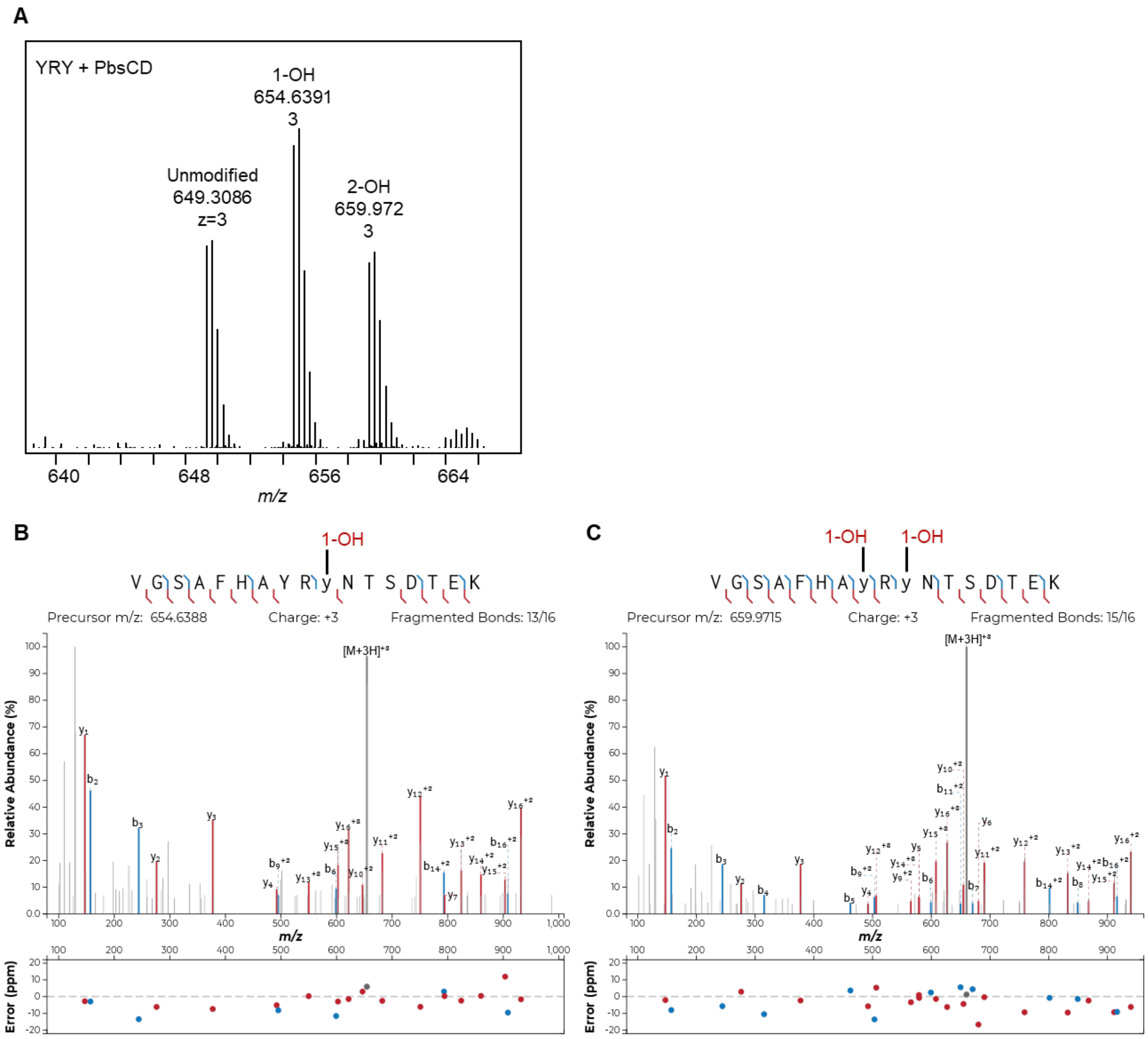


**Figure S23: ESI-HR-MS and MS/MS analysis of PbsCD-modified PbsA3-F8Y:F10Y mutant.**
(A) HR-MS spectra of the extracted ions showing the isotopic peak distribution for the GluC-digested PbsA3-F8Y:F10Y, **YRY** mutant when co-expressed with PbsCD. Calcd. mass of mono-hydroxylated product $[M+3H]^{3+}$ = 654.6388 Da, observed mass $[M+3H]^{3+}$ = 654.6391 Da; bis-hydroxylated product $[M+3H]^{3+}$ = 659.972 Da, observed mass $[M+3H]^{3+}$ = 659.9724 Da. HR-MS/MS analysis of the mutant showing the b and y fragment ions for the PbsCD-mediated (B) mono-hydroxylated product and (C) bis-hydroxylated product. Hydroxylated residues are in lower case and annotated as 1-OH and 2-OH. Plots were prepared using the Interactive Peptide Annotator Webtool.[1]

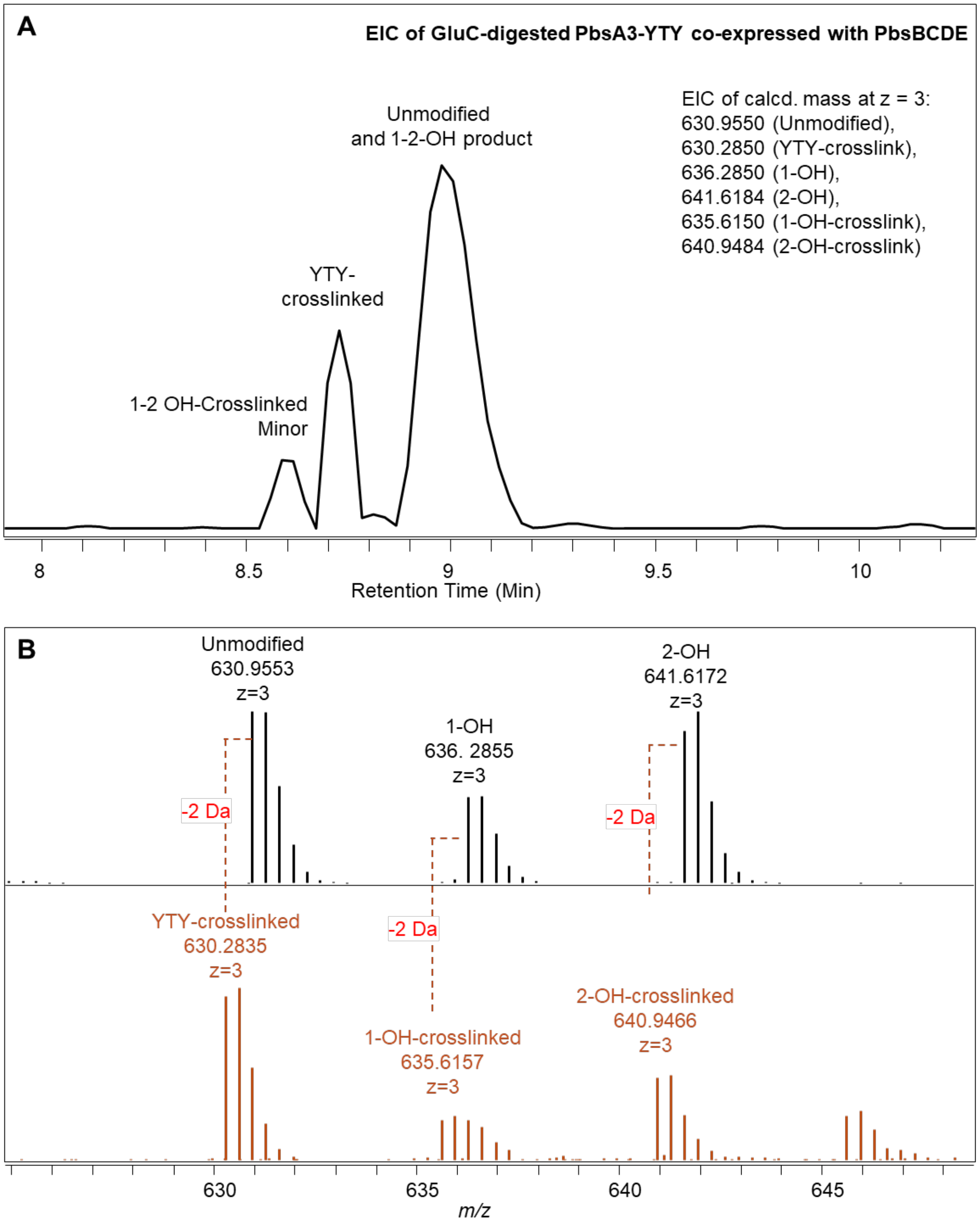


**Figure S24: LCMS and ESI-HR-MS analysis of PbsA3-YTY upon co-expression with PbsBCDE.** (A) Extracted ion chromatogram of PbsA3-YTY co-expressed with PbsBCDE is shown as GluC-digested products. (B) HR-MS spectra of the extracted ions showing the isotopic peak distribution for the unmodified GluC-digested PbsA3-YTY product, the mono-hydroxylated YTY-CD 1-OH peptide, and the bis-hydroxylated YTY-CD 2-OH product is shown in black spectra. The non-hydroxylated YTY-crosslinked product (YTY-B), the mono-hydroxylated YTY-BCD 1-OH crosslinked peptide (minor product), and the bis-hydroxylated YTY-BCD 2-OH crosslinked product (minor product) is shown in orange spectra.

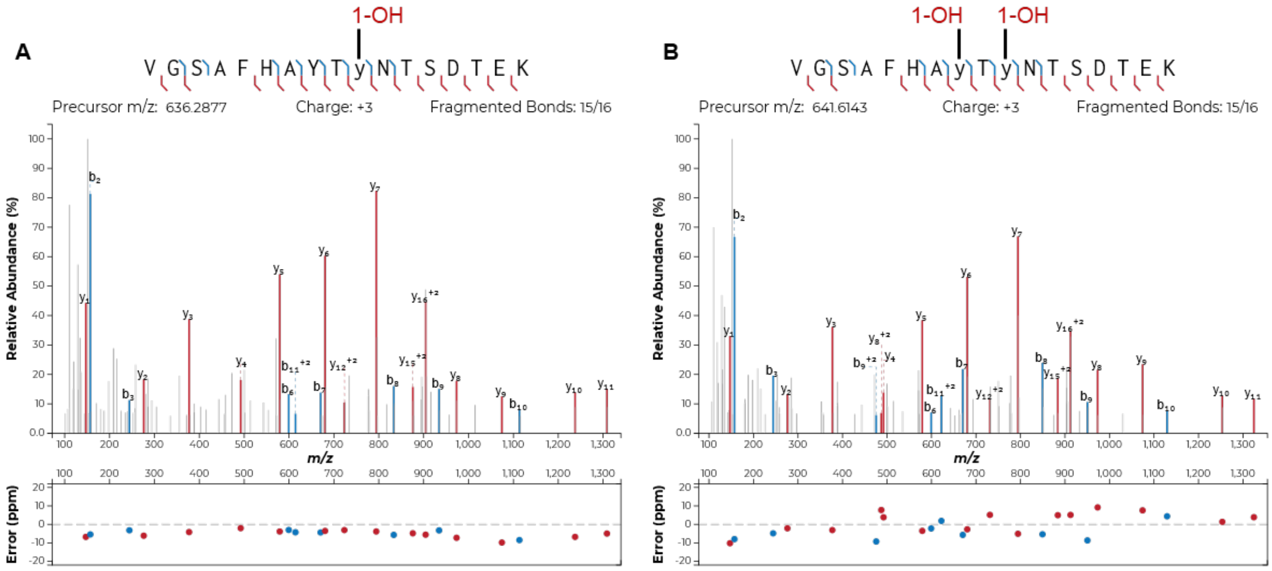


**Figure S25: ESI-HR-MS/MS analysis of PbsA3-YTY upon co-expression with PbsCD.**
HR-MS/MS spectra of (A) the GluC-digested mono-hydroxylated PbsA3-YTY-CD 1-OH product where the hydroxylation was localized on the Tyr10 residue and (B) the GluC-digested PbsA3-YTY-CD 2-OH product showing bis-hydroxylation on the Try8 and Tyr10 residues. Plots were prepared using the Interactive Peptide Annotator Webtool.[1]

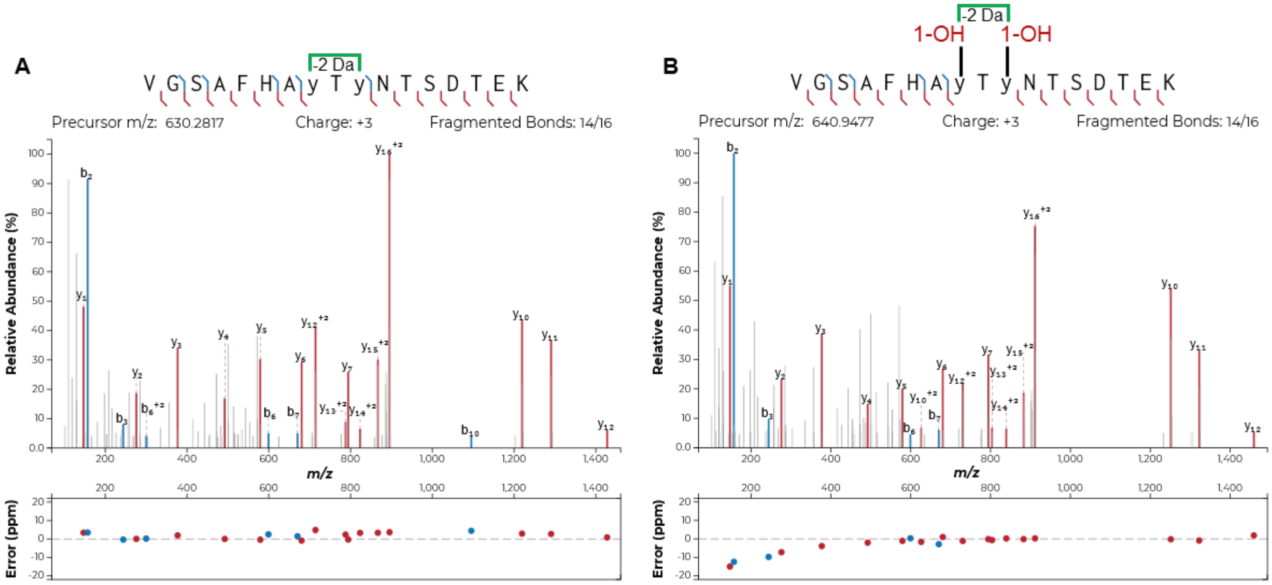


**Figure S26: ESI-HR-MS/MS analysis of PbsA3-YTY upon co-expression with PbsBCDE.**
HR-MS/MS spectra of (A) the GluC-digested PbsA3-YTY-B product showing crosslinking between the Try8 and Tyr10 residues and (B) the GluC-digested PbsA3-YTY-BCD product showing bis-hydroxylation and crosslinking on the Try8 and Tyr10 residues. Plots were prepared using the Interactive Peptide Annotator Webtool.[1]

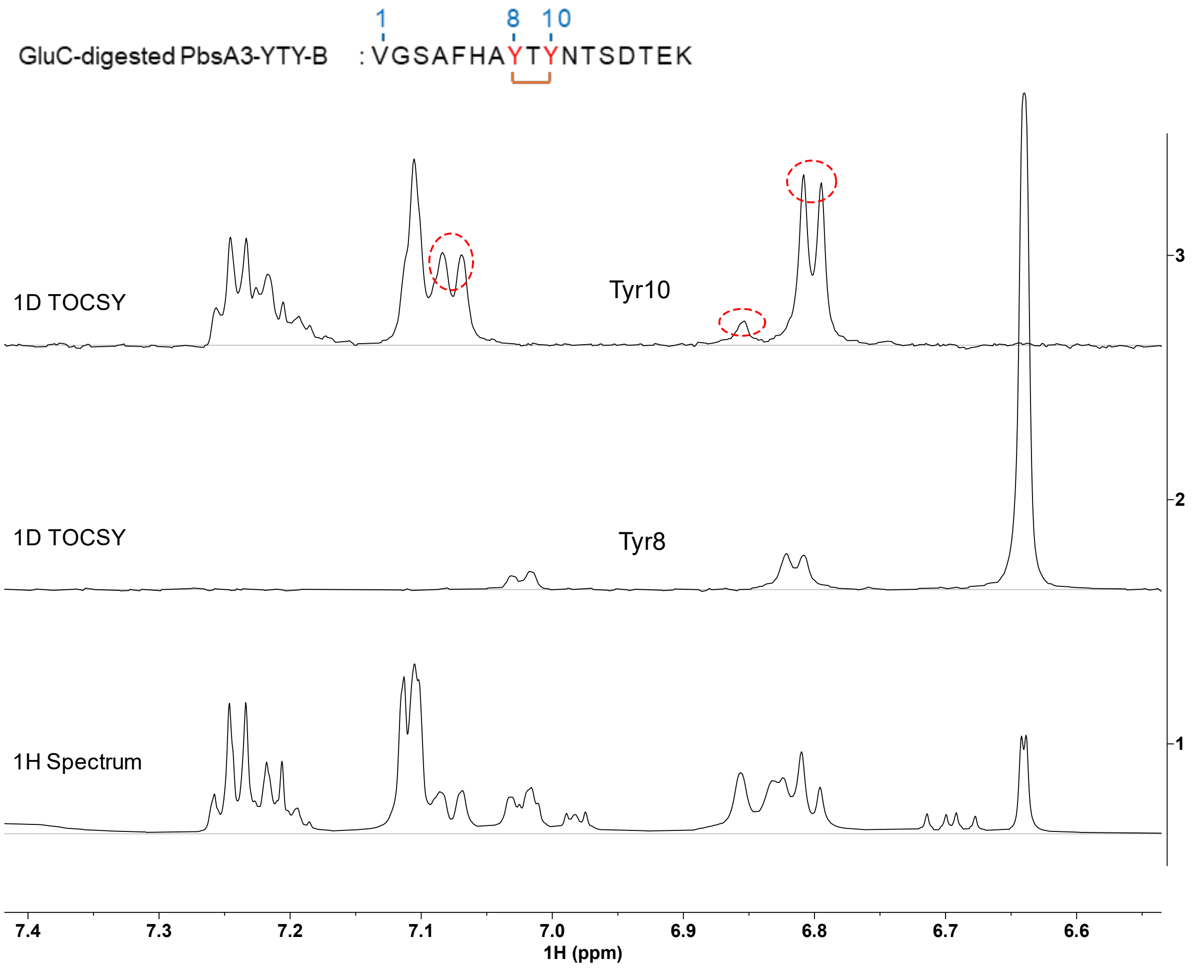


**Figure S27: 1D $^1$H-$^1$H TOCSY spectra of GluC-digested YTY-B in 90% $H_2O$ and 10% $D_2O$ and 0.1% FA-$d_2$ at 25 °C.** Three protons for each Tyr amino acid (Tyr10 and Tyr8) are observed (shown in red ovals), with two doublets and one singlet peak. The protons in Phe5 are shown as well in this region. Sequence of the analyzed peptide is shown at the top. Crosslinked residues are connected in orange brackets.

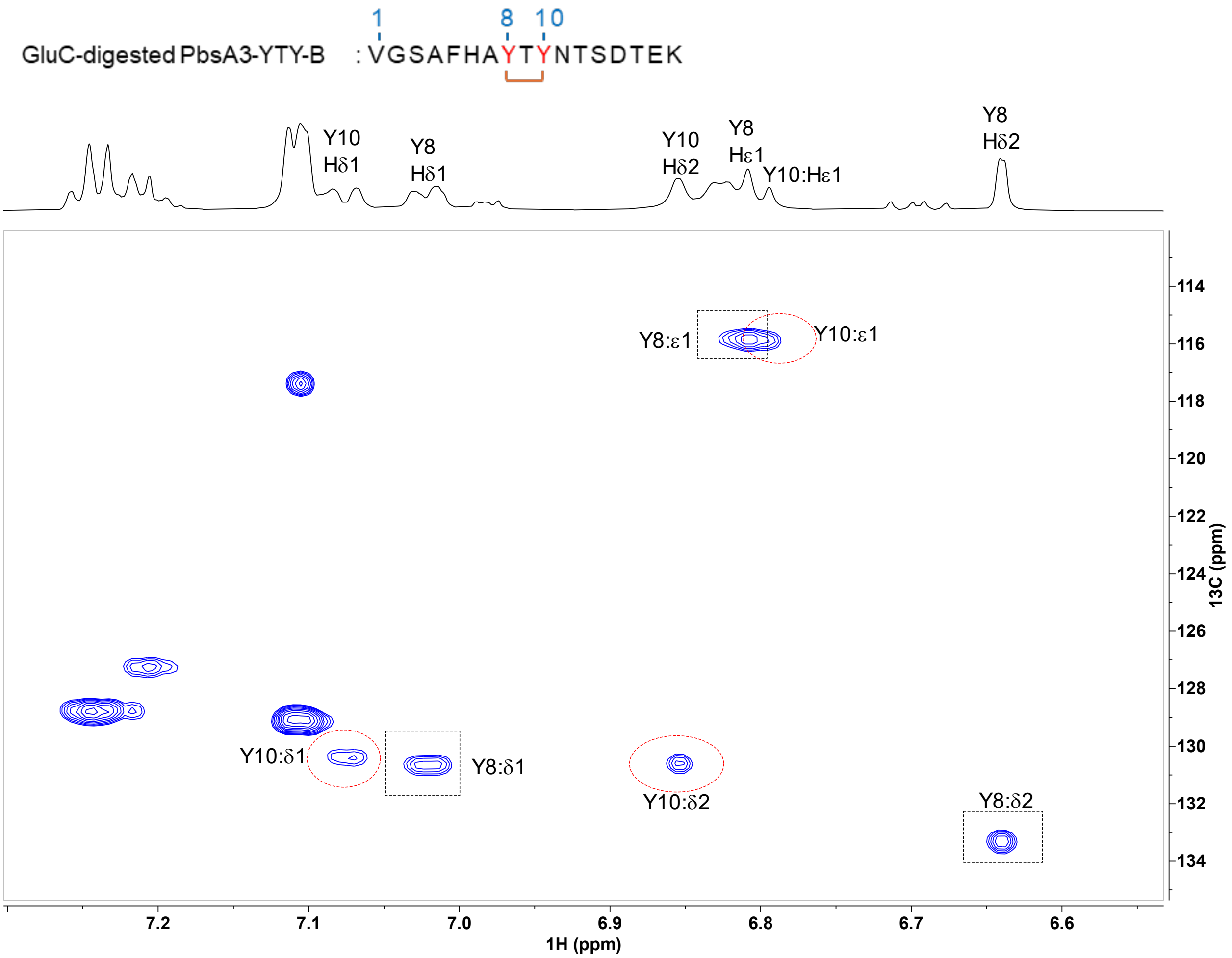


**Figure S28: $^1$H-$^{13}$C HSQC spectrum of the aromatic region for the GluC-digested YTY-B in 90% $H_2O$ and 10% $D_2O$ and 0.1% FA-$d_2$ at 25 ºC**. The expanded aromatic region show the Tyr8 and Tyr10 signals. The one-bond $^1$H-$^{13}$C cross peaks are highlighted in red ovals for Tyr10 and in black rectangles for Tyr8, respectively. The peaks with $^{13}$C chemical shifts around 116 ppm suggest crosslinking at the Cε1 of the Tyr residues. The sequence of the analyzed peptide is shown at the top. Crosslinked residues are connected in orange brackets.

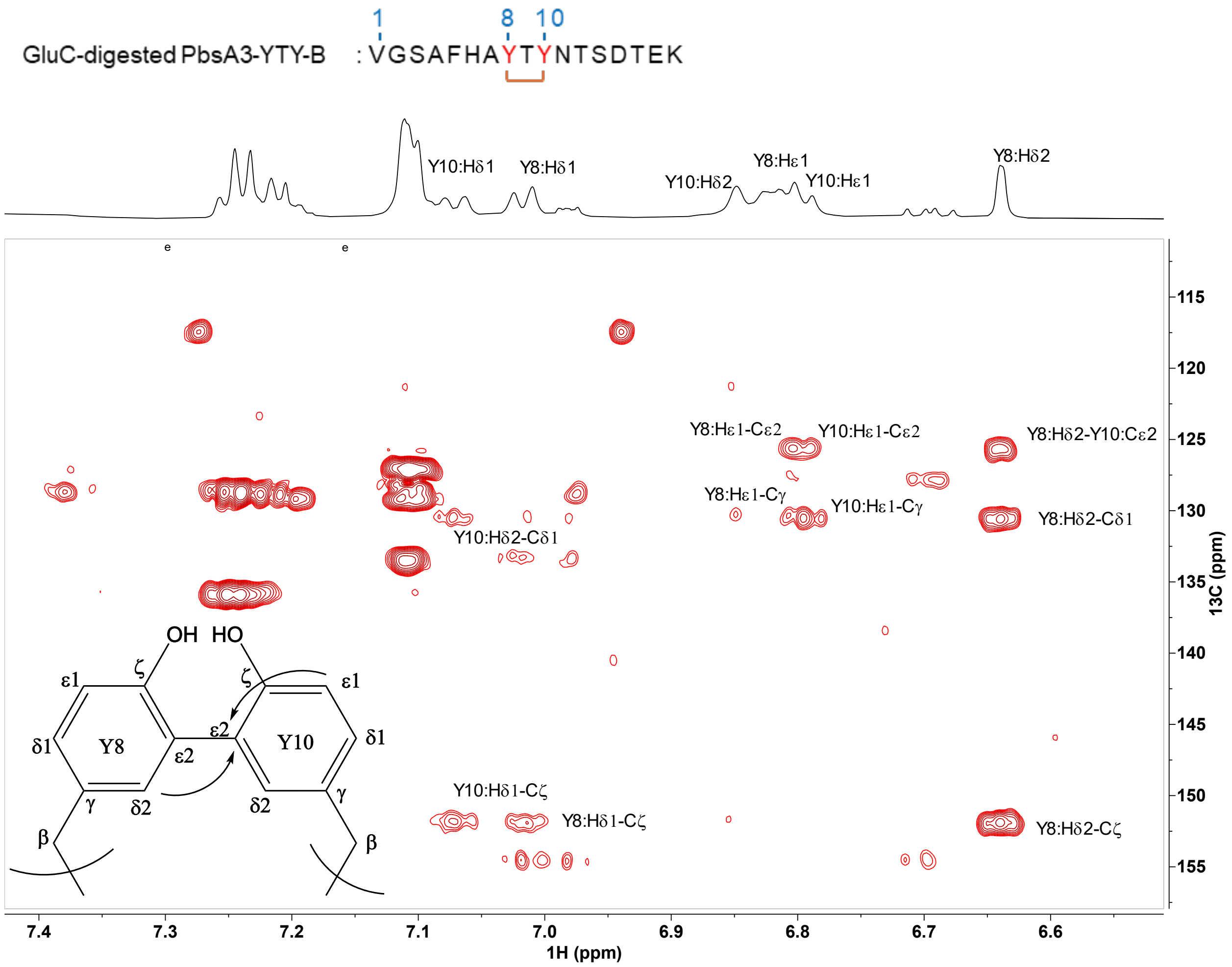


**Figure S29: $^{1}H$-$^{13}C$ HMBC of the aromatic region for the GluC-digested YTY-B in 90%$H_2O$ and 10% $D_2O$ and 0.1% FA-$d_2$ at 25 ºC.** The arrows show the C-C cross link between the two Tyr rings which was confirmed by the cross peak observed between Hδ2 of Tyr8 and Cε2 of Tyr10. The Hδ2 proton of Tyr8 at 6.64 ppm showed a cross peak to the Cε2 carbon of Tyr10 at 125.4 ppm. Some of the cross peaks in assistance to the assignment are labeled. Sequence of the analyzed peptide is shown at the top. Crosslinked residues are connected in orange brackets**.**

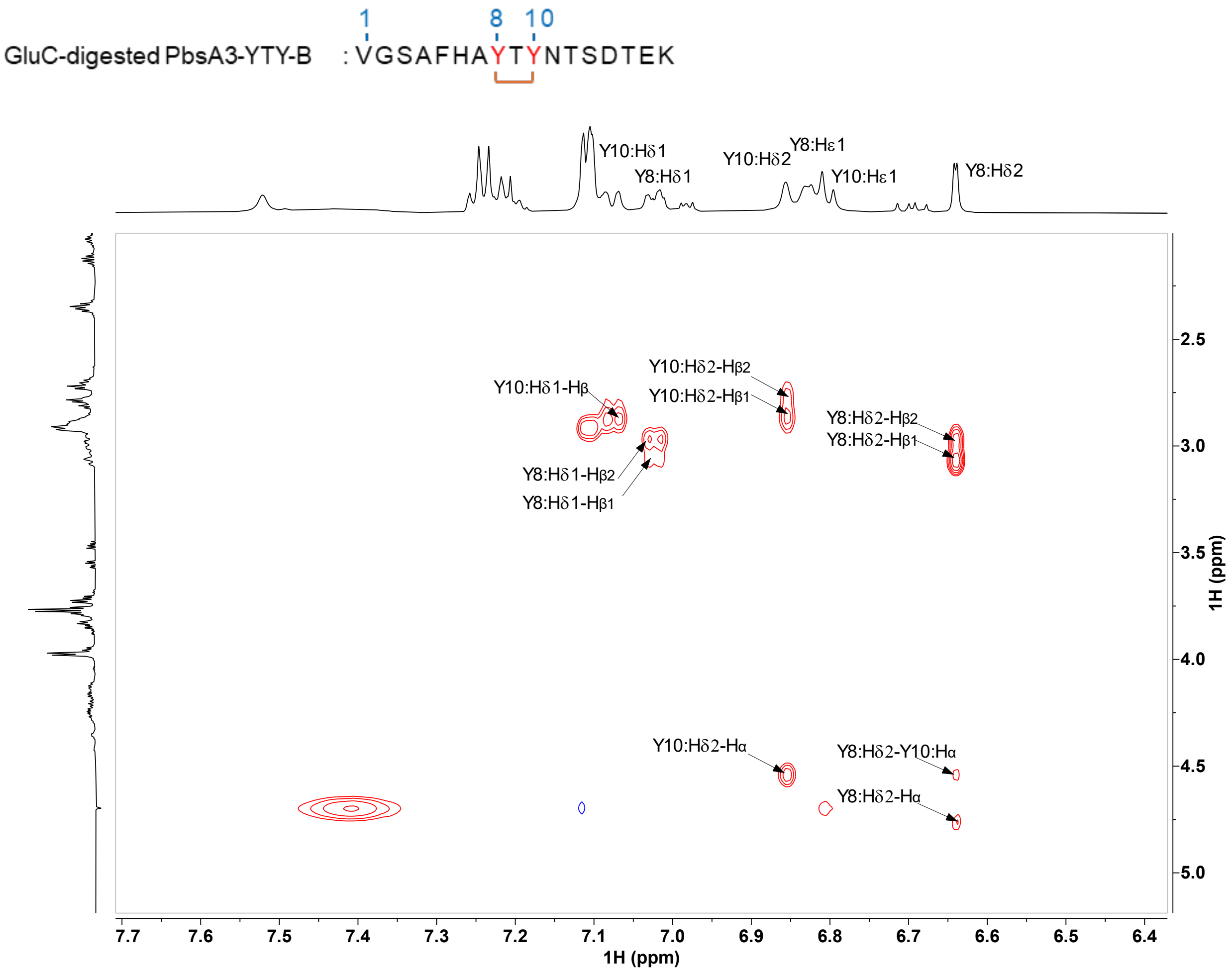


**Figure S30: $^1$H-$^1$H NOESY spectrum of the GluC-digested YTY-B in 90% $H_2O$ and 10% $D_2O$ and 0.1% FA-$d_2$ at 25 °C.** The NOE cross peaks between the aromatic protons Hδ1 and Hδ2 of Tyr8 and Tyr10 to their respective beta-protons are shown, further supporting that the cross-link occurring at the Cε2 position. Sequence of the analyzed peptide is shown at the top. Crosslinked residues are connected in orange brackets.

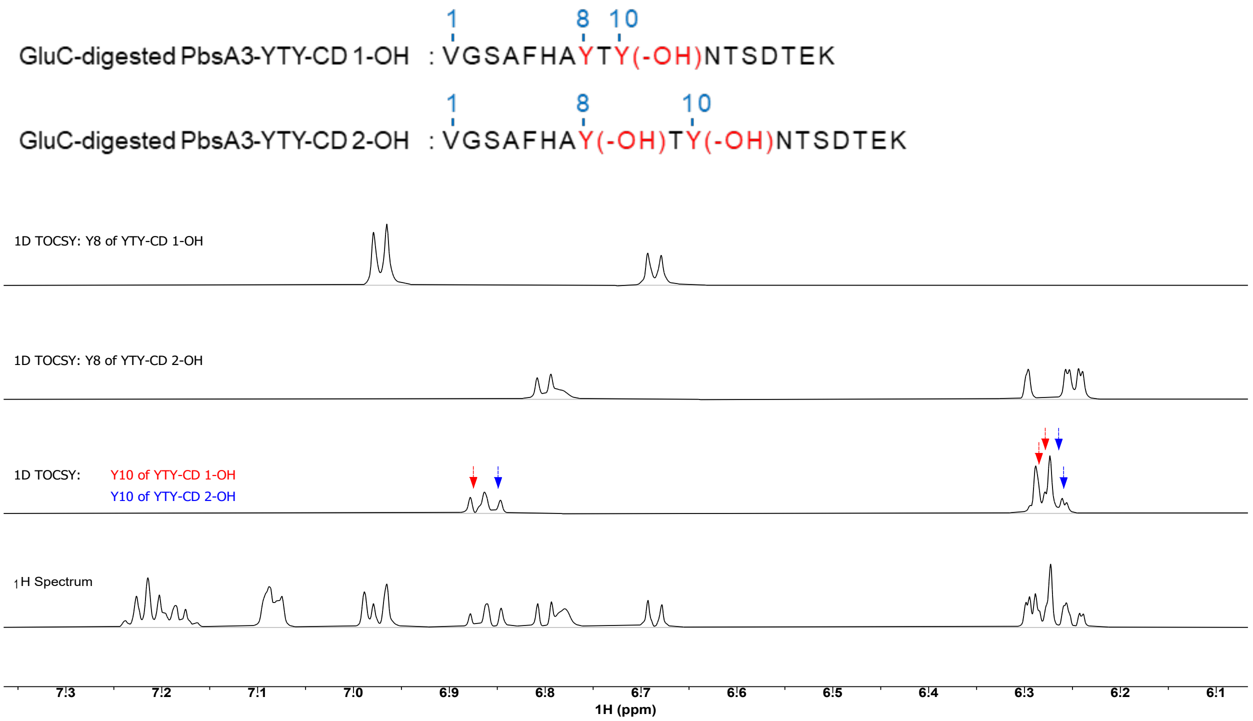


**Figure S31: 1D $^1$H-$^1$H TOCSY spectra of the GluC-digested YTY-CD 1-OH and YTY-CD 2-OH peptides in 90% $H_2O$ and 10% $D_2O$ at 25 ºC.** Two distinct types of spin system were observed. The original Tyr pattern of Try8 in the YTY-CD 1OH peptide (top spectrum), and three protons for each Tyr amino acid (Tyr10 of YTY-CD 1-OH; Tyr8 and Tyr10 of YTY-CD 2-OH) were observed, with two doublets peak and one singlet peak. The red and blue arrows highlight the annotated Tyr10 residues of YTY-CD 1OH and YTY-CD 2OH, respectively. Sequence of the analyzed peptides are shown at the top. Modified residues are indicated in brackets.

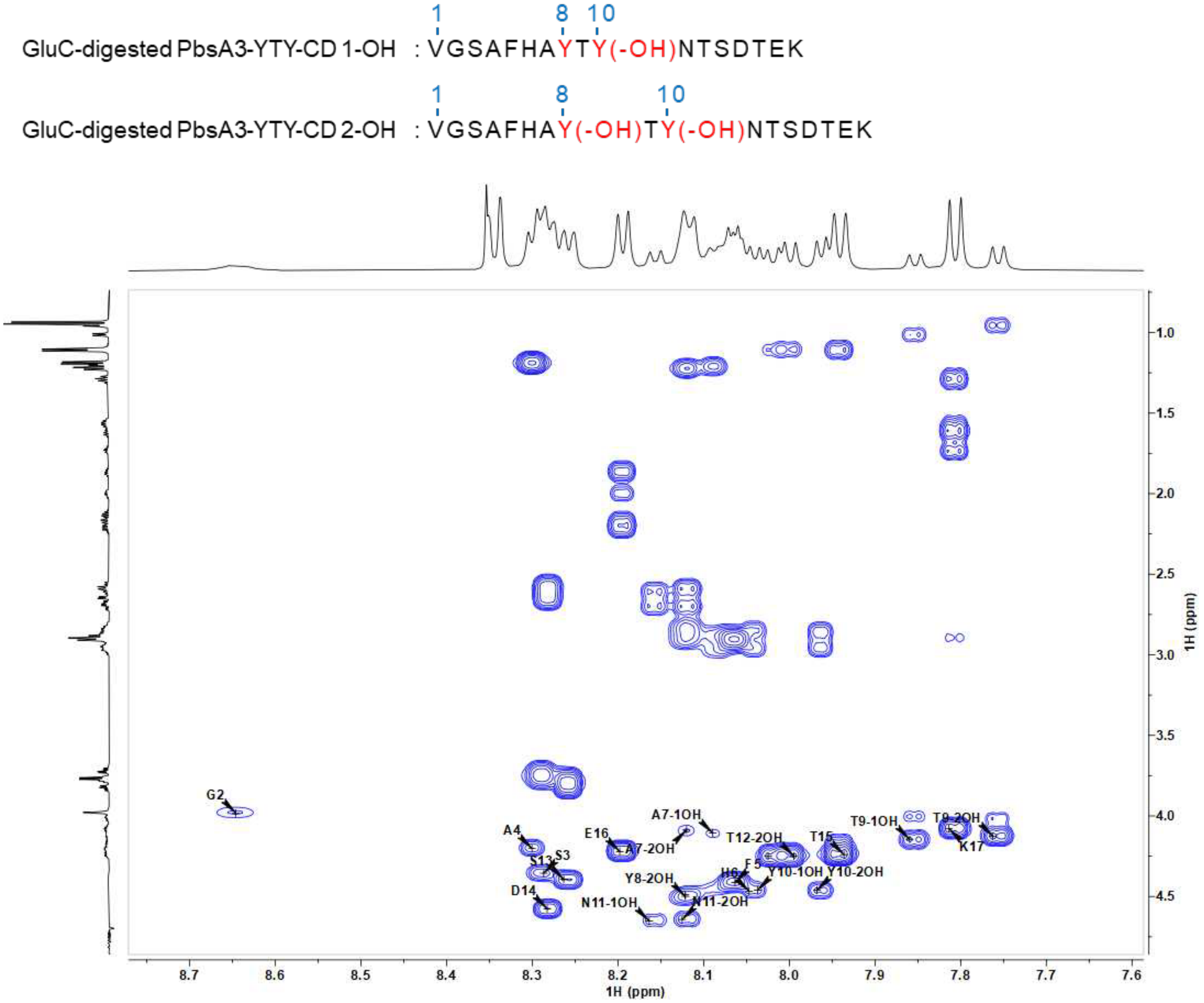


**Figure S32: 2D $^1$H-$^1$H TOCSY spectra of the GluC-digested YTY-CD 1-OH and YTY-CD 2-OH peptides in 90% $H_2O$ and 10% $D_2O$ at 25 ºC.** The cross peaks between HN and Hα of each residue in the two peptides are labeled. YTY-CD 1-OH and YTY-CD 1-OH are abbreviated as -1OH and -2OH in the figure for clarity. The peaks without -1OH or -2OH correspond to residues with the same chemical shifts in both peptides. Sequence of the analyzed peptides are shown at the top. Modified residues are indicated in brackets.

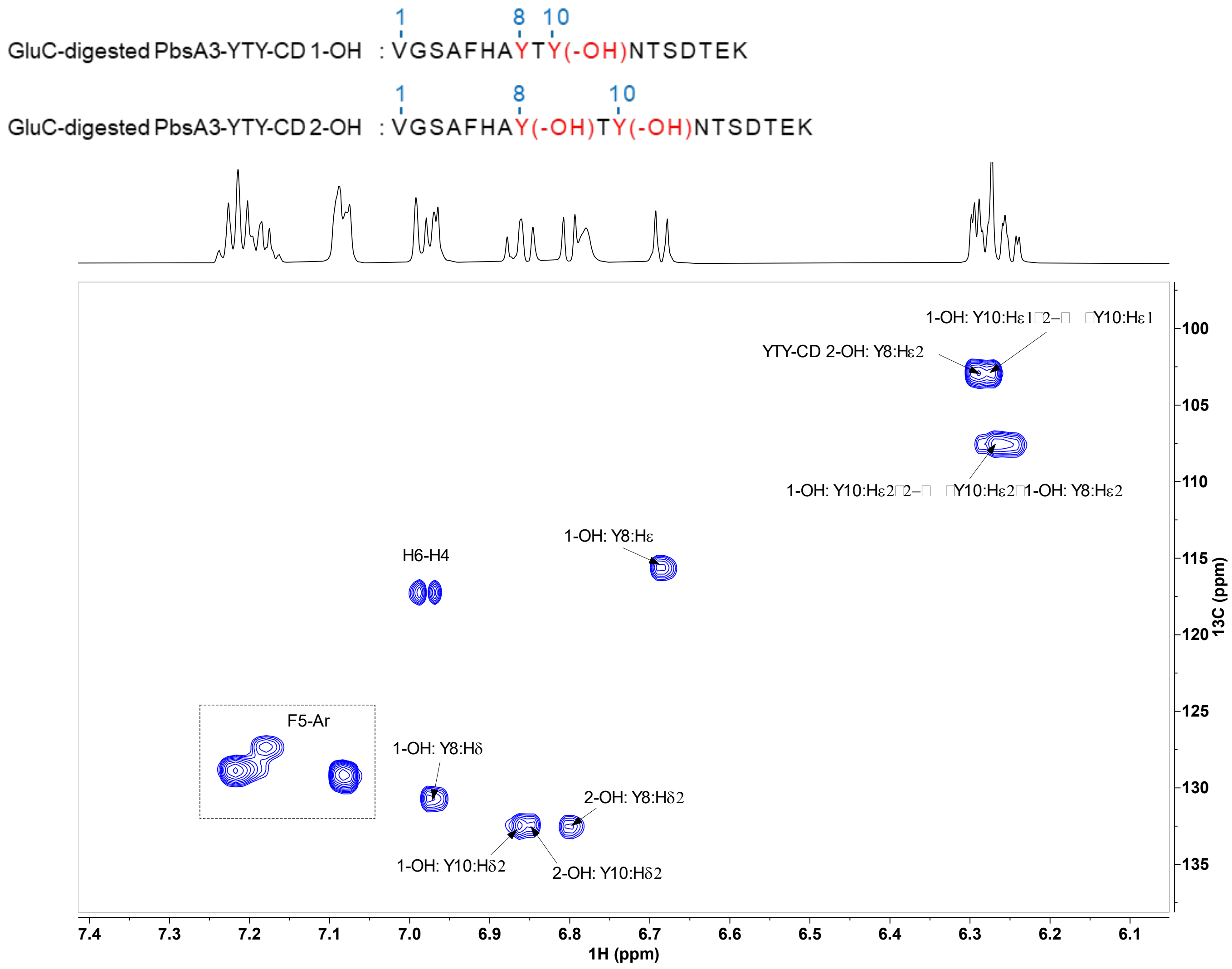


**Figure S33: Expanded aromatic region of $^{1}H$-$^{13}C$ HSQC spectra of the GluC-digested YTY-CD 1-OH and YTY-CD 2-OH peptides in 90%$H_2O$ and 10% $D_2O$ at 25 ºC.** $^{1}H$-$^{13}C$ HSQC spectrum of YTY-CD 1-OH and YTY-CD 2-OH showing the cross peaks in the zoomed-in aromatic region of Tyr8 and Tyr10 residues in the two peptides, isolated as a 2:3 mixture, respectively. The one-bond $^{1}H$-$^{13}C$ cross peaks are labeled. For clarity, the -CD was eliminated in the labels. Sequence of the analyzed peptides are shown at the top. Modified residues are indicated in brackets.

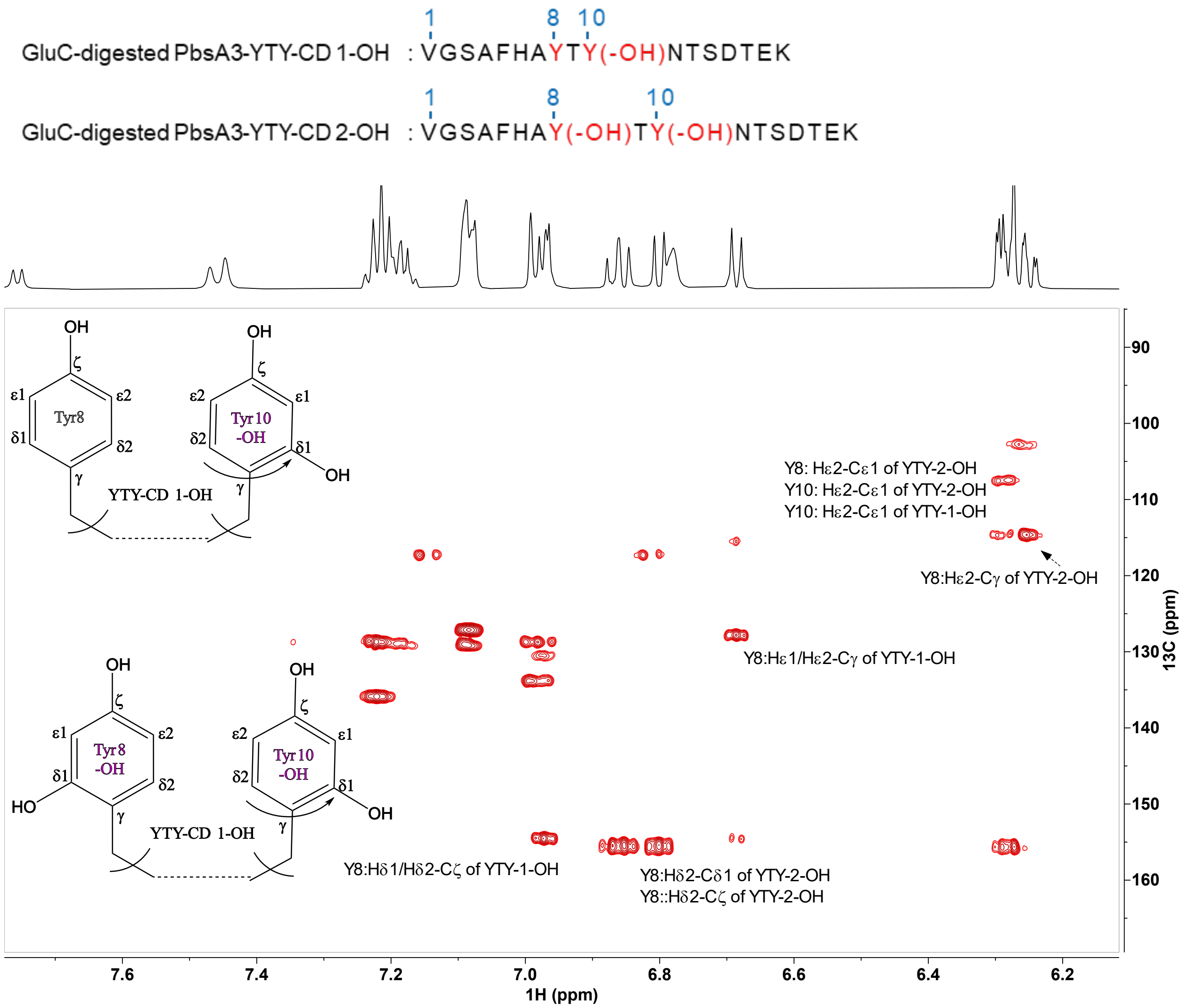


**Figure S34: The aromatic region $^{1}$H-$^{13}$C HMBC spectra of the GluC-digested YTY-CD 1-OH and YTY-CD 2-OH peptides in 90% $H_2O$ and 10% $D_2O$ at 25 °C.** The cross peaks between the Hε1 to Cε2 or vice versa of each Tyr ring were observed. Also, the cross peak between Hδ2 to Cδ1 was observed for Tyr10 in YTY-CD 1-OH, Tyr8 and Tyr10 in YTY-CD 2-OH, respectively, as a downfield shifted carbon (black arrow), indicating the hydroxylation occurring at this carbon. Relevant cross peaks are highlighted. The -CD in the naming is eliminated for clarity. Sequence of the analyzed peptides are shown at the top. Modified residues are indicated in brackets.

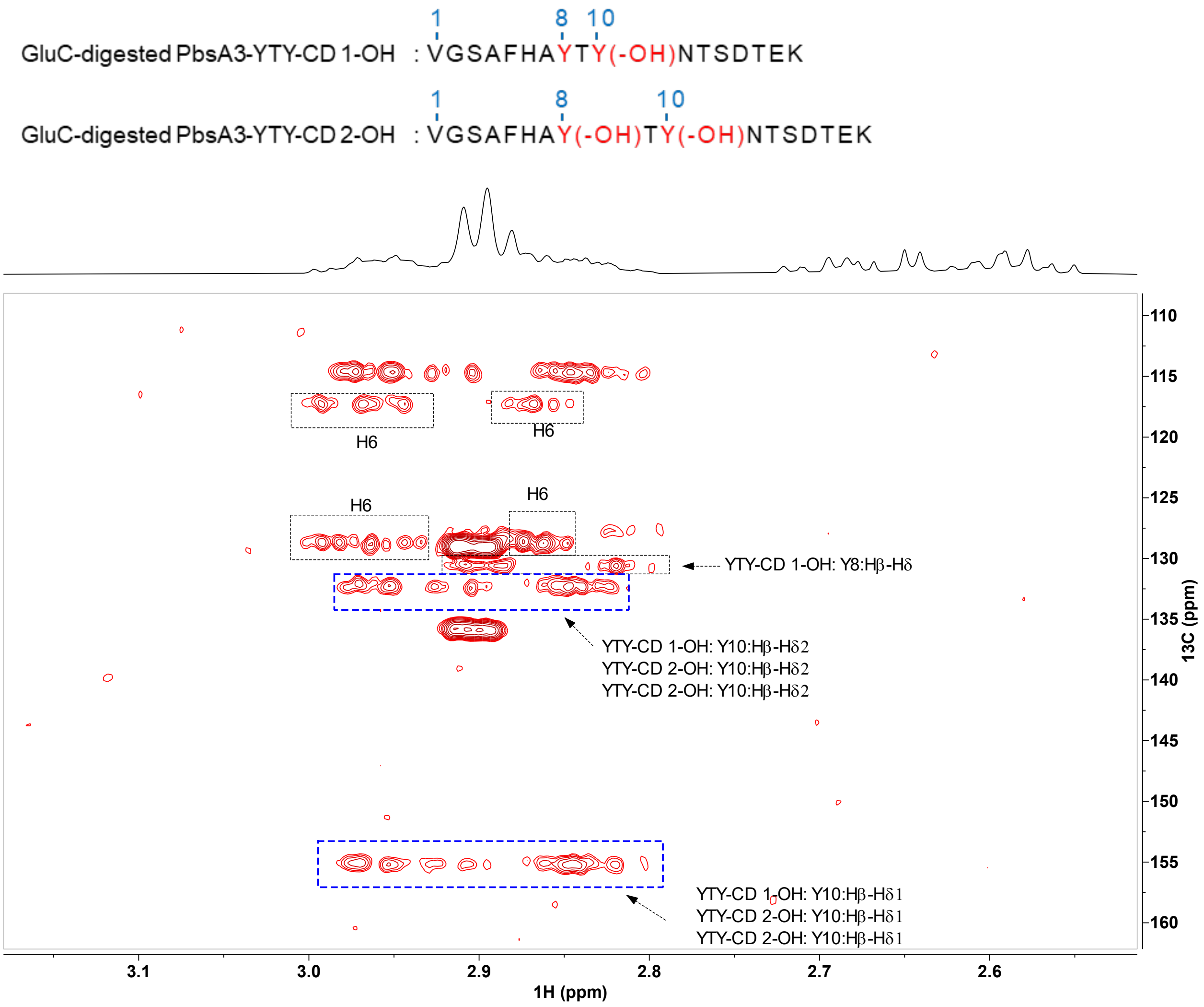


**Figure S35: The aliphatic region $^{1}H$-$^{13}C$ HMBC spectra of the GluC-digested YTY-CD 1-OH and YTY-CD 2-OH peptides in 90% $H_2O$ and 10% $D_2O$ at 25 °C.** The beta protons of Tyr 10 in YTY-CD 1-OH and beta protons of Tyr 8 and Tyr 8 in YTY-CD 2-OH show cross peaks to the Cδ1 carbons at 155.3 ppm, confirming the site of hydroxylation. The relevant cross peaks are highlighted. Sequence of the analyzed peptide is shown at the top. Modified residues are indicated in brackets.

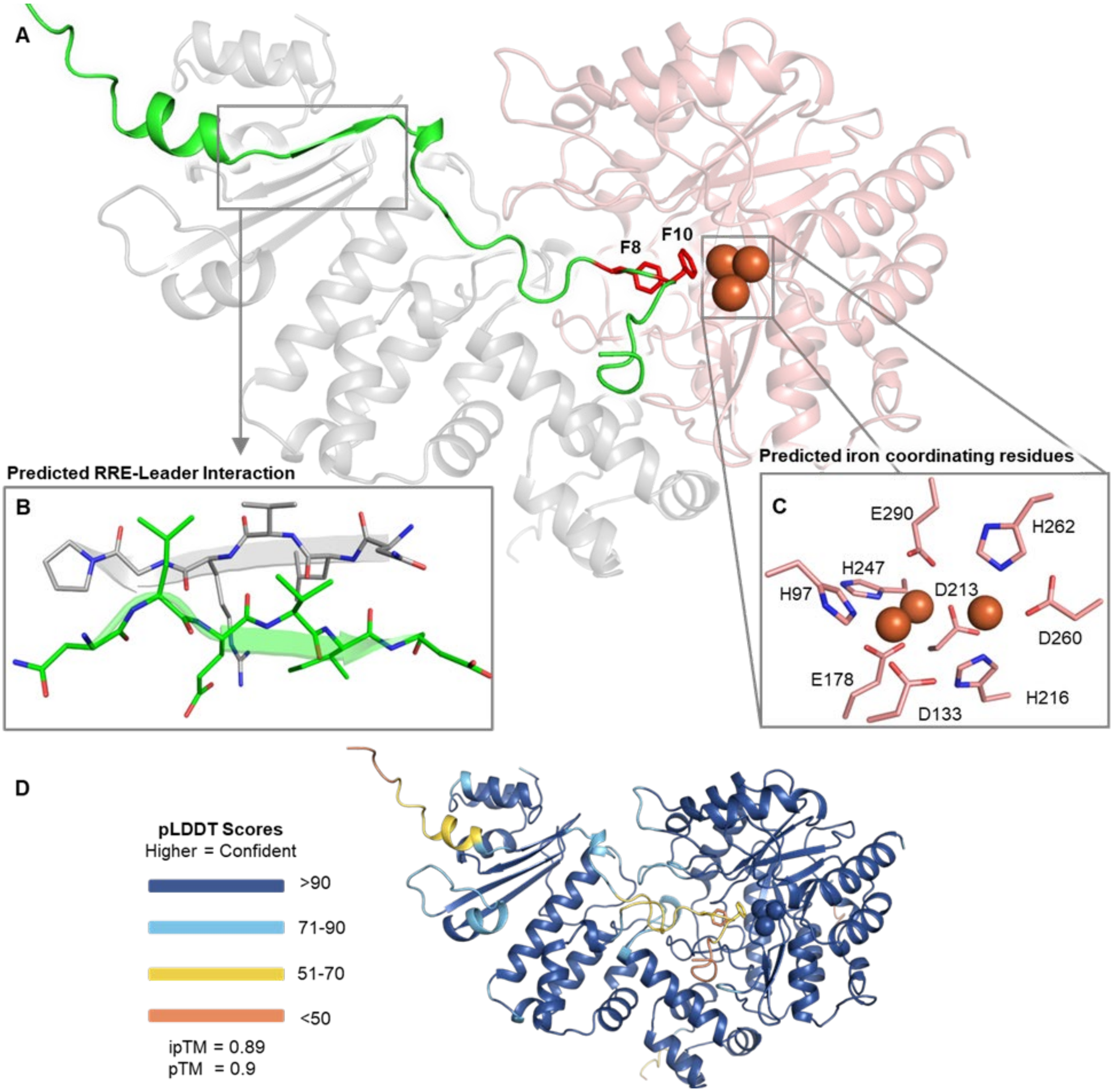


**Figure S36: AlphaFold 3 model of PbsCD in complex with PbsA2.** (A) Alphafold 3-predicted structure of PbsC and PbsD in complex with PbsA2. Three Fe ions were included in the model. PbsC and PbsD cartoon diagrams are shown in salmon and grey colors at a transparency setting of 60%. PbsA2 is displayed as a green cartoon diagram. The side chains of the Phe8 and Phe10 residues are in red sticks. Fe-ions are shown as orange spheres. (B) Zoomed-in image of the putative anti-parallel beta-sheet interaction between PbsD and the leader peptide of PbsA2. (C) Side chains of residues predicted to be ligands of the Fe ions. (D) Confidence metrics of the AlphaFold 3-predicted structures. pLDDT scores are shown as colored outputs (higher value means higher confidence as shown in the legend). The predicted template modeling (pTM) score and the interface predicted template modeling (ipTM) scores measure the accuracy of the entire structure (pTM>0.5 means higher similarity of predicted and true structures; iPTM>0.8 means confident prediction).

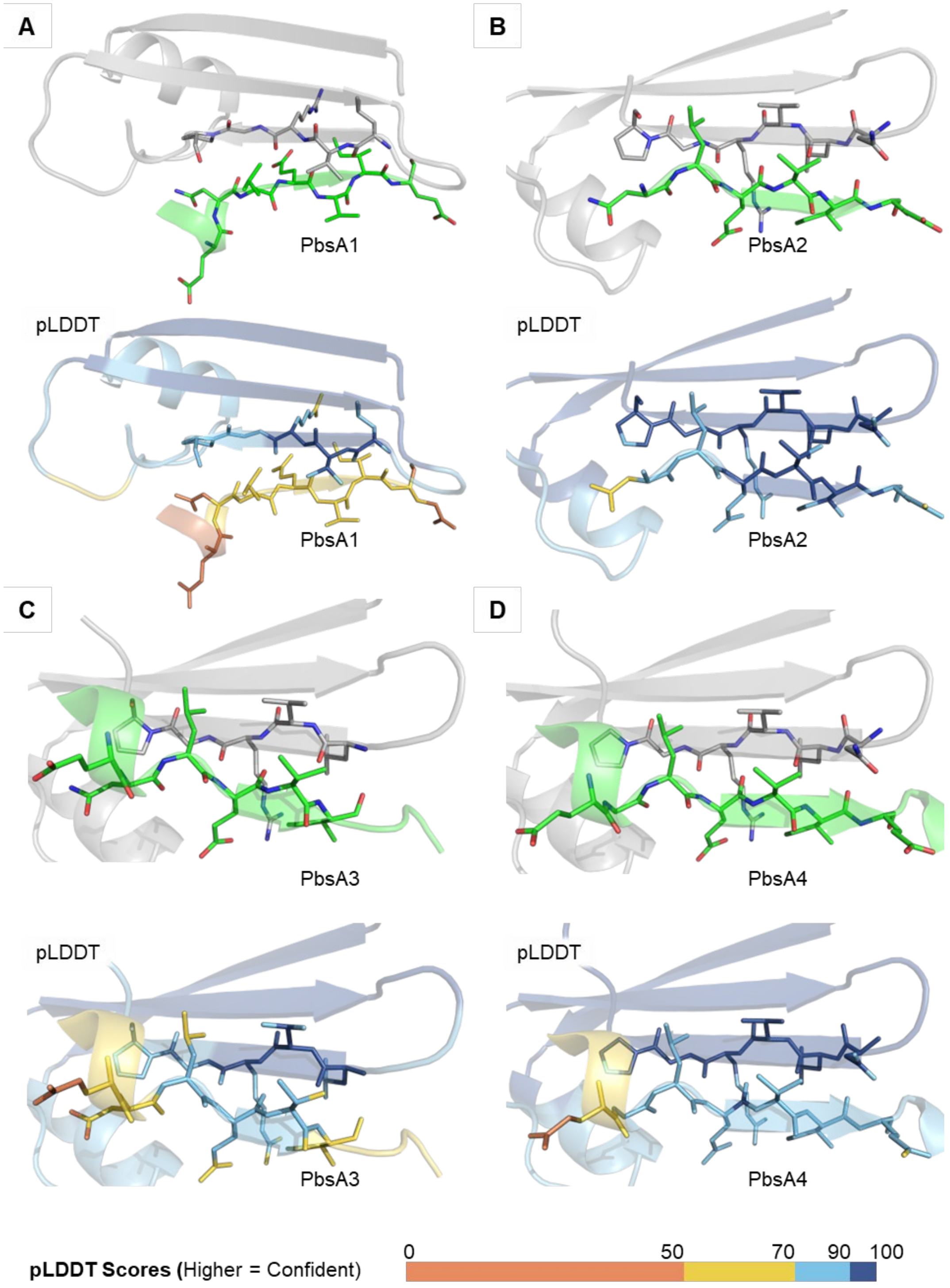


**Figure S37: AlphaFold 3-predicted interaction between PbsD and the leader peptide regions of PbsA1-A4.** Zoomed-in image of the putative anti-parallel beta-sheet interaction between PbsD (in grey) and the leader peptides of (A) PbsA1, (B) PbsA2, (C) PbsA3 and (D) PbsA4, colored by element. Hypothesized interacting residues are drawn as sticks. Confidence metrics of the AlphaFold 3-predicted structures are shown as pLDDT score-colored outputs for each panel.

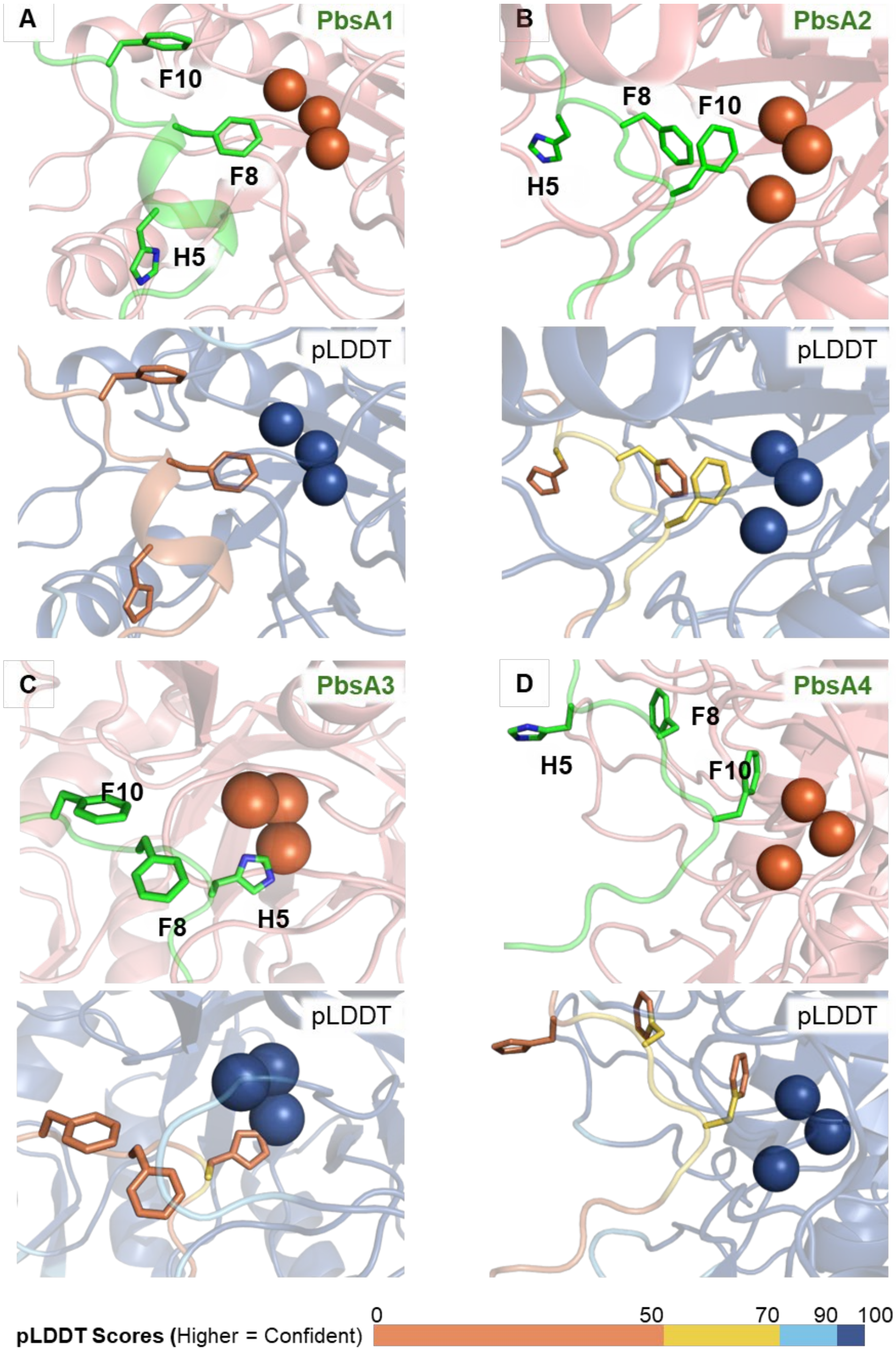


**Figure S38: Zoomed-in active site of PbsC as predicted by AlphaFold 3 in complex with PbsA1-A4.** (A) Phe8 residue of PbsA1 predicted in close proximity to the Fe-ions in the predicted active site of PbsC is displayed. (B) In the model complexed with PbsA2, the Phe10 residue was placed in proximity to the Fe ions. (C) In contrast to our experimental observations, the H5 residue was placed near the Fe ions by AlphaFold 3 while the modifiable Phe8 and Phe10 residues were further away. (D) In case of PbsA4-PbsCD complex, again the Phe10 residue was observed proximal to the active site iron. Confidence metrics of the AlphaFold 3-predicted structures are shown as pLDDT score-colored outputs for each panel.

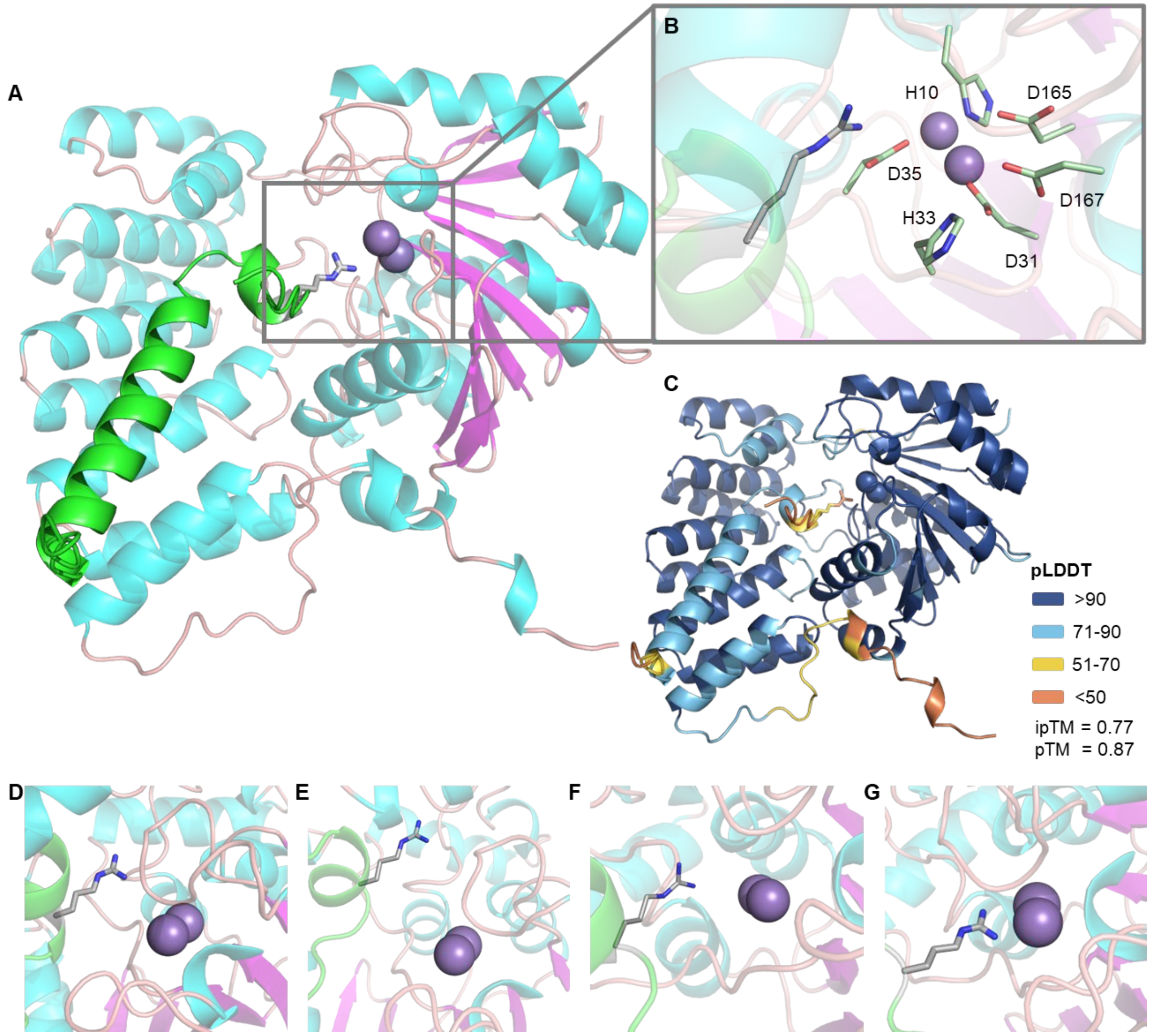


**Figure S39: AlphaFold 3-predicted model of PbsE in complex with PbsA1-A4.** An AlphaFold model was generated for the arginase PbsE in complex with the precursors PbsA1-A4. (A) The complete cartoon diagram of the PbsE-PbsA3 complex is shown. Two Mn ions were included in the prediction parameters. PbsE cartoon is colored in light blue for helices, pink for sheets and salmon for loops and disordered regions. Precursors were colored in green cartoon. The side chain of the R9 substrate residue is shown as grey sticks, colored by elements. Mn ions are depicted in purple spheres. (B) Zoomed-in image of the predicted PbsE active site showing the R9 residue in proximity to the Mn ions. Side chains of the residues hypothesized to be involved in coordinating with the Mn ions are shown in light green sticks, colored by element. (C) Confidence metrics of the AlphaFold 3-predicted structures are shown. pLDDT scores are shown as colored outputs in the cartoon diagram (higher value means higher confidence in the prediction). The predicted template modeling (pTM) score and the interface predicted template modeling (ipTM) scores measure the accuracy of the entire structure (pTM>0.5 means higher similarity of predicted and true structures; iPTM>0.8 means confident prediction). Predicted placement of the R9 residue in proximity to the Mn ions is shown for (D) PbsA1, (E) PbsA2, (F) PbsA3 and (G) PbsA4. Arg9 of PbsA4 was predicted closest to the Mn ions in comparison with other precursor complexes.

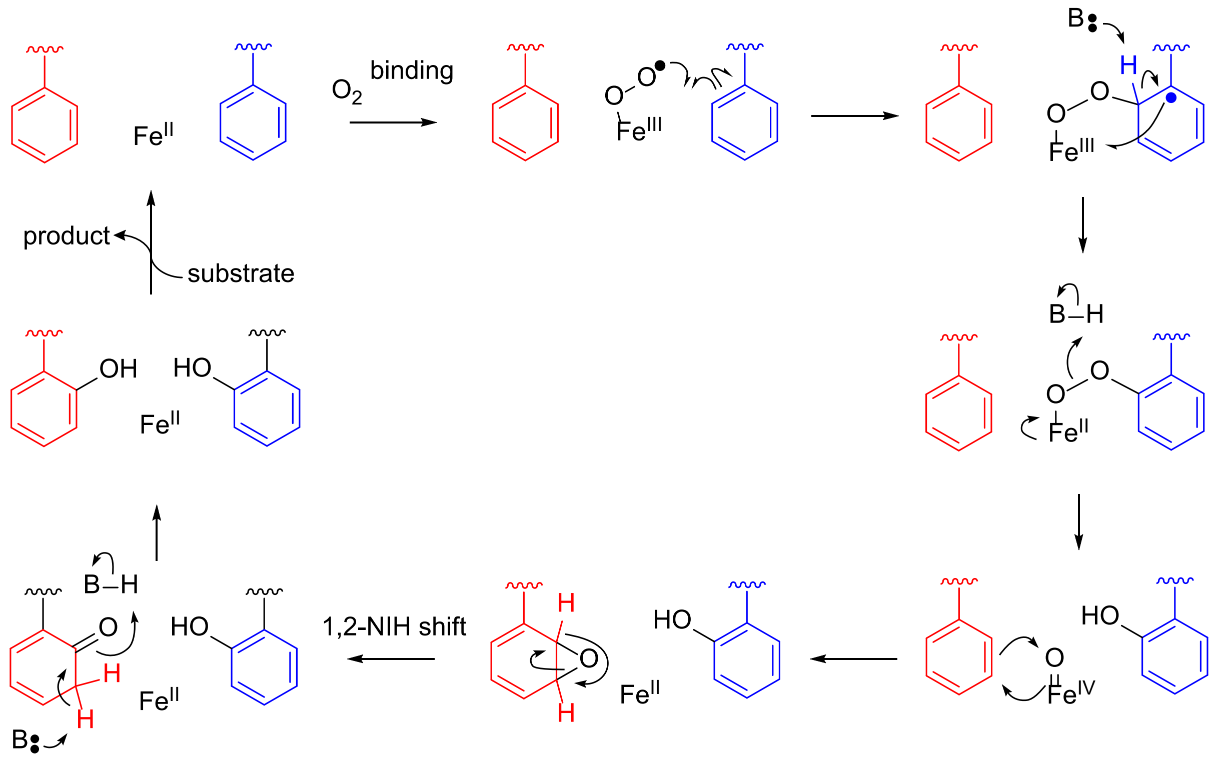


**Figure S40: Potential mechanism for a four-electron oxidation catalyzed by PbsCD.** Following oxygen binding to Fe(II), the resulting Fe(III)-superoxo could add to the phenyl ring of one phenylalanine residue (shown in blue), generating a resonance stabilized radical. Electron transfer to Fe(III) and rearomatization by deprotonation, the Fe(II)-peroxo could generate a ferryl, the second oxidizing species in the pathway. Epoxidation of the second phenylalanine residue (shown in red), followed by a hydride shift (1,2-NIH shift) and rearomatization could result in the final bis-hydroxylated product. Alternatively, the phenyl ring could attack the ferryl in an electrophilic aromatic substitution-like mechanism which after rearomatization would also give the *ortho*-Tyr.

**Table S1: ESI-HR-MS analysis of peptides used in this study.**

| Peptide | Sequence (GluC-digested core) | Charge state | Calculated mass (Da) | Observed mass (Da) | Error (Δppm) |
|---|---|---|---|---|---|
| PbsA3 Unmodified | VGSAFHAFRFNTSDTEK | $[M+3H]^{3+}$ | 638.6429 | 638.6443 | 2.19 |
| PbsA3-CD 1-OH | VGSAFHAFRFNTSDTEK | $[M+3H]^{3+}$ | 643.9756 | 643.9779 | 3.57 |
| PbsA3-CD 2-OH | VGSAFHAFRFNTSDTEK | $[M+3H]^{3+}$ | 649.3082 | 649.3123 | 6.31 |
| PbsA3-CDE | VGSAFHAFRFNTSDTEK | $[M+3H]^{3+}$ | 635.3016 | 635.3003 | -2.05 |
| PbsA3-BCDE | VGSAFHAFRFNTSDTEK | $[M+3H]^{3+}$ | 634.6315 | 634.6303 | -1.89 |
| PbsA3-YTY Unmodified | VGSAFHAYTYNTSDTEK | $[M+3H]^{3+}$ | 630.9550 | 630.9553 | 0.48 |
| PbsA3-YTY-CD 1-OH | VGSAFHAYTYNTSDTEK | $[M+3H]^{3+}$ | 636.2850 | 636.2855 | 0.79 |
| PbsA3-YTY-CD 2-OH | VGSAFHAYTYNTSDTEK | $[M+3H]^{3+}$ | 641.6183 | 641.6172 | -1.71 |
| PbsA3-YTY-B | VGSAFHAYTYNTSDTEK | $[M+3H]^{3+}$ | 630.2850 | 630.2835 | -2.38 |
| PbsA3-YTY-BCD 1-OH | VGSAFHAYTYNTSDTEK | $[M+3H]^{3+}$ | 635.6150 | 635.6157 | 1.10 |
| PbsA3-YTY-BCD 2-OH | VGSAFHAYTYNTSDTEK | $[M+3H]^{3+}$ | 640.9483 | 640.9466 | -2.65 |

**Table S2: $^{1}$H and $^{13}$C NMR assignments of the GluC-digested PbsA3-CD peptide in 90% $H_2O$ and 10% $D_2O$ at 25°C**

| Number | AA | NH | αH | βH | γH | δH | others |
|---|---|---|---|---|---|---|---|
| 1 | V | N/A | 3.76<br>58.5 | 2.13<br>29.8 | 0.93<br>17.2 | | |
| 2 | G | 8.64 | 3.98<br>42.1 | | | | |
| 3 | S | 8.22 | 4.36<br>55.2 | 3.81, 3.77<br>60.6 | | | |
| 4 | A | 8.03 | 4.07<br>49.9 | 1.19<br>16.3 | | | |
| 5 | F | 8.07 | 4.42<br>55.1 | 2.91, 2.89<br>36.9 | | 135.8 (Cγ)<br>7.11 (2H, d), 129.0<br>7.23 (2H, t), 128.8<br>7.19 (1H, t), 127.1 | |
| 6 | H | 8.05 | 4.45<br>51.5 | 3.01, 2.90<br>26.3 | | 122.5 (Cγ)<br>8.45 (H2), 133.2<br>7.05 (H4), 117,1 | |
| 7 | A | 8.317 | 4.20<br>49.9 | 1.18<br>16.3 | | | |
| 8 | F-OH | 8.15 | 4.51<br>54.0 | 3.03, 2.90<br>31.7 | | 122.9 (C4)<br>154.1 (C5-OH)<br>6.81 (H6, d), 115.4<br>7.09 (H7, t), 128.8<br>6.81 (H8, t), 120.7<br>7.05 (H9, d), 131.5 | |
| 9 | R | 7.93 | 4.10<br>53.1 | 1.54<br>27.8 | 1.29<br>23.8 | 2.99<br>40.3 | NHε: 6.99 (t)<br>156.6 (Cζ) |
| 10 | F-OH | 8.154 | 4.50<br>53.9 | 2.91<br>31.6 | | 122.7 (C4)<br>154.1 (C5-OH)<br>6.81 (H6, d), 115.4<br>7.08 (H7, t), 128.7<br>6.73 (H8, t), 120.6<br>6.93 (H9, d), 131.4 | |
| 11 | N | 8.328 | 4.66<br>50.4 | 2.77, 2.68<br>36.5 | | | 7.46<br>6.79 |
| 12 | T | 7.99 | 4.23<br>59.3 | 4.19<br>66.8 | 1.09<br>18.8 | | |
| 13 | S | 8.28 | 4.34<br>55.3 | 3.77, 3.70<br>61.0 | | | |
| 14 | D | 8.17 | 4.62<br>50.0 | 2.70, 2.61<br>36.0 | | | |
| 15 | T | 8.00 | 4.23<br>59.3 | 4.15<br>66.9 | 1.09<br>18.8 | | |
| 16 | E | 8.14 | 4.26<br>53.1 | 2.02, 1.89<br>25.5 | 2.35,<br>30.0 | | |
| 17 | K | 8.04 | 4.14<br>54.0 | 1.76, 1.63<br>30.3 | 1.54<br>26.2 | 1.30<br>21.8 | 2.88, 38.6<br>7.41 (br, NZ) |

**Table S3: $^{1}$H and $^{13}$C NMR assignments of the GluC-digested PbsA3-CDE peptide in 90% $H_2O$ and 10% $D_2O$ at 25ºC**

| Number | AA | NH | αH | βH | γH | δH | others |
|---|---|---|---|---|---|---|---|
| 1 | V | N/A | 3.77<br>58.5 | 2.13<br>29.8 | 0.93<br>17.1 | | |
| 2 | G | 8.64 | 3.98<br>42.1 | | | | |
| 3 | S | 8.24 | 4.37<br>55.5 | 3.81, 3.77<br>60.8 | | | |
| 4 | A | 8.03 | 4.08<br>49.9 | 1.19<br>16.3 | | | |
| 5 | F | 8.06 | 4.39<br>55.1 | 2.90<br>37.0 | | 7.07 (2H, d), 129.1<br>7.20 (2H, t), 128.7<br>7.17(1H, t), 127.4 | |
| 6 | H | 8.13 | 4.45<br>51.7 | 3.01, 2.91<br>26.3 | | 8.43, 133.0<br>7.03, 116.9 | |
| 7 | A | 8.30 | 4.20<br>49.9 | 1.18<br>16.3 | | | |
| 8 | F-OH | 8.14 | 4.44<br>54.3 | 3.02, 2.91<br>31.8 | | 6.78 (H6, d), 115.4<br>7.04 (H7, t), 128.8<br>6.79 (H8, t), 120.7<br>7.03 (H9, d), 131.7 | |
| 9 | Orn | 7.99 | 4.16<br>53.0 | 1.62, 1.55<br>27.7 | 1.47<br>22.9 | 2.85<br>38.7 | $-NH_3^+$ (3ºC)<br>7.51 |
| 10 | F-OH | 8.14 | 4.43<br>54.2 | 2.87<br>32.0 | | 6.78 (H6, d), 115.4<br>7.04 (H7, t), 128.7<br>6.67 (H8, t), 120.7<br>6.86 (H9, d), 131.6 | |
| 11 | N | 8.22 | 4.66<br>50.7 | 2.69, 2.60<br>36.2 | | | 7.45<br>6.79 |
| 12 | T | 7.95 | 4.23<br>59.1 | 4.15<br>66.8 | 1.09<br>18.8 | | |
| 13 | S | 8.29 | 4.34<br>55.3 | 3.75, 3.70<br>61.2 | | | |
| 14 | D | 8.29 | 4.63<br>50.0 | 2.65, 2.57<br>36.2 | | | |
| 15 | T | 8.00 | 4.24<br>59.1 | 4.18<br>66.8 | 1.09<br>18.8 | | |
| 16 | E | 8.15 | 4.23<br>53.2 | 1.98, 1.86<br>26.1 | 2.22<br>30.5 | | |
| 17 | K | 7.82 | 4.07<br>53.8 | 1.73, 1.63,<br>30.4 | 1.54<br>26.4 | 1.28<br>21.9 | 2.89<br>39.1<br>7.40 br (NZ) |

Orn = Ornithine

**Table S4: $^1$H and $^{13}$C NMR assignments of GluC-digested PbsA3-BCDE in 90% $H_2O$ and 10% $D_2O$ at 25°C**

| Number | AA | NH<br>C=O | αH<br>Cα | βH<br>Cβ | γH<br>Cγ | δH<br>Cδ | others |
|---|---|---|---|---|---|---|---|
| 1 | V | N/A<br>170.1 | 3.76<br>58.7 | 2.12<br>29.9 | 0.93<br>17.3 | | |
| 2 | G | 8.64 | 3.97<br>42.1 | | | | |
| 3 | S | 8.27 | 4.34<br>55.3 | 3.73<br>61.2 | | | |
| 4 | A | 8.28 | 4.19<br>49.6 | 1.17<br>16.3 | | | |
| 5 | F | 8.05<br>172.5 | 4.33<br>55.2 | 2.87, 2.82<br>36.9 | | 135.8 (Cγ)<br>7.05 (2H, d), 129.0<br>7.20 (2H, t), 128.7<br>7.20 (1H, t), 128.8 | |
| 6 | H | 7.83<br>170.7 | 4.41<br>51.0 | 2.79, 2.60<br>26.8 | | 127.3 (C)<br>8.19 (H2), 132.5<br>6.95 (H4), 117.5 | |
| 7 | A | 8.38 | 4.06<br>49.9 | 1.31<br>16.1 | | | |
| 8 | F-OH | 7.27 | 4.67<br>54.1 | 3.46, 2.85<br>30.3 | | 131.3 (C8)<br>6.89 (s, H9), 129.7<br>122.3 (C4)<br>153.4 (C5-OH)<br>6.80 (d, H6), 116.1<br>7.17 (d, H7), 125.2 | |
| 9 | R (Orn) | 8.86 | 4.57<br>52.3 | 1.87, 1.74<br>29.2 | 1.59<br>26.4 | 2.95<br>38.7 | $NH_3^+$ (br), 7.51 |
| 10 | F-OH | 9.11 | 4.81<br>53.0 | 3.21, 2.82<br>28.7 | | 131.0 (C8)<br>7.16 (s, H9), 125.4<br>124.8 (C4)<br>152.6 (C5-OH)<br>6.77 (d, H6), 116.0<br>7.12 (d, H7), 124.6 | |
| 11 | N | 8.69 | 4.76<br>50.4 | 2.81, 2.74<br>36.2 | | | 7.56 (HD21)<br>6.87 (HD22) |
| 12 | T | 8.21<br>170.9 | 4.31<br>59.1 | 4.25<br>66.8 | 1.16<br>18.9 | | |
| 13 | S | 8.29<br>174.4 | 4.40<br>55.8 | 3.81<br>60.8 | | | |
| 14 | D | 8.35 | 4.66<br>50.6 | 2.78, 2.71<br>36.1 | | | |
| 15 | T | 8.01 | 4.22<br>59.1 | 4.16<br>66.8 | 1.10<br>18.8 | | |
| 16 | E | 8.14 | 4.28<br>53.3 | 2.02, 1.85<br>26.2 | 2.36<br>30.4 | 177.9 | |
| 17 | K | 8.00 | 4.11<br>54.0 | 1.75, 1.63<br>30.3 | 1.56<br>26.1 | 1.29<br>22.1 | 2.87 (He),<br>39.2<br>7.40 br ($NH_3^+$) |

Note: Orn = ornithine

**Table S5: $^{1}H$ and $^{13}C$ NMR assignments of the GluC-digested YTY-B peptide in 90% $H_2O$, 10% $D_2O$ and 0.1% $d_2$-formic acid at 25°C.**

| Number | AA | NH<br>C=O | αH<br>Cα | βH<br>Cβ | γH<br>Cγ | δH<br>Cδ | | others |
|---|---|---|---|---|---|---|---|---|
| 1 | V | N/A<br>170.0 | 3.77<br>58.8 | 2.13<br>29.9 | 0.93<br>17.2 | | | |
| 2 | G | 8.64<br>171.0 | 3.96<br>42.3 | | | | | |
| 3 | S | 8.28<br>171.5 | 4.35<br>55.4 | 3.76, 3.72<br>61.3 | | | | |
| 4 | A | 8.30<br>174.3 | 4.20<br>49.7 | 1.19<br>16.4 | | | | |
| 5 | F | 8.07<br>172.4 | 4.43<br>55.1 | 2.92<br>36.9 | | 135.8 (Cγ)<br>7.11 (2H, d), 129.0<br>7.24 (2H, t), 128.9<br>7.21 (1H, t), 127.4 | | |
| 6 | H | 8.11<br>172.5 | 4.53<br>51.7 | 3.02, 2.91<br>26.8 | | 128.0 (C)<br>8.47 (H2), 133.4 (222Hz)<br>7.11 (H4), 117.4 (201Hz) | | |
| 7 | A | 8.31<br>174.0 | 4.28<br>49.7 | 1.31<br>16.5 | | | | |
| 8 | Y | 7.67<br>(br)<br>172.3 | 4.76<br>53.8 | 3.07, 2.97<br>36.4 | | 130.2 (Cγ)<br>7.02 (Hδ1, d), 130.6<br>6.64 (Hδ2, s), 133.3<br>6.81 (Hε1, d), 115.8<br>151.8 (Cζ-OH)<br>125.4 (Cε2-linker) | | |
| 9 | T | 8.57 | 4.55<br>58.9 | 3.95<br>68.1 | 1.08<br>19.0 | | | |
| 10 | Y | 8.90<br>172.4 | 4.54<br>54.5 | 2.87, 2.76<br>36.8 | | 130.4 (Cγ)<br>7.077 (Hδ1, d), 130.4<br>6.85 (Hδ2, s), 130.6<br>6.802 (Hε1, d), 115.8<br>151.7 (Cζ -OH)<br>125.4 (Cε2-linker) | | |
| 11 | N | 8.63<br>1`75.5 | 4.80<br>50.4 | 2.78, 2.72<br>36.5 | | 7.52 (HD21)<br>6.83 (HD22) | | |
| 12 | T | 8.25<br>170.9 | 4.34<br>59.3 | 4.26<br>67.0 | 1.16<br>18.7 | | | |
| 13 | S | 8.32<br>171.4 | 4.41<br>55.8 | 3.84, 3.78<br>61.1 | | | | |
| 14 | D | 8.36<br>172.9 | 4.67<br>51.1 | 2.79, 2.73<br>36.5 | | | | |
| 15 | T | 8.01<br>173.2 | 4.24<br>59.2 | 4.16<br>66.9 | 1.11<br>18.8 | | | |
| 16 | E | 8.17<br>172.4 | 4.28<br>53.3 | 2.03, 1.90<br>26.2 | 2.35<br>30.7 | 178.0 | | |
| 17 | K | 7.99<br>177.4 | 4.13<br>54.2 | 1.75, 1.63<br>30.5 | 1.57<br>26.3 | 1.31<br>21.9 | 2.89 (Hε),<br>39.4 | 7.41 ($NH_3^+$)<br>(br) |

**Table S6: $^{1}$H and $^{13}$C NMR assignments of the GluC-digested mono- and bis-hydroxylated YTY-CD in 90%$H_2O$ and 10%$D_2O$ at 25 °C.** For residues having 2 sets of chemical shifts (residue 6 to 12), the top is for mono and the bottom is for bis-hydroxylated YTY-CD peptide.

| Number | AA | NH<br>C=O | αH<br>Cα | βH<br>Cβ | γH<br>Cγ | δH<br>Cδ | others |
|---|---|---|---|---|---|---|---|
| 1 | V | N/A<br>170.2 | 3.77<br>58.8 | 2.13<br>29.8 | 0.94<br>17.2 | | |
| 2 | G | 8.65 (br)<br>170.7 | 3.98<br>42.2 | | | | |
| 3 | S | 8.29<br>171.7 | 4.36<br>55.5 | 3.78, 3.72<br>61.3 | | | |
| 4 | A | 8.30<br>174.5 | 4.20<br>49.7 | 1.19<br>16.3 | | | |
| 5 | F | 8.06<br>172.6 | 4.40<br>55.5 | 2.90<br>36.8 | | 135.9 (Cγ)<br>7.08, 129.2<br>7.21, 128.8<br>7.18, 127.3 | |
| 6 | H | 8.05<br><br>170.7 | 4.46<br>52.0<br><br>4.42<br>52.0 | 2.98,2.88<br>26.8<br><br>2.95,2.87<br>26.7 | | 128.0 (C)<br>(8.34, 133.7), (6.99, 117.2)<br><br>(8.35, 133.7), (6.97, 117.2) | |
| 7 | A | 8.09<br>174.6<br><br>8.12<br>174.55 | 4.11<br>49.7<br><br>4.10<br>49.7 | 1.21<br>16.3<br><br>1.22<br>16.3 | | | |
| 8 | Y | 8.11<br>173.0<br><br>8.12<br>173.0 | 4.49<br>54.7<br><br>4.51<br>54.8 | 2.88, 2.83<br>36.3<br><br>2.91, 2.83<br>31.2 | | 127.9 (Cγ)<br>6.69 (Hε, d), 115.6<br>6.97 (Hδ, d), 130.6<br>154.5 (Cζ)<br><br>114.7 (Cγ)<br>6.80 (Hδ2, d), 132.5<br>6.25(Hε2, dd), 107.6<br>155.7 (Cζ)<br>6.30(Hε1, s), 102.9<br>155.3 (Cδ1) | |
| 9 | T | 7.86<br><br>7.76 | 4.14<br>59.1<br>4.12<br>59.1 | 4.00<br>66.9<br>4.02<br>66.9 | 1.01<br>18.7<br>0.96<br>18.6 | | |
| 10 | Y | 8.04<br>173.0<br><br>7.96<br>173.0 | 4.46<br>54.5<br><br>4.46<br>54.5 | 2.96, 2.85<br>31.2<br><br>2.95, 2.86<br>31.2 | | 114.7 (Cγ)<br>6.87(Hδ2, d), 132.4<br>6.28(Hε2, dd), 107.6<br>155.7 (Cζ)<br>6.27(Hε1, s), 102.9<br>155.2 (Cδ1)<br><br>114.7 (Cγ)<br>6.85(Hδ2, d), 132.4<br>6.26(Hε2, dd), 107.6<br>155.7 (Cζ)<br>6.27(Hε1, s), 102.9<br>155.3 (Cδ1) | |
| 11 | N | 8.16 | 4.65 | 2.70, 2.61 | | 7.47, 6.79 | |

| | | | | | | | | |
|---|---|---|---|---|---|---|---|---|
| | | <br><br>8.12<br>174.3 | 50.4<br><br>4.65<br>50.4 | 36.5<br><br>2.70, 2.60<br>36.5 | | <br>7.45, 6.78<br>172.4 | | |
| 12 | T | 8.01<br>170.9<br><br>7.99<br>172.2 | 4.24<br>59.4<br><br>4.25<br>59.4 | 4.20<br>67.0<br><br>4.17<br>67.1 | 1.11<br>18.7<br><br>1.11<br>18.7 | | | |
| 13 | S | 8.26<br>171.7 | 4.40<br>55.6 | 3.81, 3.77<br>61.1 | | | | |
| 14 | D | 8.28<br>173.8 | 4.58<br>51.1 | 2.66, 2.58<br>38.4 | | 177.3 | | |
| 15 | T | 7.94<br>173.2 | 4.21<br>59.3 | 4.17<br>67.1 | 1.11<br>18.7 | | | |
| 16 | E | 8.196<br>172.8 | 4.22<br>53.9 | 2.00, 1.86<br>27.2 | 2.20<br>33.3 | 181.2 | | |
| 17 | K | 7.807<br>178.4 | 4.07<br>54.8 | 1.73, 1.62<br>31.0 | 1.57<br>26.3 | 1.28<br>21.8 | 2.89 (He),<br>39.4 | 7.41 ($NH_3^+$)<br>(br) |

**Table S7: Plasmids, primers and gblocks used in this study.**

Listed are plasmid constructs, vector backbones, primers and synthesized codon optimized DNA sequences used to amplify genes encoding precursors and modifying enzymes, introduced restriction sites for cloning, and the translated protein sequence of expressed genes. Precursor sequences are shown in green font. TEV protease cleavage sites are in red font. Maturase modification sites in the precursors are displayed in bold; mutation sites are underlined. N-terminal His6 tags and carried over amino acid sequences from the vector backbone due to the fusion are shown in purple font.

| Plasmid Construct | Vector Backbone | | 5' → 3' sequence | Restriction Sites |
|---|---|---|---|---|
| **NHis$_6$-PbsA1** | pET-Duet-1 | ORF | ATGGGCAGCAGCCATCACCATCATCACCACAGCCAGGATCCGATGACGAAGCAGGTAGAGAAAACACAGGCC<br>GAAAAATTAGTTGAAAACTTGGAAGTAATTGAAGCTAAGGAACTTGAGGTTGGTAGCGCATTTCATGCCTTTCGT<br>TTCTCCACAGTGTCTGACACGGAAAAGTAA | BamHI<br>EcoRI |
| Translated sequence | MGSSHHHHHHSQDPMTKQVEKTQAEKLVENLEVIEAKELEVGSAFHA**FRF**STVSDTEK | | | |
| **NHis$_6$-PbsA2** | pET-Duet-1 | ORF | ATGGGCAGCAGCCATCACCATCATCACCACAGCCAGGATCCGATGACTAAGCAAGTCGAAAAGACTGAAGTTG<br>AGAAGCTGGTTGAAAATCTGGAAGTGATTGAGGCGAAGGAACTTGAAGTTGGGTCAGCTTTCCACGCTTTCCG<br>TTTTTCGACTGTTTCCGACACGGAAAAGTAA | BamHI<br>EcoRI |
| Translated sequence | MGSSHHHHHHSQDPMTKQVEKTEVEKLVENLEVIEAKELEVGSAFHA**FRF**STVSDTEK | | | |
| **NHis$_6$-PbsA3** | pET-Duet-1 | ORF | ATGGGCAGCAGCCATCACCATCATCACCACAGCCAGGATCCGATGACCAAACAAGTCGAGAAGACTGAGGCG<br>GAGAAGTTGGTCGAGAACTTAGAAGTCATCGAGGCCAAGGAACTGGAAGTAGGTTCTGCCTTTCACGCCTTTC<br>GTTTCAACACGAGCGATACGGAAAAATAA | BamHI<br>EcoRI |
| Translated sequence | MGSSHHHHHHSQDPMTKQVEKTEAEKLVENLEVIEAKELEVGSAFHA**FRF**NTSDTEK | | | |
| **NHis$_6$-PbsA4** | pET-Duet-1 | ORF | ATGGGCAGCAGCCATCACCATCATCACCACAGCCAGGATCCGATGACGAAACAAGTCGAGAAGACAGAGGCT<br>GAAAAACTTGTTGAGAACCTTGAGGTCATCGAAGCGAAGGAATTGGAGGTTGGCAGCGCATTTCATGCTTTCC<br>GTTTTAGTGCAGTAACCGGTGCTGAAAAGTAA | BamHI<br>EcoRI |
| Translated sequence | MGSSHHHHHHSQDPMTKQVEKTEAEKLVENLEVIEAKELEVGSAFHA**FRF**SAVTGAEK | | | |
| **NHis$_6$-PbsA3-ARF** | pET-Duet-1 | ORF | ATGGGCAGCAGCCATCACCATCATCACCACAGCCAGGATCCGATGACCAAACAAGTCGAGAAGACTGAGGCG<br>GAGAAGTTGGTCGAGAACTTAGAAGTCATCGAGGCCAAGGAACTGGAAGTAGGTTCTGCCTTTCACGCCGCAC<br>GTTTTAACACGAGCGATACGGAAAAATAA | BamHI<br>EcoRI |
| Translated sequence | MGSSHHHHHHSQDPMTKQVEKTQAEKLVENLEVIEAKELEVGSAFHAAR**F**STVSDTEK | | | |
| **NHis$_6$-PbsA3-FRA** | pET-Duet-1 | ORF | ATGGGCAGCAGCCATCACCATCATCACCACAGCCAGGATCCGATGACCAAACAAGTCGAGAAGACTGAGGCG<br>GAGAAGTTGGTCGAGAACTTAGAAGTCATCGAGGCCAAGGAACTGGAAGTAGGTTCTGCCTTTCACGCCTTTC<br>GTGCAAACACGAGCGATACGGAAAAATAA | BamHI<br>EcoRI |
| Translated sequence | MGSSHHHHHHSQDPMTKQVEKTEVEKLVENLEVIEAKELEVGSAFHA**F**RASTVSDTEK | | | |
| **NHis$_6$-PbsA3-ARA** | pET-Duet-1 | ORF | ATGGGCAGCAGCCATCACCATCATCACCACAGCCAGGATCCGATGACCAAACAAGTCGAGAAGACTGAGGCG<br>GAGAAGTTGGTCGAGAACTTAGAAGTCATCGAGGCCAAGGAACTGGAAGTAGGTTCTGCCTTTCACGCCGCAC<br>GTGCAAACACGAGCGATACGGAAAAATAA | BamHI<br>EcoRI |
| Translated sequence | MGSSHHHHHHSQDPMTKQVEKTEAEKLVENLEVIEAKELEVGSAFHAARANTSDTEK | | | |
| **NHis$_6$-PbsA3-FRY** | pET-Duet-1 | ORF | ATGGGCAGCAGCCATCACCATCATCACCACAGCCAGGATCCGATGACCAAACAAGTCGAGAAGACTGAGGCG<br>GAGAAGTTGGTCGAGAACTTAGAAGTCATCGAGGCCAAGGAACTGGAAGTAGGTTCTGCCTTTCACGCCTTTC<br>GTTATAACACGAGCGATACGGAAAAATAA | BamHI<br>EcoRI |
| Translated sequence | MGSSHHHHHHSQDPMTKQVEKTEAEKLVENLEVIEAKELEVGSAFHA**F**R**Y**NTSDTEK | | | |
| **NHis$_6$-PbsA3-YRF** | pET-Duet-1 | ORF | ATGGGCAGCAGCCATCACCATCATCACCACAGCCAGGATCCGATGACCAAACAAGTCGAGAAGACTGAGGCG<br>GAGAAGTTGGTCGAGAACTTAGAAGTCATCGAGGCCAAGGAACTGGAAGTAGGTTCTGCCTTTCACGCCTATC<br>GTTTCAACACGAGCGATACGGAAAAATAA | BamHI<br>EcoRI |
| Translated sequence | MGSSHHHHHHSQDPMTKQVEKTQAEKLVENLEVIEAKELEVGSAFHA**Y**R**F**STVSDTEK | | | |
| **NHis$_6$-PbsA3-YRY** | pET-Duet-1 | ORF | ATGGGCAGCAGCCATCACCATCATCACCACAGCCAGGATCCGATGACCAAACAAGTCGAGAAGACTGAGGCG<br>GAGAAGTTGGTCGAGAACTTAGAAGTCATCGAGGCCAAGGAACTGGAAGTAGGTTCTGCCTTTCACGCCTATC<br>GTTATAACACGAGCGATACGGAAAAATAA | BamHI<br>EcoRI |
| Translated sequence | MGSSHHHHHHSQDPMTKQVEKTEVEKLVENLEVIEAKELEVGSAFHA**Y**R**Y**STVSDTEK | | | |
| **NHis$_6$-PbsA3-F5A** | pET-Duet-1 | ORF | ATGGGCAGCAGCCATCACCATCATCACCACAGCCAGGATCCGATGACCAAACAAGTCGAGAAGACTGAGGCG<br>GAGAAGTTGGTCGAGAACTTAGAAGTCATCGAGGCCAAGGAACTGGAAGTAGGTTCTGCCGCGCACGCCTTTC<br>GTTTCAACACGAGCGATACGGAAAAATAA | BamHI<br>EcoRI |
| Translated sequence | MGSSHHHHHHSQDPMTKQVEKTEAEKLVENLEVIEAKELEVGSAAHA**F**R**F**NTSDTEK | | | |
| **NHis$_6$-PbsA3-H6A** | pET-Duet-1 | ORF | ATGGGCAGCAGCCATCACCATCATCACCACAGCCAGGATCCGATGACCAAACAAGTCGAGAAGACTGAGGCG<br>GAGAAGTTGGTCGAGAACTTAGAAGTCATCGAGGCCAAGGAACTGGAAGTAGGTTCTGCCTTTGCGGCCTTTC<br>GTTTCAACACGAGCGATACGGAAAAATAA | BamHI<br>EcoRI |
| Translated sequence | MGSSHHHHHHSQDPMTKQVEKTEAEKLVENLEVIEAKELEVGSAFAA**F**R**F**NTSDTEK | | | |
| **NHis$_6$-PbsA3-YTY** | pET-Duet-1 | ORF | ATGGGCAGCAGCCATCACCATCATCACCACAGCCAGGATCCGATGACCAAACAAGTCGAGAAGACTGAGGCG<br>GAGAAGTTGGTCGAGAACTTAGAAGTCATCGAGGCCAAGGAACTGGAAGTAGGTTCTGCCTTTCACGCCTATA<br>CGTATAACACGAGCGATACGGAAAAATAA | BamHI<br>EcoRI |
| Translated sequence | MGSSHHHHHHSQDPMTKQVEKTEAEKLVENLEVIEAKELEVGSAFHA**YTY**NTSDTEK | | | |
| **NHis$_6$-StgA** | pET-Duet-1 | ORF | ATGGGCAGCAGCCATCACCATCATCACCACAGCCAGGATCCGATGACCGCTGCTAGCCCGGAGAAACTCGCG<br>GATGAAGAATTTGGTGAGTTGGAAGAGCTGCAAGTCATTGATAGCAGTGAACTTAAATTAGGTGCCGCATATCA<br>CACATATACTTACATAGCTGCCGAGTCAACAGATGATATACCCGATTAA | BamHI<br>EcoRI |
| Translated sequence | MGSSHHHHHHSQDPMTAASPEKLADEEFGELEELQVIDSSELKLGAAYHT**Y**T**Y**IAAESTDDIPD | | | |
| **NHis$_6$-TbaA** | pET-Duet-1 | ORF | | BamHI |

| | | | | |
|---|---|---|---|---|
| | | | ATGGGCAGCAGCCATCACCATCATCACCACAGCCAGGATCCGATGAACCATGATCTTGACCAACTGGAGATCG<br>TTGAGGCTGAGGACTTGCAAGTAGGACACGCATTCCATACATTTATGTTTCAATCCCATCTGGACGAGTCCGAG<br>GAAGTATGA | EcoRI |
| Translated sequence | MGSSHHHHHHSQDPMNHDLDQLEIVEAEDLQVGHAFHT**F**M**F**QSHLDESEEV | | | |
| **NHis₆-PbsC** | pRSF-Duet | ORF | ATGGGCAGCAGCCATCATCATCATCATCACAGCAGCGGCGAGAACCTGTACTTCCAAAGTATGTCAACCGTTTA<br>CCAAAAAATTAAATTTCCTGATATTAAGGAAAAATTAGGACTGGGCTTAGGAATGGACTTACCGTGGGGAGAGG<br>ACATTGGGTTCTCGTCAAAAGGCGACCATGATGACATCACGCTGAAAATGAAGGGATTCTTTCAAAATTACAAA<br>GATGAGTTCAACTATATCTTTTTCGCCTTTCAGCCTAAAAATCGCTCCGTGCTTAAAGCGGAAGACTATTTTGAC<br>GCCTACGACCGCTTGTTTGAAGCGAACCCTCACTTAAAGGCACGTGCATTCCATCAGACCATCTTGAATATGGG<br>GGCGACCGAGTACTACAAAAAGTCAGAGATCATTGAGTTCACAAACAAGATTATCGAGCGTTATGATATTCGTT<br>GGATTGTCGAAGATCTTGGACTGTGGTCGATTAAGGGTAAAACGGTACCCTTTCCTCTTCCACCCTATATGACA<br>CCTACTGGACTTGAGGCCTGTATCAAAAATATCAATGAATATCAAAGCGAATTAGCCGTTCCTGTTTCAGTCGA<br>GTTCCCTGGCTTTACAGAGGGTACGAACTTCTTTATTGGCGAGATCAATGGGTTTGATTACTTTCGCACATTGG<br>TAGAGGAAACTAATAGTCCCATTACCATTGACATCGGACACATTCTTTCGTATCAGTGGTTACTTGGGAATACTA<br>ATGAAAAAATGTTCAACGGTTTGGAAAAATTACCGTTCGAAAATTGCTTTGAGCTGCATCTTTCGGGCTGCCAG<br>ATCATCAAAGGAAAATTCCGTGACCTGCATCATGGAGTTTTAATGGACGAACAGATCCAACTTTTGGATTATCTG<br>CTGCCGTTGTGTCCAAACCTGAAAGCCATTACTTACGAGGATCCTAAGTATACGGACGAGGGAATCTTGATCCC<br>TAAGAGCCAAAAAAATTACCAACGCATGAAAGAGGCGGTTGAGAAATGGCAGATGATTTAA | NdeI<br>XhoI |
| Translated sequence | MGSSHHHHHHSSGENLYFQSMSTVYQKIKFPDIKEKLGLGLGMDLPWGEDIGFSSKGDHDDITLKMKGFFQNYKDEFNYIFFAFQPKNRSVLKAEDYFDAYDRLF<br>EANPHLKARAFHQTILNMGATEYYKKSEIIEFTNKIIERYDIRWIVEDLGLWSIKGKTVPFPLPPYMTPTGLEACIKNINEYQSELAVPVSVEFPGFTEGTNFFIGEING<br>FDYFRTLVEETNSPITIDIGHILSYQWLLGNTNEKMFNGLEKLPFENCFELHLSGCQIIKGKFRDLHHGVLMDEQIQLLDYLLPLCPNLKAITYEDPKYTDEGILIPKSQ<br>KNYQRMKEAVEKWQMI | | | |
| **PbsD** | pRSF-Duet | ORF | ATGGCAAATGATCTGAAACCTAAAAACTACAGTTTGGATGAACTGCTGTACAACTTACTGTACGATTATCAATAT<br>CGCAACAATTTTCTTAATGACGAGTTAAACGAGCTTAATTTGTCCGCCGACAACTTGAACCACATTAAAACAATT<br>GACAAGGAGGAATTAGTAGCCACCGCGACTACTATTGTCCGCAACTTGATGTCCGGCAACATTGAACACAACG<br>GAGGGCTGCGTACTTCGTTTCCTGGGGTTTTTCAGGCGTTAGAGTTTCGTAAGACTGACATTACCCTGCTTATG<br>CACAAATTTCTTGCTTCCAAGCACTTTGAGGGTTACATGGAGCTTCCGTATGCCGGCGAAGGGACTTGCATTGA<br>GGAGGCCTTCTACAACTACCTTTCAGAAAATGAAGAATTTATTTTGGCAGCGGACAATAATCACCTTTTACTTAA<br>GCACGAGTTCTTGAATGCTATCCTGAGCATTTTGACAGTGAACAAATACCCTTTCTTCCGTATCGACAGCGACTT<br>AGTAAAAAATAACGGCCACATCTACTATGCCTACCAGACTTACAGCAAGGAAATTAGCGAAGCACTGTCGGGAA<br>AAAAAGTGGCTGGTGACTCCGAGATGGTTATCTGGCTGTACGCTGCCACTGAAAAGAACTTAGTCCGTGGCCC<br>AATTCATCCAAGCATCTTGGAGATGGTAGAAATCGGAAGCTCACGCGAAATTTCTCGCCAACAGAAATTCAACC<br>AAGATCAAATTGATTGGATCTTGAACTTAGGTTTAATTAAGAATGACGGCTAA | NdeI<br>XhoI |
| Translated sequence | MANDLKPKNYSLDELLYNLLYDYQYRNNFLNDELNELNLSADNLNHIKTIDKEELVATATTIVRNLMSGNIEHNGGLRTSFPGVFQALEFRKTDITL<br>LMHKFLASKHFEGYMELPYAGEGTCIEEAFYNYLSENEEFILAADNNHLLLKHEFLNAILSILTVNKYPFFRIDSDLVKNNGHIYYAYQTYSKEISEA<br>LSGKKVAGDSEMVIWLYAATEKNLVRGPIHPSILEMVEIGSSREISRQQKFNQDQIDWILNLGLIKNDG | | | |
| **NHis₆-PbsE** | pCDF-Duet | ORF | ATGGGCAGCAGCCATCATCATCATCATCACAGCAGCGGCGAGAACCTGTACTTCCAAAGTATGGATAAGATCTA<br>CGTTTATGAGGAGCATTCTGAAGTATTCAGCTACTGGGTGAACGAGATTCCTCGTAACTCCACGCTGGTATACT<br>TTGATCAACACCTGGACTTAAAGTTGATCGAGAATTCAAAAATGAAACGTATTTCGCAGTTCATCAAGGAAGGC<br>CTTTGTATTGACGAGTTAAAAAAAGACGTCCCCTGTCGCGAGGATGGACGTTATTGCTACGGTATCGACGACTT<br>TTTATATGCTGCCATCCAGGAAGGATTGTTTCGTAAAATTATCTGGGTTTATCCACACATGCTGGGCGAAAAAG<br>GATTCTCTGAGTTGCTTTGGGGGTTATTGTCACTGATCCCCAATCATGGAAAAGAGTTTATTTCATCTTTTCGTA<br>ACGGGAAAAACTCCGCTTCGGTACAATTGAACAACATCGAGTTGCACGTAACAAGTATTGATTACTTGGGAGAA<br>TTTATCATTAATGAGCAGGTGATTGTAGACATTGATCTTGATTACTTTTATGACCCGAAGACCAACGACCTGAGT<br>AACGACATTGACGAGACGTTAAACGTCTTGGGCAACCTGGGGATTATCAATCAAGTTAAGACTATGACGTACTC<br>AATTAAGTCGGGGTTTTTACCGGAATCATTTCGTTGGTTGGGTGAATACATCGCGGGGCAGATGGGCCGCGAG<br>ATCGTTTACTCGAAGGTAGATAAAATTAGTCCTAAGGTGACTATGGAAAAAATCTCTAGTTCCAAAAAACTGAGC<br>CTTGCTGAGATTGACGATCTTACATTCGAGCTTAAGCCCTTAGGTGCGATTGGATGGAAGTTGAAGAGCATCTT<br>GTATACCCAGAGTGGGGAGATCGAACGCGCAAAAAATTGTTACCAGAATGCTACTAAGACTGGAGATGATTCG<br>TATTGGGCCGCCTATGTTTTAGGAATCCACTTTTTTAAAGATGGACATTACGAAGAGGCATTAGATTGGTTCAGT<br>AAAACAGGCAACATTGTAGACACTATTGAAGCGCATGGATTAATCTTACGCTTGATCTGCTGCCTGCGTCTTGA<br>GTTGTATCAAAAGGGCTACGATTTGTCTAAGGAATGCATTGAGCTTTTACCTATGCGCATCGAGGGTTATCGTTT<br>AGGGGAGGTTTTCGCCTCGAAATTAGGGATGGAGAGCCTTGATTTTGACGGGAAGATCCACAACGATCCCTCC<br>CAATTGATTATTTCGCGTGCCGATACGAAACCGTGA | NdeI<br>XhoI |
| Translated sequence | MGSSHHHHHHSSGENLYFQSMDKIYVYEEHSEVFSYWVNEIPRNSTLVYFDQHLDLKLIENSKMKRISQFIKEGLCIDELKKDVPCREDGRYCYGIDDFLYAAIQE<br>GLFRKIIWVYPHMLGEKGFSELLWGLLSLIPNHGKEFISSFRNGKNSASVQLNNIELHVTSIDYLGEFIINEQVIVDIDLDYFYDPKTNDLSNDIDETLNVLGNLGIINQV<br>KTMTYSIKSGFLPESFRWLGEYIAGQMGREIVYSKVDKISPKVTMEKISSSKKLSLAEIDDLTFELKPLGAIGWKLKSILYTQSGEIERAKNCYQNATKTGDDSYWAA<br>YVLGIHFFKDGHYEEALDWFSKTGNIVDTIEAHGLILRLICCLRLELYQKGYDLSKECIELLPMRIEGYRLGEVFASKLGMESLDFDGKIHNDPSQLIISRADTKP | | | |
| **PbsB** | pRSF-Duet | ORF | ATGGCTAAGAACTCTGATGCCATGATTGTATTTCCACCTTTGACAGAAGCGCGTTTGTTTCCTTACCTTAGTCTT<br>CCTATGATCACATCCTTTTTACGCTCAAAGGGTATGTCCGTTTCGCAGATTGATCTTAACATCGAATTGTGCCAC<br>ACTCTTTTCTCTGAGGACTGTCTGAATGAATATGTCTCCATTAATGAAAACGGGTCTAAAGACTTAAAGTTAATTT<br>ACAAGGTAGAAATGGCGAAATACCTTTTTAAAGAGCAAAAACAGCTTTATAACAACGTATTCATTGAGAAACGCT<br>CGTCAGACTCCCTTAAATATGATGTCCGTCTGGTCCGTCAGGGTATTGAGCTTCTTCTTGTCAATTCATTCTTGA<br>AACTTGAAATTACGAGTTTGGAGGAGATTCTTGAATCTGTGATGGACTTTAGCTGGAAAAAAAGTGATATCGCTA<br>CAAAAGTATTATACGAGAATATTAAAGAAAAAATCCTGAACGATAAACCCAAGATTTTCGCACTGTCCATCGCGT<br>ATTACTCCCAGATCCTTCCCTCTTTATTGATTTGTAAGTGGATTCGTGAATTGTCTCCCAATACCCATATTATCCT<br>GGGGGGGCAGCAGATCATGATTCGCCAGAGCTCCTTCTTGAGTTTGAACGGATTAAACCAATTTGTCGACAGT<br>TTAGGTATTAGCGCAGGTGAGGAGACTTTATTTATGTTAAATCGTTACTTAAAAAATGATTGTCAGATCGTGGAA<br>GTTCCTGACATTGTGTGGCTGAACGAGAAGAACGTTGAAAATGTGCCATCGAAATCAGCCTACCGTATCACTGA<br>TGCCCTTCCCCCAGACTTTAGTGACCTTCCGTATAAGAACTACTTAGACGAGGAGGTACACATGTCACTTATCA<br>CTTGTGTCGGTTGTTATTGGGGTCGCTGTACTTTCTGCTCGTATGGAAATCGTTCGCGCAAGGAGAAGTCCTAC<br>CAACAAAAAACAGCGCGTCAAGTGGCGAACGAGTGTGAAGATATTATCAATAATTATGGGGTCAATCGTATCAA<br>CTTCATCGATGAAAATACAAATCTTCGTCTGGTTGTGAATGCTGTGAAGATCCTGGAAAGCCGCGGTTATGAGA<br>TCAAATTCTCGACCCGCAATCGCTTGGAGAACGTGCTTTTGGATGCCGGGTTCTGTTCCGAGTTATCCAATTTA<br>GGGTGTATTTTAATGTCGGTTGGTTATGAGACGAACTCTCAACGCATTCTGGACAGCCTTGACAAGGGTGTACA<br>ATCGTCGAATTATCAACAGATCATCGACAACTTGCACAACGCTAACATCACATTGCGTATGTCGATTATTGGTGG<br>GCTTCCAGGGGAAACTGAAGATGAGATTGAATGCAGTGAGGAGTTCCTTTTAAAGAATCAGGACAAAATCGGAA<br>TTGACGTTATGCAAATGCTGGTGTTAGAGCCCGGTACTTACATTTATGAAGACACTAATAACCCGGATATTCGC<br>ATTCAATCAAGTAAAAACCTTCGTGGCAACAAGTTGTTGAATTACGGCATGGGTCGCATGGGCGCGACTTTCCA<br>GTATAGTGATGGCATTACATTCGAAGAAAAACTTAATCGCTTTTTGCAATTACACAAAAATATCAATCCGCAAAA<br>GAACGACGAATTACCGCCGGATAAGTATAAAAAAGAAGGGCAAGATATCACCTCAAACACCTTATTGATTAACC<br>CATGGACTAAAATCATCAAATTGGACAAAACCTATATCATGGATTTTGTCTGGCAGCGCATTTTCTTAGTACCGG<br>AGTCATTAGAGGTCTTTGGAAATAGTCTTATTATGAGCAAGGCCGAGGACAAGAAGTACCTGGAATATTTTGTG<br>GAGAAGGGTGTAATTAACCTGAATTAA | NcoI<br>BamHI |
| Translated sequence | MAKNSDAMIVFPPLTEARLFPYLSLPMITSFLRSKGMSVSQIDLNIELCHTLFSEDCLNEYVSINENGSKDLKLIYKVEMAKYLFKEQKQLYNNVFIEKRSSDSLKYD<br>VRLVRQGIELLLVNSFLKLEITSLEEILESVMDFSWKKSDIATKVLYENIKEKILNDKPKIFALSIAYYSQILPSLLICKWIRELSPNTHIILGGQQIMIRQSSFLSLNGLNQ<br>FVDSLGISAGEETLFMLNRYLKNDCQIVEVPDIVWLNEKNVENVPSKSAYRITDALPPDFSDLPYKNYLDEEVHMSLITCVGCYWGRCTFCSYGNRSRKEKSYQQ<br>KTARQVANECEDIINNYGVNRINFIDENTNLRLVVNAVKILESRGYEIKFSTRNRLENVLLDAGFCSELSNLGCILMSVGYETNSQRILDSLDKGVQSSNYQQIIDNLH<br>NANITLRMSIIGGLPGETEDEIECSEEFLLKNQDKIGIDVMQMLVLEPGTYIYEDTNNPDIRIQSSKNLRGNKLLNYGMGRMGATFQYSDGITFEEKLNRFLQLHKNI<br>NPQKNDELPPDKYKKEGQDITSNTLLINPWTKIIKLDKTYIMDFVWQRIFLVPESLEVFGNSLIMSKAEDKKYLEYFVEKGVINLN | | | |

### Materials and methods

**Genome mining and sequence similarity network**

Sequence similarity network (SSN) of MNIOs was created using the EFI-EST toolkit[2] and using the PFAM family PF05114 as query accounting for ca. 13,500 entries as of September 2024. An alignment score of 55 was applied for network generation and the representative node (rep-node) 50 was visualized in Cytoscape v3.10. Uniprot IDs of the MNIO hits were extracted from the SSN and analyzed utilizing the RODEO toolkit.[3] Sequence alignment and sequence logo generation was conducted in the Geneious application using Muscle 5.1 parallel perturbed probcons (PPP) algorithm through 100 hidden Markov model (HMM) perturbations.[4]

**Plasmid constructs**

Codon-optimized (for *E. coli* K12) synthetic genes encoding precursor peptides were ordered as gBlocks™ from Integrated DNA Technologies (IDT) or adapter-removed gene fragments from Twist Bioscience with sequences homologous to the 5' and 3' ends of the plasmid vector insertion site. The gene fragments were cloned in pETDuet™-1 vector (Novagen) between the BamHI and EcoRI or HindIII restriction enzyme recognition sites (as stated in Table S7) through Gibson assembly for heterologous expression and production of N-terminally 6xHis-tagged precursor peptides. Similarly, genes encoding the modifying enzymes were ordered from TWIST Biosciences and were cloned into the pRSFDuet™-1 vector (Novagen) at NdeI and KpnI or XhoI sites (Table S7). For monocistronic expression of codon-optimized genes encoding PbsC, PbsD, PbsE and PbsE (in order), between the NdeI and KpnI sites of pRSF-Duet vector, with the ribosome binding sequence (RBS) 5'-GCGAAGGAGATATACCAG-3' in between each gene leading to the construct pRSF-PbsBCDE. For the PbsB- and PbsE-encoding plasmid, *pbsB* was inserted between the NcoI and BamHI sites, while *pbsE* was cloned between the NdeI and XhoI sites of the same pRSF-Duet vector, leading to the assembly of the pRSF-PbsBE construct. Similarly, the PbsC and PbsD encoding plasmid (pRSF-PbsCD) was created by inserting *pbsC* gene between the NdeI-XhoI sites and *pbsD* between the NcoI-EcoRI restriction sites of pRSF-Duet vector. For co-expression studies involving individual enzymes, pRSF-PbsB pRSF-PbsC, pRSF-PbsD and pRSF-PbsE constructs containing each gene in a different vector were cloned into the NdeI and KpnI restriction sites of pRSF-Duet vector. *E. coli* DH5α was used as the cloning strain for propagation of the vectors.

**Heterologous expression and purification of peptides**

Bacterial growth and protein production was conducted in a modified Terrific Broth according to a previous report[5] (TB; constituting yeast extract 24 g/L, tryptone 20 g/L, 17 mM $KH_2PO_4$, and 72 mM $K_2HPO_4$) supplemented with 5% glycerol, appropriate antibiotics and 1x final concentration of trace metal mix (Teknova). *E. coli* BL21(DE3) Tuner was used for protein production. An overnight starter culture of the appropriate genetically modified organism was used to inoculate a 100 mL subculture (for optimization experiments) or 1 L subculture in 2 L flasks (for larger scale production) as 1% inoculum and grown at 37 ºC until the optical density at 600 nm ($OD_{600}$) reached ~0.8. The cells were then transferred to ice for 15 minutes followed by addition of 0.5 mM IPTG to induce gene expression, 1 mM freshly prepared ferrous citrate (0.1 mg $FeSO_4$ in 1 mL $Na_3C_6H_5O_7$). For co-expression experiments involving PbsB, 0.2% (w/v, final) arabinose and 3 mM of hydroxocobalamine (OH-Cbl; Sigma) were added. Induced cells were further cultivated at 16 ºC for the next 36 hours at 120 rpm. Cells were then harvested by centrifugation at 6,000 × g at 4 ºC and re-suspended in 50 mM Tris-HCl buffer containing 300 mM NaCl, 10% glycerol, and 20 mM imidazole at pH 8.0 ($NPI_{20}$). DNase I (Sigma) and lysozyme (GoldBio) were added at 0.01 and 0.1 mg/mL, respectively. A final concentration of 1 mM $MgCl_2$ was added to support DNase activity and the cell mixture was stirred for 1 hour at 4 ºC. The cell suspension was sonicated at an amplitude of 40% for 20 cycles of 5 seconds on/off with a 6 mm probe. The lysate was centrifuged at 18000 × g at 4 ºC for 1 hour and Ni-NTA beads (MCLAB) were added to the supernatant and rotated 4 ºC for 1 hour. The pellet was further resuspended in denaturation buffer (50 mM Tris-HCl buffer containing 300 mM NaCl, 10% glycerol, and 6M guanidine hydrochloride at pH 8.0) followed by sonication and centrifugation as mentioned above. The denatured supernatant was collected and Ni-NTA beads were added. The mixtures were rotated at 4 ºC for 1 hour before loading on to a column. At this point, NI-NTA beads collected from both native and denaturing purification were loaded on the same column. The beads were washed 2 times with 3 × column volume (CV) of denaturing buffer, then 3 × CV of $NPI_{20}$ followed by 3 × CV of $NPI_{40}$. The desired protein was eluted twice with 2 × CV of $NPI_{750}$ (i.e., lysis buffer containing 750 mM imidazole). The above conditions remained true for all expression conditions unless stated otherwise.

**Isolation of matured C-terminal core peptide**

Following IMAC purification, the peptide eluent was desalted and buffer exchanged into elution buffer (50 mM Tris-HCl, 100 mM NaCl at pH 8.0) using a Sephadex PD10 column. To isolate the matured core peptide, proteolysis was conducted overnight using GluC endopeptidase (NEB) at 1:1000 enzyme:substrate ratio. Ni-NTA was added to the digest and rotated at 4 ºC for 1 hour before loading on to a column. The flowthrough containing the digested core fragment was collected and desalted by solid phase extraction (Chromabond® c18ec) and subsequently purified by High-performance liquid chromatography (HPLC).

**Isolation of modified peptide core fragments**

SPE column-eluted samples were lyophilized followed by resuspension in 5% acetonitrile containing 0.1% formic acid. Dissolved peptide sample was centrifuged at 12000 xg for 10 min and the supernatant was subjected to multiple rounds of HPLC purification. All purifications were conducted on a Vanquish UHPLC system (Thermo Fisher Scientific). In the first round of purification, a Phenomenex Aeris™ peptide C18-XB LC column (Part nr. 00G-4632-N0; particle size: 5 µm; dimensions 250 x 10 mm; pore size 100 Å) was used. Mobile phase used was solvent A containing water+0.1% formic acid and solvent B containing acetonitrile+0.1% formic acid. Separation was achieved over a 40 min method consisting of a 2 min equilibration segment at 5% solvent A, followed by a gradient of solvent B from 5 to 40% over 18 min. Following the gradient, the method contained a cleaning step at 95% solvent B over 10 min, followed by a re-equilibration step at 5% solvent A over the next 10 min. A flow rate of 2 ml/min was used throughout the method. Elution of the PbsA3-CDE was achieved at a retention time of 13.5 min, while the PbsA3-BCDE peptide eluted at 12.5 min. In the next purification cycle, the respective peptides were purified on an Accucore-C18 LC column from Thermo Fisher Scientific (Part nr: 17126-154630; particle size: 2.6 µm; dimensions: 150 x 4.6 mm; pore size: 80 Å). The same mobile phase was used as the first round. Separation was achieved over a 14 min method consisting of a 2 min equilibration segment at 5% solvent A, followed by a gradient of solvent B from 10 to 25% over 8 min. Following the gradient, the method contained a cleaning step at 95% solvent B over 5 min, followed by a re-equilibration step at 5% solvent A over the next 5 min. A flow rate of 1 ml/min was used throughout the method. PbsA3-BCDE eluted at 6.1 min while PbsA3-CDE eluted at 6.4 min. In case of PbsA3-YTY variants, all parameters remained same, except a flow rate of 1.2 and 1.5 ml/min were used for YTY-CD and YTY-B peptides, respectively in the second round of HPLC purification.

**High-resolution tandem mass spectrometry**

The desalted protease-digested peptides were injected on to an Agilent 1290 LC-MS QToF for ESI-HR-MS and MS/MS analysis. LC separation was conducted at 50 ºC on a 10%-80% gradient of Acetonitrile-water (+0.1% formic acid) over 11 mins at 0.6 ml/min flowrate on a Phenomenex Aeris 2.6 µm PEPTIDE XB-C18 LC column (part nr. 00F-4505-E0). Mass spectra was collected in positive mode at 10 spectra/s and 100 ms/spectrum. Tandem-MS Fragmentation was achieved at normalized collision energies of 20, 25 and 30. HR-MS/MS analysis was performed using the Interactive Peptide Spectral Annotator (IPSA) tool[6] and verified manually.

**Advanced Marfey's analysis**

The absolute stereochemistry of the hydroxylated Phe8 and Phe10 as well as the Orn9 residues was determined as previously described, with slight modifications.[7] Briefly, for the PbsA3-CD product, 100 µg of dried peptides was hydrolyzed in 0.8 mL of 6 M HCl in $H_2O$ in a 4 mL amber glass vial. In case of the PbsA3-CDE and PbsA3-BCDE products for the stereochemical analysis of the Orn9 residue, 6M DCl in $D_2O$ was used for hydrolysis. The mixture was heated at 120 °C for 16 h with stirring. The resulting product was vacuum-dried followed by resuspension in 1 mL of deionized water and dried again. The drying step was repeated to remove any remaining acid. Next, 0.6 mL of 0.8 M NaHCO3 (in $H_2O$) and 0.4 mL of a 3 mg/mL solution of Nα-(5-fluoro-2,4-dinitrophenyl)-l-leucinamide (L-FDLA; Sigma) in acetonitrile were added, followed by stirring in the dark (amber glass vial) at 67 °C for 8 hours. Post-derivatization, 100 µL of 6 M HCl was added to neutralize the reaction, and the mixture was vortexed. The mixture was lyophilized and subsequently resuspended in 200 µL of acetonitrile by sonication. The suspension was centrifuged at 12,000g for 10 min, and the supernatant was analyzed by LC–MS on an Agilent 6545 LC/Q-TOF instrument. Chromatographic separation was obtained on a Kinetex 1.7 µm F5 100 Å, LC column (100 × 2.1 mm; Phenomenex; part no.: 00D-4722-AN). A column oven was maintained at 45 °C, and the mobile phase used was A: water with 0.1% formic acid and B: acetonitrile with 0.1% formic acid. At a constant flow rate of 0.4 mL/min, a gradient of 2–10% B over 2.5 min, 10–80% B over the next 7.5 min was maintained, followed by a wash step of 95% B for 5 min and a post-run equilibration stage of 2% B for 5 min. MS spectra were collected in the negative ion polarity mode. 0.5–1 µL injections were performed for each sample. For co-injection experiments, the data collected from the individual runs was used to roughly normalize the amount of each test and standard samples and each test:standard sample was combined in 1:1 ratio, followed by LC-MS analysis.

**NMR data acquisition and analysis**

Sample information: Six peptides were studied by NMR in this work. The peptides PbsA3-CD (~ 0.6 mM) in 90% $H_2O$ and 10% $D_2O$ or in 100% $D_2O$, PbsA3-CDE (~ 1.6 mM) in 90% $H_2O$, 10% $D_2O$ and 0.1% formic acid-$d_2$ (dFA), PbsA3-BCDE (~ 0.7 mM) in 90% $H_2O$, 10% $D_2O$ and 0.1% dFA, YTY-B in 90% $H_2O$, 10% $D_2O$ and 0.1% dFA (~ 1.4 mM), the mixture of YTY-CD 1-OH (~ 0.8 mM) and YTY-CD 2-OH (~ 1 mM) in 90% $H_2O$ and 10% $D_2O$ were dissolved in ~ 550 µL volume and transferred into a 5-mm Wilmad 535-pp NMR tube. NMR data were collected at 25 °C on a Bruker Avance NEO 600 MHz spectrometer equipped with a 5-mm BBO prodigy probe or an Agilent VNMRS 750 MHz NMR spectrometer equipped with room temperature indirect detection 5-mm HCN probe especially for experiments collected at lower temperature of 3 °C.

One-dimensional (1D) $^{1}H$ NMR, two-dimensional (2D) homonuclear $^{1}H$-$^{1}H$ COSY (correlation spectroscopy which reveals the correlation between two neighboring protons), $^{1}H$-$^{1}H$ TOCSY (total correlation spectroscopy which reveals the correlation of protons in the same spin system) at 80 ms mixing time, $^{1}H$-$^{1}H$ NOESY (Nuclear Overhauser Effect spectroscopy which reveals close proximity in space between two protons) at 400 ms mixing time, $^{1}H$-$^{13}C$ HSQC (heteronuclear single quantum coherence spectroscopy, revealing one-bond correlation between $^{1}H$ and $^{13}C$), and $^{1}H$-$^{13}C$ HMBC (heteronuclear multiple bond correlation, revealing long range $^{1}H$-$^{13}C$ connectivity such as 2-bond, 3-bond, or 4 or more bonds).

Spectra were acquired for each sample using either pulse programs in Bruker Topspin 4.1.4 or the Biopack pulse sequences in the VNMRJ 4.2A software. The spectra were processed and analyzed in Mnova (version 15.1.0.; Mestrelab Research). The sample concentrations were determined by $^{1}H$ spectra collected under the qNMR condition on either the Bruker Avance NEO 600 MHz spectrometer compared with a standard sample (ERETIC 2 method) or an Agilent VNMRS 750 MHz spectrometer with a HCN probe that was calibrated with a known standard. The calibrated parameters were saved to the probe file and used to calculate the sample concentration based on the integration values of the proton peaks.

**Purification of PbsA3 truncants for minimal substrate analysis of PbsC**

Unmodified full length PbsA3 purified by IMAC was incompletely digested using endoproteinase LysC (New England Biology) at a substrate to enzyme ratio of 10000:1 for 15 mins, followed by immediate quenching with 0.25% formic acid (final concentration). The reaction was then subjected to HPLC separation leading to the purification of the 31-mer C-terminal fragment containing the conserved region of the leader peptide (Figure 4C) and the 20-mer fragment without the leader region. For HPLC, a Phenomenex Aeris™ peptide C18-XB LC column (Part nr. 00G-4632-N0; particle size: 5 µm; dimensions 250 x 10 mm; pore size 100 Å). Mobile phase used was solvent A containing water+0.1% formic acid and solvent B containing acetonitrile+0.1% formic acid. Separation was achieved over a 40 min method consisting of a 4 min equilibration segment at 5% solvent A, followed by a gradient of solvent B from 20 to 35% over 16 min. Following the gradient, the method contained a cleaning step at 95% solvent B over 10 min, followed by a re-equilibration step at 5% solvent A over the next 10 min. A flow rate of 2 ml/min was used throughout the method.

**Heterologous expression and purification of PbsCD and PbsE**

N-terminally His-tagged *pbsC* gene was inserted between the NdeI-XhoI restriction sites of pRSF-Duet vector along with a Tobacco Etch virus (TEV) protease cleavage site (ENLYFQS translated sequence) between the 6xHis tag and the *pbsC* gene. The untagged *pbsD* gene was inserted between the NcoI-EcoRI sites of the same vector, thus forming the pRSF-6xHis-PbsCD construct. For PbsE purification, the N-terminally His-tagged *pbsE* gene was cloned between the NdeI-XhoI sites of a pCDF-Duet vector, also with a TEV protease cleavage site after the His-tag to construct the pCDF--6xHis-PbsE plasmid.

*E. coli* BL21(DE3) Tuner was used for protein production. An overnight starter culture in LB media was used to inoculate a 1 L culture (1% inoculum) of TB with the appropriate antibiotic and grown at 37 ºC until the optical density at 600 nm ($OD_{600}$) reached ~0.8. For PbsE, 1x trace metal mix (Teknova) was included in the media. The cells were then transferred to ice for 30 minutes followed by addition of 0.7 mM IPTG. For PbsCD, 360 μM iron(II) sulfate, and 1 mM sodium citrate were added at the time of induction. Induced cells were further cultivated at 18 ºC for 16-18 hours. Cells were then harvested by centrifugation at 6,000 × g at 4 ºC and frozen at -80 ºC. For purification, cells were resuspended in 25 mM HEPES, 300 mM NaCl, 30 mM imidazole, pH 7.6. DNAse and lysozyme were added at 0.01 and 0.1 mg/mL, respectively, in addition to a EDTA-free protease inhibitor tablet (Pierce), and the cell mixture was stirred for 15 minutes at 4 ºC. The cell suspension was sonicated at an amplitude of 30% for 80 cycles of 10 seconds on/off with a 6 mm probe. The lysate was centrifuged at 18000 × g at 4 ºC for 1 hour and the supernatant was loaded onto a 5 mL HisTrap Ni-NTA column (Cytiva) on an AKTA Start purification system. The column was washed with 10 × column volume (CV) of lysis buffer, then eluted in 25 mM HEPES, 100 mM NaCl, 300 mM imidazole, pH 7.6. Fractions containing protein were purified using size exclusion chromatography directly, by injecting the eluate onto an equilibrated HiLoad 16/60 Superdex 200 pg column (Cytiva) in storage buffer (25 mM HEPES, 300 mM NaCl, pH 7.6). Glycerol was added to 10% final prior to flash freezing and storage in -80 ºC.

For PbsE purification, gravity-based IMAC purification was conducted. Briefly, cells were resuspended in 25 mM HEPES, 300 mM NaCl, 30 mM imidazole, pH 8. DNAse I and lysozyme were added at 0.01 and 0.1 mg/mL, respectively. $MgCl_2$ was added at 1 mM final concentration to support DNase I activity. The cell mixture was stirred for 30 minutes at 4 ºC. The cell suspension was sonicated at an amplitude of 40% for 40 cycles of 5 seconds on/off with a 6 mm probe. The lysate was centrifuged at 18000 × g at 4 ºC for 1 hour and the supernatant was incubated with the Ni-NTA beads for 1 hour at 4 ºC. The beads were loaded onto a column and washed 3 times with 3 × column volume (CV) of $NPI_{20}$ followed by 3 × CV of $NPI_{40}$. The desired protein was eluted twice with 2 × CV of $NPI_{750}$. Eluted protein was buffer-exchanged in a PD10 column into 25 mM

HEPES containing 100 mM NaCl at pH 8.0 and stored in 10% glycerol (final) at -80 ºC until further use.

***In vitro* assays**

For *in vitro* assays, PbsCD (25 μM) was mixed with HPLC-purified PbsA3 full length or truncated peptide (50 μM) in reaction buffer (25 mM HEPES, 200 mM NaCl, pH 7.6). For reactions containing iron(II) sulfate or sodium ascorbate, these additives were freshly dissolved in buffer and added to the reaction at a final concentration of 1 mM. The reactions were incubated at room temperature overnight. Reactions were then desalted with a C18 ZipTip (Millipore) and analyzed using MALDI-TOF or LC-MS/MS. In case of PbsE, 25 μM of the purified full length unmodified PbsA3, or the PbsA3-CD product (obtained from co-expression of PbsA3 with PbsCD, purified as a mixture of unmodified, mono-hydroxylated and bis-hydroxylated PbsA3) were added to reactions containing 25 μM of PbsE and 1 mM MnSO4. The reactions were incubated at 37 ºC for 2 hours followed by desalting with a C18 ZipTip and analysis using MALDI-TOF.

#### References


(1) Brademan, D. R.; Riley, N. M.; Kwiecien, N. W.; Coon, J. J. *Mol. Cell. Proteom.* **2019**, *18*, S193.

(2) Oberg, N.; Zallot, R.; Gerlt, J. A. *J. Mol. Biol.* **2023**, *435*, 168018.

(3) Tietz, J. I.; Schwalen, C. J.; Patel, P. S.; Maxson, T.; Blair, P. M.; Tai, H. C.; Zakai, U. I.; Mitchell, D. A. *Nat. Chem. Biol.* **2017**, *13*, 470.

(4) Edgar, R. C. *Nat. Commun.* **2022**, *13*, 6968.

(5) Padhi, C.; Field, C. M.; Forneris, C. C.; Olszewski, D.; Fraley, A. E.; Sandu, I.; Scott, T. A.; Farnung, J.; Ruscheweyh, H.-J.; Narayan Panda, A.; Oxenius, A.; Greber, U. F.; Bode, J. W.; Sunagawa, S.; Raina, V.; Suar, M.; Piel, J. *Proc. Natl. Acad. Sci. USA* **2024**, *121*, e2409026121.

(6) Brademan, D. R.; Riley, N. M.; Kwiecien, N. W.; Coon, J. J. *Mol. Cell. Proteomics* **2019**, *18*, S193.

(7) Eslami, S. M.; Padhi, C.; Rahman, I. R.; van der Donk, W. A. *ACS Synth. Biol.* **2024**, *13*, 2128.
